# hPSC-Derived Enteric Ganglioids Model Human ENS Development and Function

**DOI:** 10.1101/2022.01.04.474746

**Authors:** Homa Majd, Ryan M Samuel, Jonathan T Ramirez, Ali Kalantari, Kevin Barber, Zaniar Ghazizadeh, Angeline K Chemel, Andrius Cesiulis, Mikayla N Richter, Subhamoy Das, Matthew G Keefe, Jeffrey Wang, Rahul K Shiv, Conor J McCann, Samyukta Bhat, Matvei Khoroshkin, Johnny Yu, Tomasz J Nowakowski, Hani Goodarzi, Nikhil Thapar, Julia A Kaltschmidt, Faranak Fattahi

**Affiliations:** Department of Cellular and Molecular Pharmacology, University of California, San Francisco, San Francisco, CA 94158, USA; Eli and Edythe Broad Center of Regeneration Medicine and Stem Cell Research, University of California, San Francisco, San Francisco, CA, 94143, USA; Division of Cardiovascular Medicine, Stanford University School of Medicine, Stanford, CA, USA; Department of Neurosurgery, Stanford University School of Medicine, Stanford, CA 94305, USA; Department of Anatomy, University of California, San Francisco, San Francisco, CA, USA; Department of Psychiatry and Behavioral Sciences, University of California, San Francisco, San Francisco, CA, USA; Department of Biochemistry and Biophysics, University of California, San Francisco, San Francisco, CA 94158, USA; Department of Electrical engineering, Stanford University, Stanford, CA 94305, USA; Stem Cells and Regenerative Medicine, UCL Great Ormond Street Institute of Child Health, London, UK; NIHR Great Ormond Street Hospital BRC, London, UK; Department of Neurological Surgery, University of California, San Francisco, San Francisco, CA, USA; Department of Urology, University of California, San Francisco, San Francisco, CA 94158, USA; Gastroenterology, Hepatology and Liver Transplant, Queensland Children’s Hospital, Brisbane, Queensland, Australia; School of Medicine, University of Queensland, Brisbane, Australia; Woolworths Centre for Child Nutrition Research, Queensland University of Technology, Brisbane, Australia; Wu Tsai Neurosciences Institute, Stanford University, Stanford, CA 94305, USA; Program in Craniofacial Biology, University of California, San Francisco, San Francisco, CA 94110, USA

## Abstract

The enteric nervous system (ENS) plays a central role in gut physiology and mediating the crosstalk between the gastrointestinal (GI) tract and other organs. The human ENS has remained elusive, highlighting the need for an *in vitro* modeling and mapping blueprint. Here we map out the developmental and functional features of the human ENS, by establishing robust and scalable 2D ENS cultures and 3D enteric ganglioids from human pluripotent stem cells (hPSCs). These models recapitulate the remarkable neuronal and glial diversity found in primary tissue and enable comprehensive molecular analyses that uncover functional and developmental relationships within these lineages. As a salient example of the power of this system, we performed in-depth characterization of enteric nitrergic neurons (NO neurons) which are implicated in a wide range of GI motility disorders. We conducted an unbiased screen and identified drug candidates that modulate the activity of NO neurons and demonstrated their potential in promoting motility in mouse colonic tissue *ex vivo*. We established a high-throughput strategy to define the developmental programs involved in NO neuron specification and discovered that PDGFR inhibition boosts the induction of NO neurons in enteric ganglioids. Transplantation of these ganglioids in the colon of NO neuron-deficient mice results in extensive tissue engraftment, providing a xenograft model for the study of human ENS *in vivo* and the development of cell-based therapies for neurodegenerative GI disorders. These studies provide a framework for deciphering fundamental features of the human ENS and designing effective strategies to treat enteric neuropathies.

## Introduction

The enteric nervous system (ENS) is the largest and most complex division of the autonomic nervous system (De Giorgio, 2006). More than 500 million enteric neurons and roughly seven times as many enteric glia form interconnected enteric ganglia embedded in two distinct layers within the gut wall: the myenteric plexus residing between the longitudinal and circular muscles, and the submucosal plexus residing between the circular muscle and the mucosa (Grubišić and Gulbransen, 2017; Grundmann et al., 2019; Hamnett et al., 2021; Sasselli et al., 2012).

The ENS, uniquely, is not dependent on input from the central nervous system (CNS) to command GI tract functions (Furness et al., 2014). This autonomy is exemplified by studies in which segments of the bowel removed from the body continue to generate complex motor patterns *ex vivo*. ENS autonomy is the result of extraordinarily diverse neuronal and glial cell types with distinct neurochemical signatures working together in harmony (Brehmer, 2021; Qu et al., 2008; Fung and Vanden Berghe, 2020). Thus, the ENS is equipped to control complex gut functions including motility, secretion, absorption, blood flow regulation and barrier function support. Furthermore, the ENS communicates extrinsically with the CNS, enteroendocrine system, immune system, and the gut microbiome in order to maintain vitality and proper gut homeostasis (Furness et al., 2014; Long-Smith et al., 2020; Muller et al., 2014; Obata and Pachnis, 2016; Schneider et al., 2019; Yoo and Mazmanian, 2017).

The neurochemical and functional complexity of the ENS resemble the CNS (Gershon, 1999), yet much slower progress has been made in the field of ENS research. Despite being the largest and most complex division of the peripheral nervous system and playing a central role in the development and progression of enteric neuropathies and diseases of the gut-brain axis, ENS research has been disproportionately affected by multiple longstanding technical challenges. For example, enteric neurons are diluted throughout the GI tract, comprising less than 1% of gut tissue (Drokhlyansky et al., 2020). Therefore, scientists must rely on samples collected during GI resection surgeries, rather than more routine GI biopsies, to access ENS tissue. Furthermore, it is difficult to isolate the ENS without significant sampling bias related to harsh tissue dissociation techniques that damage fragile neurites, and reliable surface markers suitable for FACS purification of enteric neurons and glia are lacking.

The complex developmental processes and the elaborate cellular architecture of the ENS, as well as its remarkable communication with the rest of the body provide a wide array of possibilities for abnormalities to arise. Comprising some of the most challenging clinical disorders, enteric neuropathies, also known as disorders of gut brain interaction (DGBI), result from loss, degeneration or functional impairment of the ENS cell types (De Giorgio et al., 2016; Niesler et al., 2021). Our incomplete understanding of ENS development and function is accountable for the long-term morbidity and mortality of GI disorders and limited availability of therapeutic interventions. Specifically, there has been an extraordinary interest in gaining a better understanding of enteric nitrergic neurons (NO neurons) that release nitric oxide (NO) to relax the smooth muscle tissue and promote GI motility. This is due to the selective dysfunction, and degeneration of NO neurons in different forms of DGBI including, esophageal achalasia, infantile hypertrophic pyloric stenosis and gastroparesis (Bódi et al., 2019; Rivera et al., 2011).

Here, we describe an experimental system for deriving ENS tissue from human pluripotent stem cells (hPSCs) that recapitulate the remarkable cellular diversity of the human ENS. These three-dimensional (3D) cultures, termed enteric ganglioids, along with two-dimensional (2D) ENS cultures provide scalable sources of human enteric neurons and glia, that are compatible with a wide array of high-throughput applications. We use single cell transcriptomics to map cell-type specific molecular features of human enteric neurons and glia which offers new strategies for their enrichment, isolation, or functional targeting. We leverage hPSC-derived enteric ganglioids as a model system to investigate the development of NO neurons, characterize their molecular and physiological properties and identify clinically relevant strategies to modulate their function *in vitro* and in the mouse colon *ex vivo*. Further, we demonstrate the extensive engraftment and regenerative potential of NO neuron ganglioids in the colon of adult mice, providing a new xenograft model to study the human ENS *in vivo*.

## Results

### Derivation of enteric ganglioids from hPSCs to model development, function and molecular diversity of the human ENS

The ENS is derived from the vagal and sacral neural crest (NC). Vagal NC cells extensively migrate and colonize the entire length of the GI tract, whereas sacral NC cells only colonize the most distal end of the colon (Serbedzija et al., 1991; Burns and Douarin, 1998; Heanue and Pachnis, 2007; Nagy and Goldstein, 2017). We have previously established hPSC differentiation methods to derive enteric neural crest cells (ENCs) under highly defined conditions (**Figure 1A**) (Barber et al., 2019; Fattahi et al., 2016). This protocol involves two steps that follow embryonic NC development. In step 1, we induce enteric neural crest by activating bone morphogenic protein (BMP) and Wnt signaling in combination with retinoic acid (RA) treatment. RA caudalizes the differentiating NC, specifying a vagal NC identity. In step 2, we generate enteric crestospheres in the presence of Wnt and fibroblast growth factor (FGF) signaling (Barber et al., 2019; Fattahi et al., 2016).

**Figure 1.**
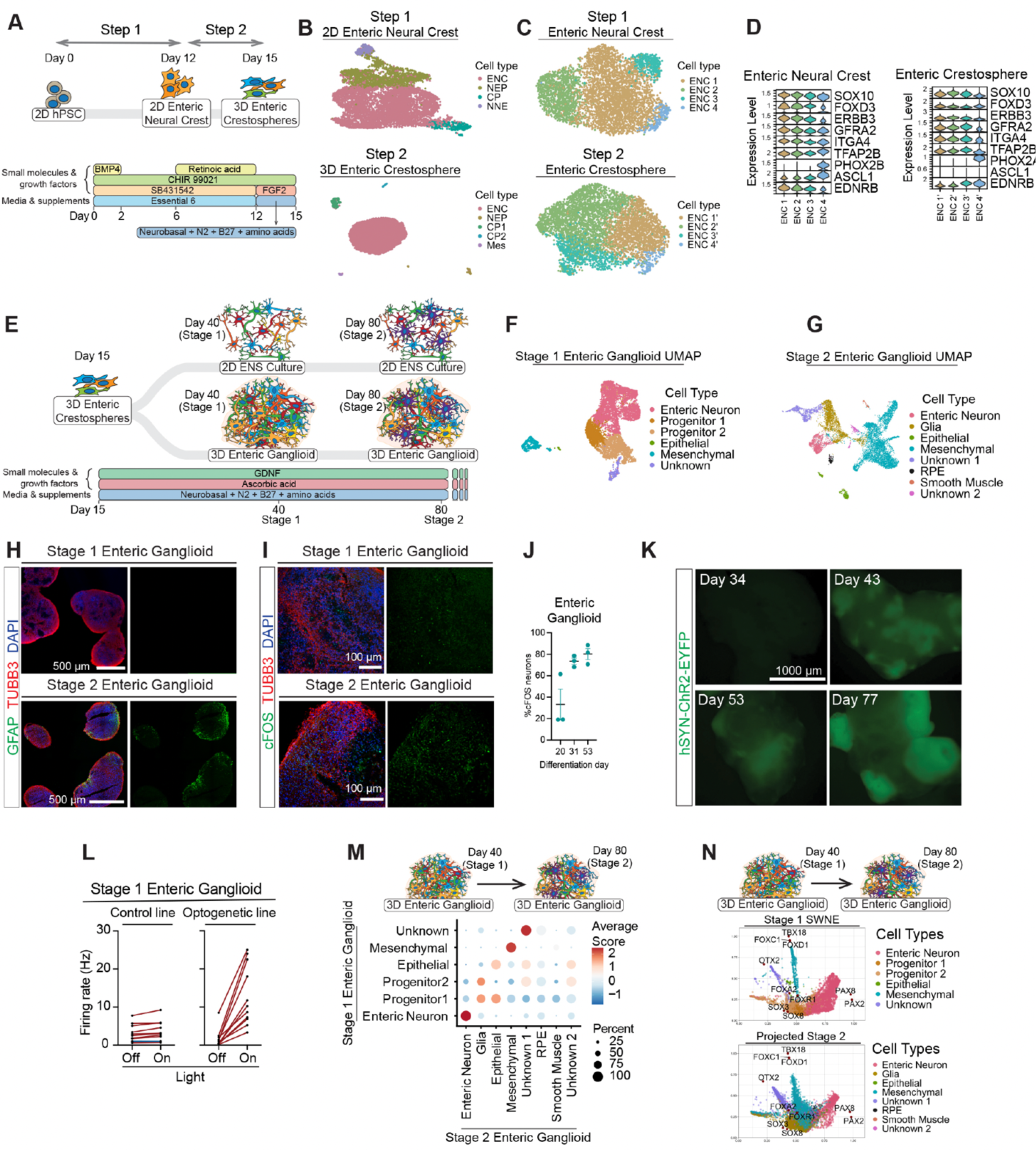
hPSC-derived enteric ganglioids model development, function and molecular diversity of human ENS. **A)** Protocol schematic for *in vitro* differentiation and maturation of hPSCs into enteric neural crest and enteric crestospheres. **B)** scRNA-seq UMAP of cell types present in enteric neural crest cells (D10, top panel) and enteric crestosphere cells (D15, bottom panel) of the differentiation cultures depicted in (**A**). **C)** UMAP of enteric neural crest (D10, top) and enteric crestosphere (D15, bottom) subtypes in differentiation cultures. **D)** Violin plot stack showing the expression of canonical enteric neural crest markers in enteric neural crest (top) and enteric crestosphere (bottom) subtypes. **E)** Protocol schematic for *in vitro* differentiation and maturation of hPSC-derived enteric crestospheres into 2D ENS cultures and 3D ganglioids. **F)** snRNA-seq UMAP of cell types present in stage 1 enteric ganglioids. **G)** snRNA-seq UMAP of cell types present in stage 2 enteric ganglioids. **H)** Immunofluorescence analysis for expression of neuronal TUBB3 and glial GFAP in stage 1 and stage 2 enteric ganglioids. **I)** Immunofluorescence analysis for expression of neuronal activity marker cFOS in stage 1 and stage 2 enteric ganglioids. **J)** Flow cytometry quantification of neuronal activity marker cFOS in enteric ganglioids as they mature. **K)** Live fluorescence images of human hSYN-ChR2-EYFP in enteric ganglioids as they mature. **L)** Quantification of multi-electrode array (MEA) analysis of baseline and blue light-stimulated neuronal activity in stage 1 hSYN-ChR2-EYFP (left) and control (right) enteric ganglioids. **M)** Dot plot of the average module scores of stage 1 enteric ganglioid cell type transcriptional signatures in stage 2 enteric ganglioid cell types. **N)** Projection of stage 2 cell types (right) onto the SWNE of stage 1 enteric ganglioid cells with overlayed projection of stage 1 cell-type specific transcription factors from **Figure S3**.

To characterize these developmental processes in the human ENS lineages at the molecular level, we performed single cell RNA-seq (scRNA-seq) on enteric neural crest and enteric crestospheres. During the enteric neural crest stage, four transcriptionally distinct cell types are present: enteric neural crest (ENC) (SOX10**^+^**, FOXD3**^+^**), neuro-epithelial progenitor (NEP) (WNT2B**^+^**, PAX6**^+^**), cranial placode (CP) (SIX1**^+^**, EYA2**^+^**) and non-neural ectoderm (NNE) (EPCAM**^+^**, CDH1**^+^**) (**Figure 1B top, Figure S1A left**). In the next step, suspension culture of enteric neural crest cells serves as a purification strategy that leads to enteric crestospheres consisting primarily of ENCs with a small population of NEPs, two CP clusters (CP1 and CP2), and a mesenchymal (Mes) (TWIST1**^+^**, MSX1**^+^**) cluster (**Figure 1B** **bottom, Figure S1A right**). Module scoring the transcriptional signature of cell types in step 1 and step 2 verifies the shared transcriptional identity of the ENC clusters (**Figure S1B**).

Further subclustering of the ENC populations identified four subtypes (ENC 1-4 for the enteric neural crest stage and ENC 1’-4’ for the enteric crestospheres) that differentially express canonical ENC markers such as SOX10, EDNRB, TFAP2B and FOXD3, and are chronologically transitioning from PHOX2B to PHOX2A expression (**Figure 1C and D**). To study how ENCs progress during the differentiation steps, we module scored ENC 1-4 transcriptional signatures in the enteric crestosphere ENC 1’-4’ (**Figure S1C**). ENC 1’ and 2’ showed high transcriptional similarity to ENC 4, and ENC 2 and 3, respectively, the three most transcriptionally distinct ENC subtypes (**Figure S1C, Figure S1D and E**).

hPSC-derived ENCs are previously shown to recapitulate key migratory features of ENS precursors in health and disease and are capable of giving rise to enteric neurons upon further differentiation (Barber et al., 2019; Fattahi et al., 2016) but their ability to generate the diverse array of neuronal and glial subtypes that comprise the human ENS has not been characterized. To determine the potential of enteric crestospheres to differentiate into ENS cell types, we established 2D and 3D culture conditions that facilitate the transition of ENCs into mature ENS cell types (**Figure 1E**). While 2D culture offers unique technical advantages for applications such as high content imaging assays, we chose to focus primarily on 3D cultures, termed enteric ganglioids, given their scalability and potential for capturing higher order cell-cell interactions that occur in the developing and adult ENS tissue. In addition, 3D culture platforms are technically advantageous in applications such as cell therapy.

To define the cellular composition of enteric ganglioids we performed single nuclei RNA-seq (snRNA-seq) on stage 1 (differentiation day 35-50) and stage 2 (differentiation day 70-90) enteric ganglioids (**Figure 1E**). Unbiased clustering of stage 1 enteric ganglioids revealed a large population of enteric neurons, two progenitor populations and small populations of contaminating epithelial cells, mesenchymal cells, and one cluster of unknown identity (**Figure 1F, Figure S1F**). Module scoring based lineage analysis revealed that the progenitor 1 population shared high transcriptional similarity to ENC 2’ and 4’. Furthermore, the mesenchymal population was highly similar to ENC 3’ (**Figure S1G**). Importantly, stage 2 enteric ganglioids contained enteric glia in addition to enteric neurons, indicating that gliogenesis follows neurogenesis during *in vitro* differentiation, which is consistent with the *in vivo* developmental timeline (**Figure 1G**) (Rothman et al., 1986; Young et al., 2003). We confirmed the presence of glia in our stage 2 enteric ganglioids by immunostaining for GFAP (**Figure 1H**). Stage 2 ganglioids contained a larger proportion of contaminating epithelial cells, mesenchymal cells, and two unknown clusters (**Figure 1G, Figure S1H**). At stage 2 epithelial and mesenchymal populations showed higher transcriptional diversity compared to stage 1 clusters and could be further sub-clustered into two unique epithelial populations and five unique mesenchymal populations (**Figure S1I and J**). Additionally, at this later stage, contaminating retinal pigmented epithelium (RPE) and smooth muscle cell populations emerged (**Figure 1G, Figure S1H**). To evaluate the functional maturation state of the enteric neurons over the course of differentiation, we evaluated the expression of the neuronal activity marker cFOS. Neuronal depolarization leads to expression of cFOS, a proto-oncogene that has been used as a marker for neuronal activity (Hunt et al., 1987; Bullitt, 1990; Santos et al., 2018). cFos expression increased as the enteric ganglioids progressed during differentiation (**Figure 1I and J**). To demonstrate synaptic maturation and electrical excitability of enteric neurons within gangliods, we used optogenetics by differentiating a reporter hESC line that expresses enhanced yellow fluorescent protein (EYFP) tagged channelrhodopsin-2 under control of the human synapsin promoter (Steinbeck et al., 2016). EYFP was readily detectable as early as day 43 (**Figure 1K**). Light stimulation of stage 1 ganglioids increased electrical firing rates, as detected by microelectrode array (MEA) (**Figure 1L, Figure S2A and B**), leading to increased cFOS expression as compared to unstimulated enteric ganglioids (**Figure S2C**). Thus, stage 1 and stage 2 enteric neurons are functional and continue to gain maturity over time.

Next, we explored the transcriptional differences, lineage relationships, and functional properties of stage 1 and stage 2 enteric ganglioids. Many genes, including transcription factors, neurotransmitter receptors, neuropeptide receptors, cytokines and their receptors, secreted signaling ligands and their receptors, and surface markers were exclusively expressed (detected in >25% of cells in a single cluster) by each population in the stage 1 and stage 2 enteric ganglioids (**Figure S3A-P**). Other genes in these categories, while not exclusively expressed, showed differential expression between cell types (**Figure S4A-H, Figure S5A-H**). To determine the lineages shared between stage 1 and 2 enteric ganglioids, we conducted module scoring based lineage analysis. Transcriptional signatures were highly conserved between stage 1 and 2 enteric neurons and mesenchymal cells (**Figure 1M**). Interestingly, the glia population was most transcriptionally similar to the two progenitor populations from stage 1 (**Figure 1M**). We confirmed these transcriptional similarities between stage 1 and stage 2 enteric ganglioid cell types by generating similarity weighted non-negative embeddings (Wu et al., 2018) (SWNE) by projecting stage 2 ganglioid cells onto the stage 1 SWNE (**Figure 1N**). Stage 2 cell types mapped to similar SWNE space positions as the matched stage 1 cell types, suggesting similar expression patterns and lineage continuation (**Figure 1N**).

To compare the cellular diversity between our 2D ENS cultures and enteric ganglioids we performed scRNA-seq on 2D cultures in stage 2. Similar to the 3D ganglioids, clustering and annotation by expression of key marker genes revealed enteric neurons, glia, epithelial and mesenchymal cells, and two unknown populations (**Figure S6 A and B**). Projection of 2D cells into 3D ganglioid SWNE space showed conserved expression patterns between 2D and 3D enteric neurons, glia, mesenchymal cells and unknown cluster 1 (**Figure S6C**). These observations were confirmed by calculating the Spearman correlation of the expression of 3000 shared variably expressed genes (Seurat anchor features) between stage 2 ganglioid and 2D ENS culture cell types (**Figure S6D**). Importantly, 2D and 3D enteric neurons and glia showed high correlations of .83 and .76, respectively (**Figure S6D**). Additionally, the mesenchymal and unknown 1 clusters showed a high correlations (.82, and .80, respectively), while the epithelial clusters showed a modest correlation of .29 (**Figure S6D**). Interestingly, the 2D specific unknown cluster 2 showed a moderate correlation to the glia cluster (**Figure S6D**). Taken together, these data indicate that the enteric neurons and glia generated by the new 3D differentiation format are highly transcriptionally similar to their 2D counterparts, while the off-target/contaminating cell types may vary in composition and transcriptional identity between the two formats.

### hPSC-derived enteric ganglioids recapitulate the neuronal diversity of the human ENS

Recent characterization of human and mouse primary enteric neurons at single cell resolution have revealed many transcriptionally distinct clusters of enteric neurons (Drokhlyansky et al., 2020; Morarach et al., 2021). To characterize the diversity of our hPSC-derived enteric neurons, we sub-clustered the neuronal populations in stage 1 and 2 ganglioids. Eight transcriptionally distinct neuronal subtypes (EN 1-8) were identified in the stage 1 enteric neurons (**Figure 2A, Figure S7A and B**). Module scoring revealed that EN 1 and EN 6 showed higher similarities with enteric crestosphere ENCs potentially representing earlier stage neuronal populations (**Figure S7C**). For the stage 2 enteric neurons, sub-clustering analysis similarly revealed eight distinct subtypes of enteric neurons (EN 1’-8’) (**Figure 2B, Figure S7D and E**). Comprehensive analysis of functionally and technically relevant gene categories revealed transcription factors, neuropeptides and their receptors, neurotransmitter receptors, cytokines and their receptors, secreted signaling ligands and their receptors, and surface markers that were exclusively expressed by enteric neuronal subtypes in stage 1 and 2 (**Figure S8A-Q**). Many genes in these categories, while not exclusively expressed, showed differential expression between enteric neuron subtypes (**Figure S9A-J, Figure S10A-J**). This signifies the remarkable functional diversity of cell types in the differentiated ganglioids.

**Figure 2.**
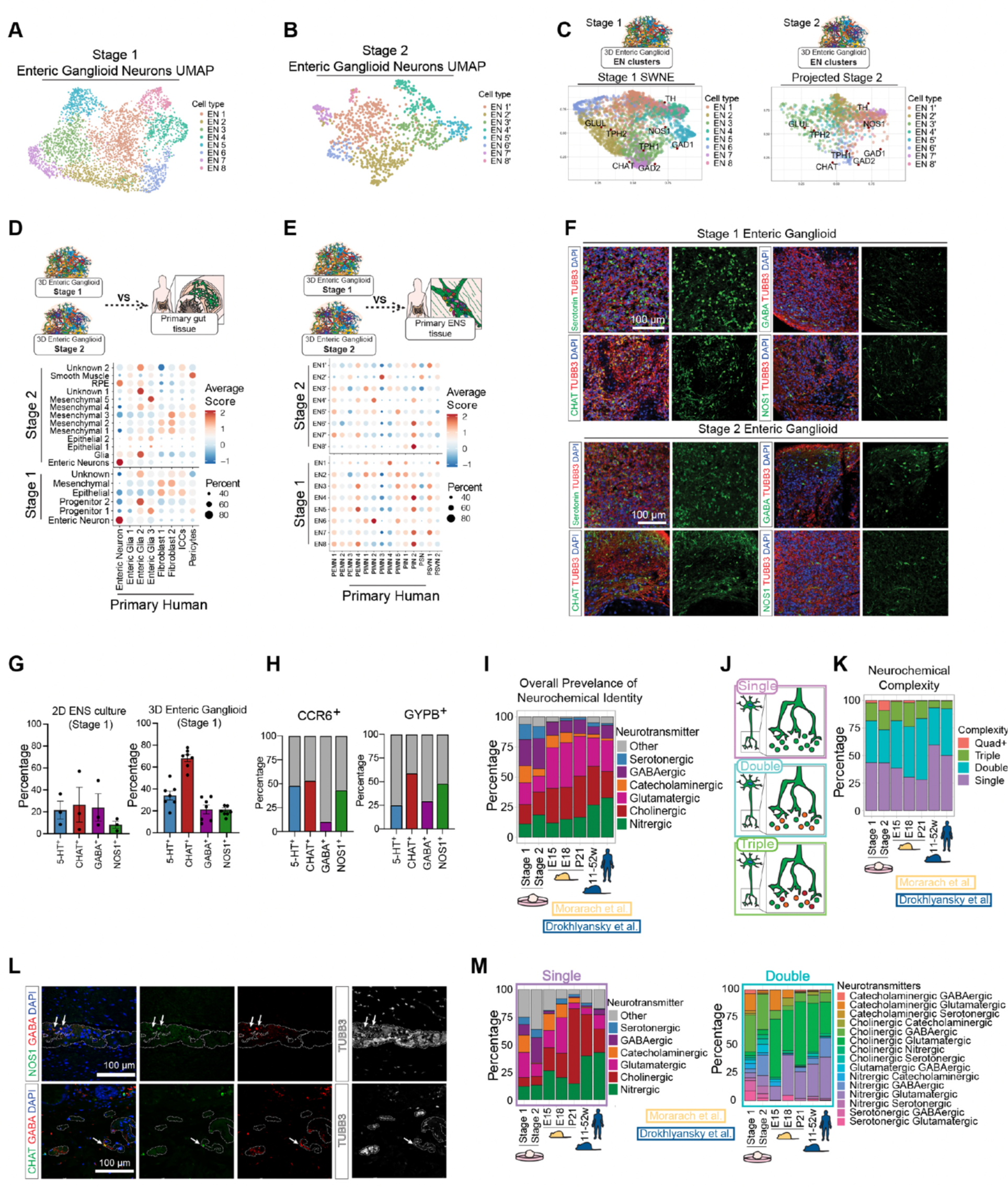
hPSC-derived enteric ganglioids model human ENS neurochemical diversity. **A)** snRNA-seq UMAP of neuronal subtypes present in stage 1 enteric ganglioids. **B)** snRNA-seq UMAP of neuronal subtypes present in stage 2 enteric ganglioids. **C)** Projection of stage 2 neuronal subtypes (right) onto the SWNE of stage 1 enteric ganglioid neurons with overlayed projection rate-limiting neurotransmitter synthesis enzymes. **D)** Dot plot of the average module scores of stage 1 (bottom) and stage 2 (top) ganglioid cell type transcriptional signatures adult human colon cell types. **E)** Dot plot of the average module scores of stage 1 (bottom) and stage 2 (top) ganglioid neuronal subtype transcriptional signatures adult human enteric neuron subtypes. **F)** Immunofluorescence analysis for expression of ENS cell-type markers (serotonin, CHAT, GABA and NOS1) in stage 1 enteric ganglioids. **G)** Quantification of flow cytometry analysis for the expression of neuronal subtype markers serotonin, CHAT, GABA and NOS1 in stage 1 2D ENS cultures (left) and 3D enteric ganglioids (right). **H)** Flow cytometry validation of stage 1 EN 8 surface markers CCR6 (left) and GYPB (right) co-labeling with neurochemical markers showing enrichment of neurochemical identities of marker positive populations normalized to baseline neurochemical population levels. **I)** Overall percentage of neurotransmitter synthesizing neurons in stage 1 and 2 enteric ganglioids compared to mouse and human primary enteric neurons. **J)** Schematic of mono- and multi-neurotransmitter synthesis in enteric neurons. **K)** Percentage of neurons showing mono-and multi-neurotransmitter profiles in stage 1 and 2 enteric ganglioid neurons compared to mouse and human primary enteric neurons. **L)** Immunostaining of primary human colon with antibodies against NOS1, GABA and TUBB3 (top), and CHAT, GABA and TUBB3 (bottom). White dash line indicates the border of TUBB3^+^ ganglia. White arrows indicate colocalization. **M)** Percentage of mono-neurotransmitter (top) and bi-neurotransmitter (bottom) producing enteric neurons in stage 1, 2 enteric ganglioids and primary datasets.

Next, we determined the lineage similarities between the stage 1 and 2 enteric neuron subtypes. Module scoring revealed the highest transcriptional similarity between EN 1 and EN 2’ (**Figure S11A**). Other stage 2 enteric neuron subtypes shared modest transcriptional similarities with multiple stage 1 enteric neuron subtypes (**Figure S11A**). To confirm this observation, we projected stage 2 enteric neurons into stage 1 enteric neuron SWNE space. In agreement, many stage 1 and 2 enteric neurons showed similar expression patterns based on regional overlap of EN 1 and EN2’, EN 2 and 6 with EN 3’, EN 4 and 8 with EN 1’ and 7’, and EN 7 with EN 6’ in the SWNE space (**Figure 2C**). Interestingly, very few stage 2 enteric neurons overlapped with stage 1 EN 3 and 5, suggesting these may be transient neuronal subtypes (**Figure 2C**).

To compare the neuronal diversity between the 2D cultures and ganglioids, we performed parallel sub-clustering analysis of the 2D stage 2 enteric neurons (**Figure S11B**). 2D enteric neurons clustered into five distinct enteric neuron populations (**Figure S11B and C**). Module scoring revealed that all ganglioid neuronal signatures are present in the 2D enteric neurons, however, EN 3’-5’ and EN 7’ and 8’ clustered together in the 2D enteric neuron dataset (**Figure S11D and E**). These results were further supported by Spearman correlation analysis of 3000 anchor features shared between 2D and ganglioid enteric neurons that revealed positive correlations between 2D EN 3’-5’ and ganglioid EN3’ and EN5’, as well as 2D EN 7’/8’ and ganglioid EN 7’ (**Figure S11F**).

To validate that hPSC-derived enteric ganglioids recapitulate *in vivo* ENS biology, we compared our stage 1 and 2 ganglioids to a snRNA-seq dataset of the primary human colon previously published by Aviv Regev and Colleagues (Drokhlyansky et al., 2020) (**Figure S11G**). Remarkably, module scoring of ganglioid cell type signatures on relevant primary cell types demonstrated that *in vitro* and *in vivo* enteric neurons were highly similar (**Figure 2D**). Likewise, correlating the expression of 3000 anchor features shared between the three datasets showed Spearman correlation values of .89 and .87 between the primary and the stage 1 and 2 enteric neurons, respectively (**Figure S11H top**). Further, module scoring and Spearman correlation analyses revealed that stage 1 and 2 enteric neuron subtypes represent transcriptional signatures of primary neuron subtypes from all neuron classes (**Figure 2E, Figure S11H bottom**). These data demonstrate that our ganglioids capture the diversity of neuronal transcriptional identities in the human ENS.

Another important component of an enteric neuron’s identity is its location in the myenteric or submucosal plexus. In order to generate myenteric and submucosal gene signatures, we utilized metadata associated with the human samples sequenced by Drokhlyansky et al. denoting the tissue layer from which each sample was collected. Module scoring the primary enteric neuron subtypes with these plexus gene modules found the tissue layer signatures to be mutually exclusive, with each neuronal subtype having a positive score for either one module or the other (**Figure S11I left**). Interestingly, this analysis suggests that PEMN, PIMN, PSN and PSVN categories contain both myenteric and submucosal subtypes. However, both PIN subtypes scored positively for the submucosal signatures (**Figure S11I left**). Module scoring our stage 1 and 2 enteric neurons suggests that our ganglioids generate both myenteric and submucosal neurons largely recapitulate the mutual exclusivity of these signatures (**Figure S11I middle and right**).

Enteric neuron identity is often described based on their neurochemical properties including nitrergic, cholinergic, glutamatergic, catecholaminergic, GABAergic or serotonergic. We first verified the presence of neurons with various neurochemical features in our stage 1 and stage 2 ganglioids by immunostaining (**Figure 2F**). These neurochemical markers were consistently represented in both 2D and ganglioid culture formats (**Figure 2G**). Intriguingly, our transcriptional analysis of both stages showed that multiple enteric neuron subtypes express the same neurotransmitter markers (**Figure S9B, Figure S10B**). For example, neurons in EN 4, 5 and 8 neurons expressed NOS1 which is the enzyme that produces NO and is a marker for nitrergic neurons. Additionally, individual enteric neuron subtypes expressed markers for multiple neurotransmitters. For example, EN 5 expressed NOS1 and GAD1 which is a gabaergic marker and EN 8 expressed the cholinergic marker SLC5A7 and the glutamatergic marker SLC17A6 (**Figure S9B**). We next performed flow cytometry to validate the co-expression of neurotransmitter markers within neuronal subclusters. As a proof of principle, we used two surface markers, CCR6 and GYPB, which specifically expressed in stage 1 EN 8 (**Figure S9J**), to label this subcluster and quantify the proportion of serotonin**^+^**, CHAT**^+^**, GABA**^+^**, and NOS1**^+^** enteric neurons. All four neurotransmitter markers were detected in CCR6**^+^** and GYPB**^+^** populations confirming that transcriptionally distinct enteric neuron subclusters are not defined by a single neurotransmitter identity (**Figure 2H**). This suggests that a single neurotransmitter cannot serve as a specific marker for annotation of transcriptionally distinct enteric neuron subtypes, and reaffirms the hypothesis that each neuron can take on multiple neurochemical identities.

To further explore this hypothesis at the single cell resolution, we designed a stringent two-step approach to define an enteric neuron’s neurochemical identity. In the first step, we identified neurons that expressed the hallmark rate limiting neurotransmitter synthesis enzymes (**Figure S12A and B**). In the second step, we module scored these neurons based of their expression of a curated list of neurotransmitter metabolism enzymes and transport proteins (**Table S1, Figure S12C and D**). Neurons that passed both steps were binned into a particular class of neurotransmitter identity (**Figure S12E and F**). For example, neurons were annotated as nitrergic if they expressed NOS1 and scored highly for NO metabolism and transport genes NOS1AP, ARG1/2, ASL and ASS1. We found that all EN subtypes, both in 2D and 3D, contained neurons from every neurotransmitter identity class (**Figure S12G-I**). Furthermore, at the single cell level, many neurons were equipped to synthesize multiple neurotransmitters (**Figure S12A-F**).

A defining aspect of an enteric neuron’s function is its ability to sense and respond to specific neurotransmitters released by other neurons. As a first step towards mapping the neuronal communication networks in enteric ganglioids, we profiled the expression of neurotransmitter receptor gene families in the EN subtypes. Interestingly, neurons in stage 1 and 2 showed the same phenomenon of falling into one of three major neurotransmitter responsive groups: NO/serotonin/GABA/glutamate responsive, acetylcholine responsive, or dopamine responsive (**Figure S12J and K**). For example, at stage 1, EN 3-5,7 and 8 neurons are predicted to be responsive to NO, serotonin, GABA and glutamate, EN 2 and 6 neurons are predicted to be responsive to acetylcholine and a subset of EN 1 neurons are predicted to be responsive to dopamine (**Figure 2A, Figure S12J**). It is important to note that many individual neurotransmitter receptor genes within the same family were differentially expressed between our enteric neuron subtypes (**Figure S9C, Figure S10C**). For example, within the family of acetylcholine receptors expressed by stage 1 enteric neuron subtypes, CHRM1 and CHRNA10 are exclusively expressed by EN 6, while CHRNB3 is exclusively expressed by EN 5 (**Figure S9C**).

These observations, in conjunction with the multiple neurotransmitter synthesis properties present in individual enteric neurons, highlights the complexity of ENS circuits, where a single neuron may both synthesize and respond to multiple neurotransmitters. Further, neurons can show subtype-specific responses to neurotransmitters depending on the receptor family member expression.

To verify that these complex neurochemical and transcriptional identities are physiologically relevant, we applied the same characterization criteria to both primary mouse and human ENS datasets (Drokhlyansky et al., 2020; Morarach et al., 2021) (**Figure S13A and B**). Interestingly, previously annotated subtypes are predicted to contain enteric neurons that synthesize multiple neurotransmitters, confirming our *in vitro* observations (**Figure S13A and B**). We then compared the overall abundance of neurons within each neurochemical class across each dataset irrespective of whether a neuron is predicted to synthesize multiple neurotransmitters (**Figure 2I**). We found that our ganglioids recapitulate temporal features of ENS development and maturation, such as the increase in NO neurons and loss of catecholaminergic neurons over time (Baetge and Gershon, 1989; Baetge et al., 1990; Bergner et al., 2014; Lake and Heuckeroth, 2013; Obermayr et al., 2013) (**Figure 2I**). Additionally, by concatenating the individually predicted neurochemical identities, we found that primary enteric neurons also contain complex neurochemical identities, where neurons are predicted to synthesize either one, two, three or more neurotransmitters (**Figure 2J and K**). We confirmed this by immunostaining of primary human colon and identifying neurons that were positive for both GABA and NOS1 or GABA and CHAT (**Figure 2L**). Interestingly, hPCS-derived enteric ganglioid neurons show relatively similar proportions of neurotransmitter complexity categories to primary neurons, and the neurochemical complexity appears to change during development in mice (**Figure 2K**). The breakdown of neurons belonging to each single and double neurochemical class confirms the presence of similar types of neurons across all datasets (**Figure 2M**).

### hPSC-derived enteric ganglioids recapitulate the glial diversity of the human ENS

Enteric glia play crucial roles in ENS physiology and disease but their molecular and functional characteristics have remained elusive. We showed that our stage 2 enteric ganglioids and 2D ENS cultures contain glia (**Figure 1G** **and H, Figure S6A**). To characterize these hPSC-derived enteric glia and determine if they recapitulate the transcriptional properties of primary enteric glia (Drokhlyansky et al., 2020; Morarach et al., 2021), we sub-clustered the glial population (**Figure 3A**) in enteric ganglioids and 2D ENS datasets and performed independent sub-clustering analyses of the primary glia sequenced by Drokhlyansky et al. and Morarach et al. We identified four glial subtypes (Glia 1-4) in the enteric ganglioids in two independent replicates (**Figure 3B**). Similarly, clustering of the primary glia in the Drokhlyansky et. al. human dataset showed four distinct subtypes (pGlia 1-4), differing from the 6 subtypes (three shared and three patient-specific subtypes) originally annotated by the authors (**Figure 3C**). Interestingly, visualizing the proportion of glial subtypes isolated from each patient sample in the primary human dataset suggests that the representation of glial subtypes varies from patient to patient possibly due to differences in sample collection (**Figure 3D**). Further, we subclustered the transcriptionally distinct glial lineages in our 2D ENS cultures and previously published mouse datasets (**Figure S14A-D**). All glial subtypes in 2D ENS cultures, enteric ganglioids and primary datasets expressed canonical glial markers (**Figure 3E, Figure S14E**). We confirmed this using immunofluorescence staining of S100 and GFAP in enteric ganglioids and primary human colon tissue (**Figure 3F**). Intriguingly, GFAP transcript was not detected in all glial populations and was restricted to Glia 1 in enteric ganglioids, Glia 1 and Glia 4 in 2D ENS cultures, and was low in human primary glial populations (**Figure 3E, Figure S14E**). Immunofluorescence staining of S100 and GFAP confirmed that these markers are not co-expressed in all glial cells (**Figure 3G**). To determine similarities between glial subtypes between the 3D and 2D subclusters, we next compared their transcriptional signatures. Module scoring and Spearman correlation revealed that a single 2D subtype shared the Glia 2 and 3 signatures in 3D enteric ganglioids (**Figure S14F and G**), confirming that all glial subtypes in enteric ganglioids are present in the 2D ENS cultures (**Figure S14F**). Module scoring showed that pGlia 1 is the most transcriptionally similar to Glia 1 and 4 subtypes, whereas pGlia 4 is most similar to the Glia 2 and 3 subtypes (**Figure 3H**). These data demonstrate that our 2D ENS cultures and enteric ganglioids capture the glial diversity of the human ENS.

**Figure 3.**
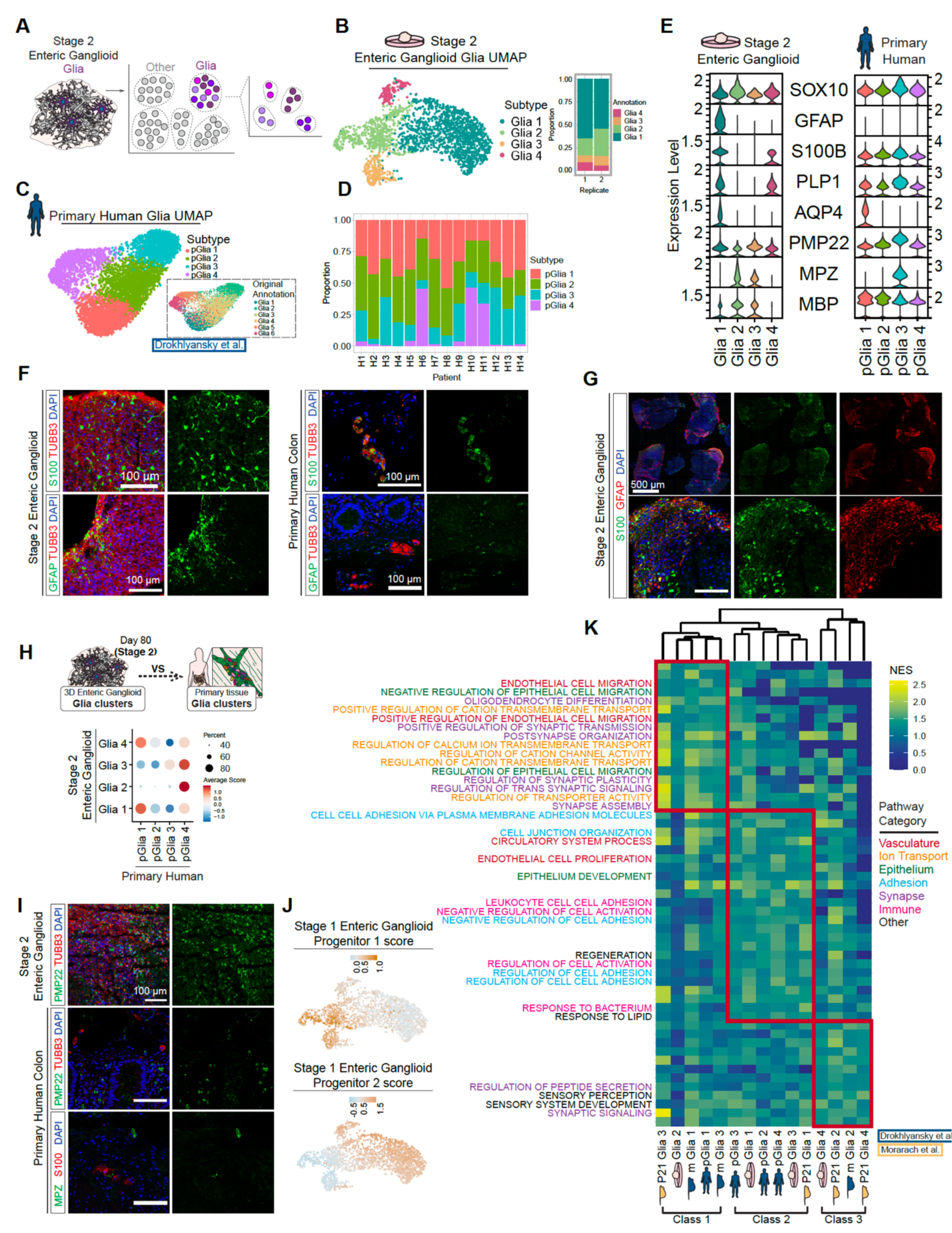
hPSC-derived enteric ganglioids model human ENS glial diversity. **A)** Schematic of snRNA-seq analysis and subsequent glial subclustering of stage 2 enteric ganglioids. **B)** UMAP of glial subtypes present in stage 2 enteric ganglioid (snRNA-seq, left) and distribution of glial subtypes in biological replicates of enteric ganglioid cultures (right). **C)** UMAP of enteric glial subtypes present in a primary adult human dataset **D)** Distribution of enteric glial subtype representation in individual human tissue samples. **E)** Violin stack plot of the expression of canonical glial markers in stage 2 enteric ganglioid and adult human glial subtypes. **F)** Immunofluorescence staining of canonical glial markers GFAP and S100 in stage 2 enteric ganglioids and human primary colon tissue. **G)** Co-staining of GFAP and S100 in stage 2 enteric ganglioids. **H)** Dot plot of the average module scores of stage 2 enteric ganglioid glial subtype transcriptional signatures in adult human enteric glial subtypes. **I)** Immunostaining of myelinating markers in human colon and enteric ganglioids. PMP22 expression in stage 2 enteric ganglioid (top), PMP22 (middle) and MPZ (bottom) expression in human colon. **J)** Feature plots showing the module scores of stage 1 enteric ganglioid progenitor 1 (top) and 2 (bottom) transcriptional signatures in ganglioid glia cells. **K)** Heatmap showing the normalized enrichment scores of GO pathways enriched in each glia class determined by hierarchical clustering.

Remarkably, we detected high level expression of myelinating markers PMP22, MPZ and MBP in our cultures (**Figure 3E**). Similarly, MBP transcript was present in all four human primary enteric glial subtypes, and pGlia3 showed MPZ expression (**Figure 3E**). Immunofluorescence staining confirmed the expression of myelin markers in stage 2 ganglioids and human primary colon tissue (**Figure 3I**). This is intriguing given the longstanding assumption that myelination does not occur in the ENS While S100B and PLP1 were expressed by all pGlia subtypes, they were only detected in Glia 1 and Glia 4 (**Figure 3E**). On the other hand, MPZ and MBP were predominantly expressed by the other two subtypes Glia 2 and 3 (**Figure 3E**). The mutually exclusive expression pattern of some of the canonical glial markers in our enteric ganglioids prompted us to explore their developmental origin. We aimed to infer lineage relationships between our stage 1 progenitor populations and the Glia subtypes. We found that Glia 2 and 3 shared a similar signature with progenitor 1, whereas Glia 1 and 4 were most similar to the progenitor 2 population (**Figure 3J**). This may suggest that unique enteric progenitor populations give rise to distinct MPZ**^+^**/MBP**^+^** and GFAP**^+^**/AQP4**^+^**/S100B**^+^**/PLP1**^+^** enteric glia (**Figure 3E**).

Given that enteric glial diversity has not been comprehensively and transcriptionally profiled, we performed deeper characterization of our stage 2 enteric ganglioid glial populations. Many transcription factors, neurotransmitter receptors, neuropeptide receptors, cytokines and their receptors, secreted signaling ligands and their receptors, and surface markers were exclusively expressed by each glial subtype (**Figure S15 A-H**), whereas other genes in these categories are differentially expressed between subtypes (**Figure S16 A-H**).

Further, we developed an approach to compare functional features of the glial subtypes in our enteric ganglioids with primary glial subtypes. We performed gene set enrichment analysis (GSEA) using the biological function gene ontology (GO) gene sets on the significantly upregulated gene lists of each mature glia subtype. Next we performed hierarchical clustering of the glial subtypes across all datasets based on the normalized enrichment score of all enriched GO terms present in at least one glial subtype. This analysis revealed three overarching classes of enteric glia conserved between mouse and human (**Figure S17A**). Inspection of the GO terms enriched in all glia of a particular class revealed diverse predicted functions of each class (**Figure 3K**). Class 1 glia are enriched for terms related to synapse regulation and ion transport while class 2 glia display terms related to adhesion and immune function. Both class 1 and 2 glia also contain terms related to epithelial and endothelial regulation. Interestingly, class 3 glia also show terms related to synapse regulation but uniquely contain terms specific to sensory processes. We next generated myenteric and submucosal glial signatures based on the differentially expressed genes of all primary glia isolated from each plexus. We used these signatures to predict plexus identities for each glial subtype in the human datasets. Similar to the neurons, scoring of the primary human glial subtypes showed mutual exclusion of the tissue layer signature, with pGlia 1 and 2 scoring positively for submucosal and pGlia 3 and 4 scoring positively for myenteric (**Figure S17B**). Interestingly, pGlia 4 is predominantly located in the myenteric signature (**Figure S17 C**). Similar to the neurons, scoring Glia 1-4 with the plexus signatures suggests that our cultures generate glia from both the myenteric and submucosal layers (**Figure S17D**). All together, these data indicate that our hPSC-derived enteric glia recapitulate the transcriptional, functional, and regional features of the primary human enteric glia.

### Enteric ganglioids enable comprehensive characterization of human NO neurons

GI motility is directly controlled by the enteric excitatory and inhibitory motor neurons. A large subset of inhibitory neurons use NOS1 to synthesize the neurotransmitter NO that induces relaxation in the smooth muscle tissue(Bredt et al., 1990; Bult et al., 1990; Ward et al., 1992; Young et al., 1992). NO is also an important regulator of mucosal integrity and barrier function. Enteric NO neurons are particularly important due to their involvement in a broad range of motility disorders. Selective loss and dysfunction of NO neurons have been associated with muscular hypercontractility underlying many dysmotility conditions such as achalasia, gastroparesis, intestinal pseudo-obstruction and colonic inertia (Bódi et al., 2019; Rivera et al., 2011).

Our hPSC-derived 2D ENS cultures and enteric ganglioids comprise a diverse population of neurons including the NO neurotransmitter identity. Having access to this subtype of neurons prompted us to perform deeper characterization of their molecular and functional identities and develop assays to understand and modulate their activity. To facilitate strategies for studying NO neurons *in vitro*, we generated a hESC NOS1::GFP line by inserting a GFP cassette under control of the endogenous NOS1 promoter using CRISPR/Cas9 knock-in technique (**Figure S18A**). Following our ENS induction protocol NOS1::GFP hESCs gave rise to mature cultures with NO neurons co-expressing GFP and NOS1 (**Figure S18B and C**). We performed bulk RNA-sequencing (bulk RNA-seq) on FACS purified NOS1::GFP**^+^**/CD24**^+^** and NOS1::GFP^-^/CD24**^+^**, using CD24 as a marker for neurons, and identified differentially expressed genes in NO neurons (**Figure S18D**). In parallel, by snRNA-seq profiling of our stage 1 enteric ganglioids, the NO neurons were identified by their expression of the key marker gene, NOS1, and selected metabolic and NO transport genes (**Figure S12A-F**, **Figure 4A and B**, **Table S1**). To determine the molecular diversity within the NO neurons, we performed further sub-clustering and identified 5 subtypes (NO 1-5, **Figure 4B and C, Figure S18E and F**). In addition to NOS1, these clusters showed enrichment for other NO biosynthesis pathway genes, confirming their shared NO identity (**Figure 4D**). We module scored NO neuron enriched genes identified by bulk RNA-seq in our snRNA-seq clusters and revealed positive enrichment for all NO 1-5, and in particular NO 3, relative to other neurons further confirming the reporter line reliability (**Figure S18G**). We assessed the transcriptional profile of a number of gene categories in NO neurons by combining bulk- and snRNA-seq data and revealed many transcription factors, neuropeptides and their receptors, neurotransmitter receptors, cytokines and their receptors, secreted ligands and their receptors and surface markers were differentially expressed in NO neurons relative to other neurons (**Figure S19A-J**). The bulk RNAseq data with higher sequencing depth confirmed many of the expression patterns observed in the snRNA-seq. For example, neurotransmitter receptor GABRA3 and neuropeptide SCG2 were enriched, while secreted ligand SEMA3A and ligand receptor DDR1 were depleted in NO neurons (**Figure S19C, H and I**).

**Figure 4.**
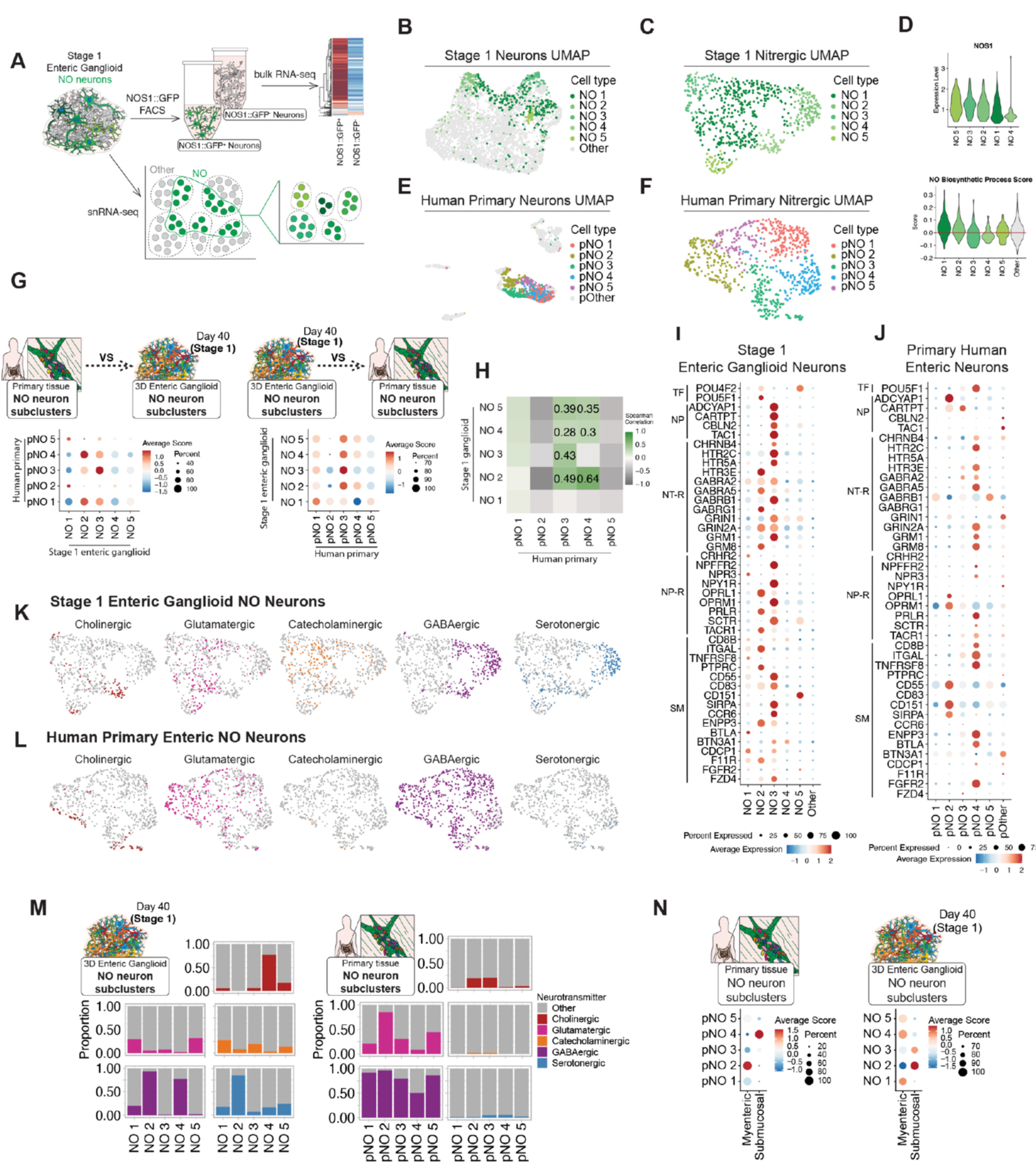
Enteric ganglioids recapitulate the NO neuron diversity of the human ENS. **A)** Schematic of CD24**^+^**/NOS1:GFP**^+^** FACS sorted neurons’ bulk RNA-seq analysis (top) and snRNA-seq analysis and subsequent NO neuron subclustering of stage 1 enteric ganglioids (bottom). **B)** snRNA-seq UMAP of NO subtypes present in stage 1 enteric ganglioid neurons. **C)** snRNA-seq UMAP of subclustered NO neuron subtypes from stage 1 enteric ganglioids. **D)** Violin plot of (top) NOS1 expression and (bottom) module scoring for nitric oxide biosynthesis gene ontology (GO) term genes by stage 1 enteric ganglioid NO subtypes. **E)** UMAP of pNO subtypes present in adult human enteric neurons. **F)** UMAP of subclustered pNO subtypes from adult human enteric neurons. **G)** Dot plot of the average module scores of adult human pNO neuron subtype transcriptional signatures in stage 1 enteric ganglioid NO neuron subtypes (left), and stage 1 enteric ganglioid NO neuron subtype transcriptional signatures in adult human pNO neuron subtypes (right). **H)** Heatmap matrix of Spearman correlations based on scaled expression of 3000 anchor features shared significantly variable genes (or anchor features) between adult human (x-axis) and stage 1 enteric ganglioid (y-axis) NO neuron subtypes. **I and J**) Dot plot of the scaled average expression of NO neurons specific transcription factors (TF), neuropeptides (NP), neurotransmitter receptors (NT-R), neuropeptide receptors (NP-R), and surface markers (SM) in stage 1 enteric ganglioid (**I**) and adult human (**J**) NO neuron subtypes versus non-NO neurons. **K and L**) Feature plots of predicted neurotransmitter producing neuron identities in stage 1 enteric ganglioid (**K**) and adult human (**L**) subclustered NO neurons. **M)** Distribution of neurochemical identities in stage 1 enteric ganglioid (left) and adult human (right) NO neuron subtypes versus non-NO neurons. **N)** Dot plot of the average module scores for myenteric and submucosal neuron transcriptional signatures in stage 1 enteric ganglioid (left) and adult human (right) NO neuron subtypes versus non-NO neurons.

Similar to enteric ganglioids, subclustering NO neurons in the adult human primary ENS dataset generated by (Drokhlyansky et al., 2020) identified five primary NO neuron subtypes (pNO 1-5 **Figure 4E and F**). Module scoring revealed similarities between NO 1-5 and pNO 1-5 (**Figure 4G**). Similarly, Spearman correlation analysis based on the expression of 3000 anchor features showed correlations for NO 2-5 with pNO 3, as well as NO 2,4,5 with pNO 4 (**Figure 4H**). Deeper characterization of these subclusters revealed differential expression patterns of some NO neuron specific features (**Figure 4I and J**, **Table S2**) such as serotonin receptors (HTR2C, HTR5A and HTR3E), GABA receptors (GABRA2 and GABRG1), glutamate receptors (GRIN1, GRIN2A, GRM1 and GRM8), and the opioid receptor OPRM1 (**Figure 4I and J**, **Table S2**). For example, we checked the expression of surface markers, transcription factors, neuropeptides and their receptors and identified genes that were specific to NO neurons, but showed subcluster specific pattern of expression, such as POU5F1, CARTPT, HTR3E, NPFFR2 and BTLA (**Figure 4I and J**, **Table S2**). These novel markers of NO neuron subtypes can be utilized for identification and further functional characterizations of unique NO neurons in both our *in vitro* cultures and from primary tissue samples. We further detected cholinergic, glutamatergic, catecholaminergic, GABAergic and serotonergic identities within the NO neuron subtypes in both datasets (**Figure 4K-M**). These data indicate that enteric NO neurons are transcriptionally diverse and may have distinct functional features.

To determine if these transcriptional differences may reflect differences in localization in the tissue, we used the previously generated neuron-specific myenteric and submucosal gene modules to identify their plexus identity. In the primary dataset, this analysis revealed one highly specific myenteric cluster (pNO 2) and one highly specific submucosal cluster (pNO 4) (**Figure 4N** **left**). Our ganglioid neurons similarly showed alignment with either a myenteric or submucosal identity (**Figure 4N** **right**).

### hPSC-derived ENS models identify modulators of NO neurons that promote colonic motility

Given the significant role of enteric NO neurons in GI motility and their selective vulnerability in a wide range of congenital and acquired enteric neuropathies (Bódi et al., 2019; Rivera et al., 2011), there has been a great interest in establishing strategies to regulate their function. Factors that modulate NO neuron activity and increase NO release will facilitate the identification of potential drug targets for treatment of enteric neuropathies. Hence, we leveraged our scalable ENS culture platforms to screen for compounds that induce NO neuron activity.

We developed a screening strategy for NO neuron activity based on the induction of cFOS expression as a readout. In order to evaluate cFOS as an accurate read-out of neurochemical-induced activity we performed side by side cFOS flow cytometry analysis and MEA neuronal firing measurements in cultures treated with epinephrine, which is known to stimulate enteric neurons. Epinephrine induced neuronal cFOS expression and resulted in increased electrical firing of ganglioid neurons. This provides a scalable read-out of activity that is suitable for high-throughput screens. (**Figure S20A-C**).

First, we performed a cFos induction screen, where NOS1::GFP enteric ganglioid cells were exposed to a library of 582 neuromodulators (Selleck neuronal signaling library^TM^) and co-expression of cFOS and GFP was measured to quantify NO neuron activity (**Figure 5A, Figure S20D**). We identified 20 compounds that increased the proportion of NO neurons in cFOS**^+^** cells with z-score>1.5. To identify the mechanisms involved in cFOS expression in NO neurons, we compiled and classified the list of target proteins and discovered multiple shared protein classes. In particular, these proteins converged on serotonin receptors, sodium channels, acetylcholine receptors, glutamate receptors, adrenergic receptors, histamine and opioid receptors and dopamine receptors (**Figure 5B**).

**Figure 5:**
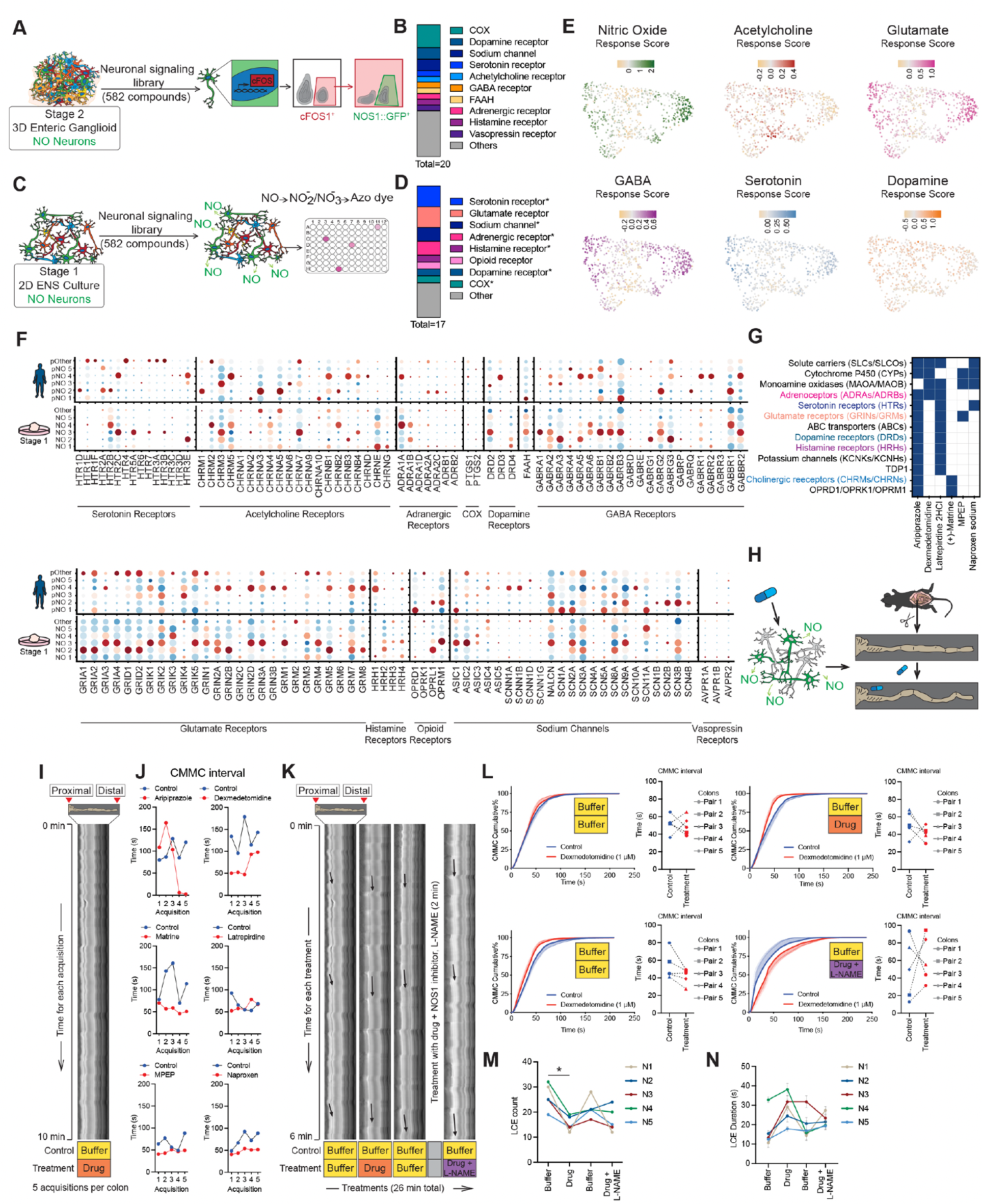
NO neuron specific functional screening identifies modulators of colonic motility. **A)** Schematic representation of a high-throughput flow cytometry-based screening to identify compounds that induce cFOS expression in hESC-derived stage 2 enteric ganglioid NO neurons. **B)** Target classes of the hits identified in enteric NO neuron cFOS induction screening (**Figure S20D**, red dots). **C)** Schematic representation of a high-throughput calorimetry-based screening to identify compounds that induce NO release in hESC-derived stage 1 2D ENS cultures. **D)** Target classes of the hits identified in NO release screening (**Figure S20E**, red dots). Protein classes that are in common with (**B**) are indicated with asterisks. **E)** Feature plots showing the predicted responsiveness of subclustered stage 1 ganglioid enteric NO neurons to neurotransmitters by module scoring of neurotransmitter receptor gene families. **F)** Dot plot of the expression of genes belonging to the target classes shown in B and D in hESC-derived stage 1 and primary human enteric nitrergic neuron subtypes versus all other neurons. **G)** Combined protein target analysis for selected screening hits showing shared protein classes. Color code matches the target classes in (**B**) and (**D**). **H)** Schematic representation of testing the effect of selected candidate hits (listed in (**G**)) on mouse colonic motility *ex vivo*. **I)** Representative spatiotemporal map of mouse colon contractions along the proximal-distal axis over a 10-min period. **J)** Quantification of colonic migrating motor complexes (CMMC) intervals at 75th percentile of CMMC cumulative percentage (**Figure S21A**) for selected hit compounds. **K)** Experimental design for measuring the effect of selected candidate hits on mouse colonic motility *ex vivo*. Representative spatiotemporal maps of a mouse colon contraction along the proximal-distal axis over a 26-min period. Three representative longitudinal contractile events (LCEs) are shown per condition (arrows). **L)** Diagrams of CMMC cumulative percentile and quantification of CMMC interval (time difference between two consecutive contractions) at 75th percentile for dexmedetomidine. Mean and SEM error bars for 5 pairs of untreated and drug-treated mouse colons are shown. **M)** Total number of colonic longitudinal contractile events (LCEs) within each 6-min treatment condition for 5 dexmedetomidine-treated mouse colons measured from spatiotemporal maps. *: p-value < 0.05. **N)** Mean of LCE duration calculated for three LCEs within each 6-min treatment (1 in the beginning, 1 in the middle, and 1 in the end of each spatiotemporal map, see (**K**)). Data are shown for 5 dexmedetomidine-treated mouse colons. SEM error bars are shown.

In an independent functional screen, we established a high-throughput read-out for assessing NO neuron activity. We utilized a commercially available kit that enables NO detection in the media. Upon release into the media, NO is spontaneously oxidized to nitrate. The kit uses nitrate reductase to convert nitrate to nitrite that is then detected as a colored azo dye. We incubated 2D ENS cultures with the neuromodulators library and measured NO release using calorimetry (**Figure 5C**). We identified 17 compounds that remarkably enhanced NO concentration in the supernatant with z-score>2.0 (**Figure S20E**). Neuromodulators that induced NO release in our ENS cultures were diverse but were predicted to commonly target protein classes including serotonin receptors, sodium channels, acetylcholine receptors, glutamate receptors, adrenergic receptors and opioid receptors (**Figure 5D**).

Interestingly, there was a high degree of similarity between the predicted targets from the cFOS induction and NO release screens (**Figure 5B and D, Figure S20F**). These targets included receptors for neurotransmitters such as serotonin and dopamine. Module scoring the neurotransmitter receptor gene families in our hPSC-derived stage 1 enteric NO neurons snRNA-seq data confirmed that NO neurons broadly express receptors for NO, serotonin, GABA, glutamate, acetylcholine or dopamine (**Figure 5E**). Interestingly, compared to other neuronal subtypes, stage 1 NO neuron clusters were enriched for all predicted hit targets (**Figure S20G**). In particular, NO 3 cluster scored high for the expression of the majority of NO neuron modulator target classes (**Figure S20H**). Profiling the expression of individual genes in each target protein category in ganglioids and primary human ENS revealed notable subtype-specific expression patterns between NO neurons. For example, GABA receptor genes were predominantly expressed by NO 2, NO 3 and pNO 4 subtypes, while the expression of acetylcholine receptor genes was less specific to a particular subtype (**Figure 5F**).

We then selected a subset of hits representing different target classes, prioritizing FDA approved compounds for follow up analyses (**Figure 5K, Figure S20F**). For selected compounds, we performed a more comprehensive and integrated target analysis by combining reported experimental data (Binding DB) and computational methods (SEA, Carlsbad, Dinies, Swisstarget, Superdrug, Pubchem Bioassays, **Figure 5G**). We next tested the effects of these compounds on colonic motility in organ bath assays (**Figure 5H**). In these assays, we maintained resected segments of mouse colon in a physiologic buffer to study motility patterns using video recording. In the first experiment we tested the effect of all selected drug candidates on mouse colonic motility, *ex vivo* (**Figure 5I**). In each experiment, we tested an untreated control and a drug-treated colon sample side by side during five consecutive 10-min acquisitions. For each acquisition, instead of analyzing fecal outputs, which are generally variable, we performed more sophisticated contraction analysis by generating spatiotemporal maps from video data based on the changes in the colonic diameter over time, and used to calculate the rate of colonic migrating motor complexes (CMMC) and slow waves (SW). CMMCs are rhythmic propulsive contractions initiated by the ENS while SWs are mediated through the pacemaking activity of the interstitial cells of Cajal(Barajas-López and Huizinga, 1989; Burns et al., 1996; Fida et al., 1997; Lyster et al., 1995; Smith et al., 1987). To probe the dynamics of CMMC and SW events during each acquisition, we generated cumulative percent graphs (**Figure S21A and B**) and calculated the intervals at the 75th percentile (**Figure 5J, Figure S21C**). Compounds that showed promising effects on lowering CMMC intervals relative to the untreated condition were chosen for follow-up assessment (i.e. aripiprazole, dexmedetomidine, matrine, MPEP) (**Figure 5J, Figure S21A-C**). To assess whether the compounds mediated their effect on CMMCs through modulating NO release, we performed sequential drug treatments in the presence and absence of the NOS1 inhibitor N(omega)-nitro-L-arginine methyl ester (L-NAME). Each experiment consisted of four 6-min acquisitions on five independent sample pairs in the control and drug-treated groups (**Figure 5K**). Of the tested drugs, the adrenergic receptor agonist dexmedetomidine, decreased CMMC intervals in four out of five colon samples, an effect that was absent when colons were treated with dexmedetomidine + L-NAME simultaneously (**Figure 5L, Figure S22A-F, Figure S23A and B, Video S1**). SWs were not affected by the drug treatment (**Figure S24A-H**). In addition to CMMCs, we quantified the effects of compounds on colonic motility by annotating and measuring anterograde contractile events detected in the spatiotemporal map. These events were termed “longitudinal contractile events” (LCE) and highlighted by representative arrows in **Figure 5K**. In dexmedetomidine-treated colons, we observed a decrease in the total number of LCEs (**Figure 5M**) and an increasing trend in their average duration (**Figure 5N**) in all five replicates. These effects were reversed after the drug removal and blocked by L-NAME co-treatment (**Figure 5M and N**). These results provide a blueprint for leveraging *in vitro* human ENS models to uncover mechanisms that regulate GI motility which are capable of identifying therapies that target specific ENS populations.

### High-throughput small molecule screen reveals PDGFR inhibition as a driver of NO neuron induction

To evaluate the potential of hPSC-derived cultures to model human ENS development, we set out to define the mechanism of NO neuron specification *in vitro*. Searching for pathways that regulate NO neuron differentiation, we performed a high-throughput small molecule screen. Identifying distinct pathways and chemical modulators that promote NO neuron induction could provide insights on NO neuron development and offer a strategy for the derivation of NO neuron enriched ENS cultures.

To identify compounds that induce NO neuron differentiation, we treated enteric crestospheres with 1694 compounds in the Selleck inhibitor library^TM^ and identified 12 hit compounds that increased the proportion of NOS1^+^ neurons by at least eight folds (**Figure 6A, Figure S25A and B**). In order to reveal the mechanisms by which these hit compounds enhanced NO neuron induction, we performed target prediction analysis by combining reported experimental data (Binding DB) and computational methods (SEA, Carlsbad, Dinies, Swisstarget, Superdrug, Pubchem Bioassays) (**Figure 6B**). After clustering the predicted protein targets, common patterns emerged for a subset of compounds. For example, PP121, ibrutinib, afatinib, and AMG-458 were all predicted to interact with EGFR, ERBBs, MAP, and TEC family kinases among others (**Figure 6B**). For follow-up analysis, we chose to focus on PP121, the hit with the highest %NOS1**^+^** fold increase in this subset of compounds. PP121 showed a dose-dependent effect on NO neuron induction efficiency as measured by flow cytometry (**Figure S25C**). To find the most effective treatment window for PP121-induced NO neuron induction, we treated the differentiating cultures for five days at various time points. Measuring GFP signal in stage 1 NOS1::GFP enteric ganglioids showed the highest induction efficiency for cells treated during day 15-20 (**Figure 6C and D, Figure S25D**).

**Figure 6:**
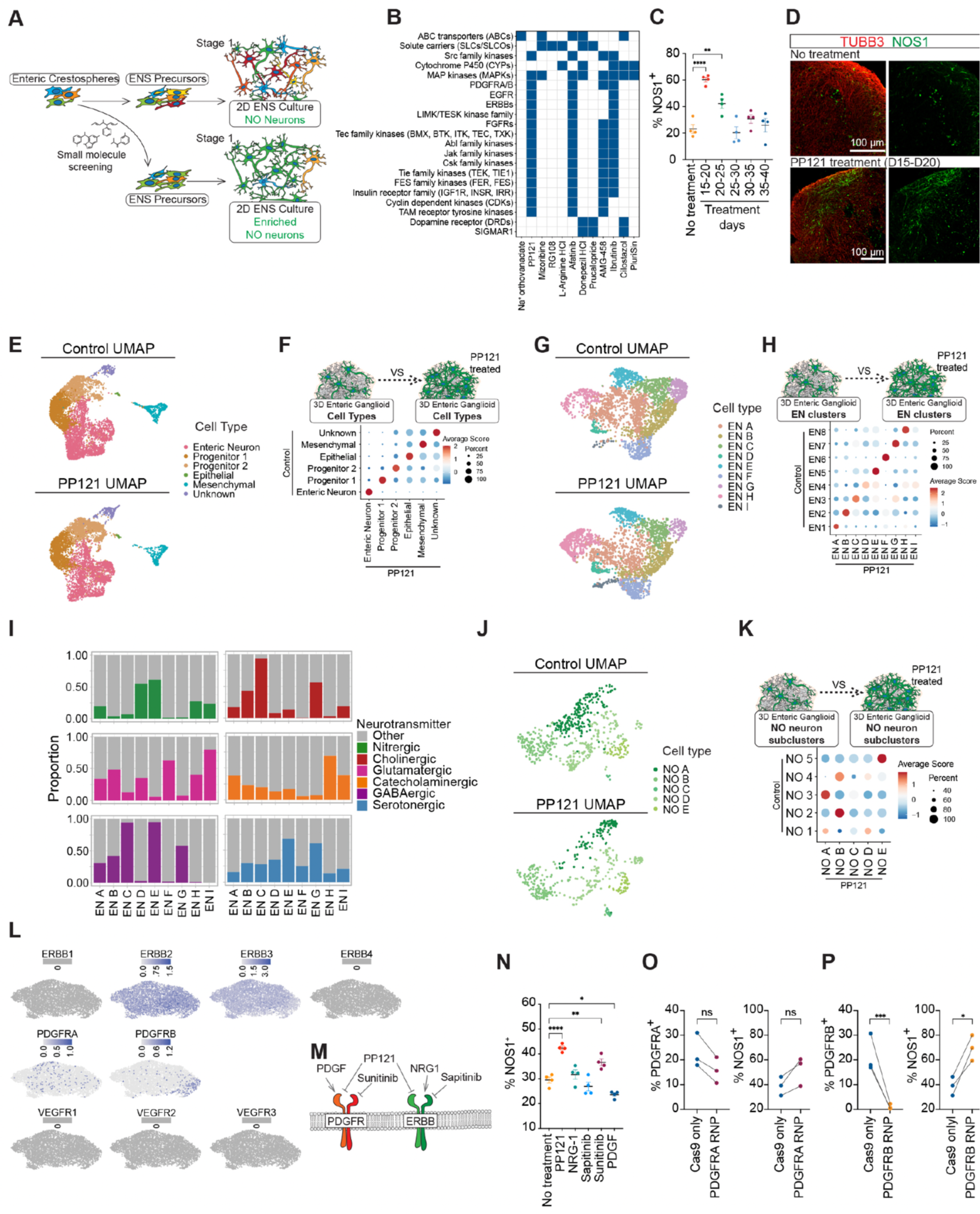
PDGFR inhibition promotes enteric NO neuron induction. **A)** Schematic representation of a high-throughput pharmacological screening to identify compounds that enrich NO neurons in hESC-derived 2D ENS cultures. **B)** Combined protein target analysis for the HTS top 12 hits showing shared protein classes between structurally similar hits. **C)** Effect of PP121 treatment window on NOS1::GFP induction efficiency. **D)** Immunofluorescence staining of NOS1 and neuronal TUBB3 in stage 1 enteric ganglioids treated with or without PP121 between days 15 and 20. **E)** Split UMAP of cell types present in stage 1 control (top) and PP121 treated (bottom) enteric ganglioid cultures. **F)** Dot plot of the average module scores of control only enteric ganglioid subtype transcriptional signatures in PP121 treated ganglioidl subtypes. **G)** Split UMAP of neuronal subtypes present in stage 1 control (top) and PP121 treated (bottom) enteric ganglioid cultures. **H)** Dot plot of the average module scores of control only neuronal subtype transcriptional signatures in PP121 treated ganglioid neuronal subtypes. **I)** Distribution of NO neuron subtypes in control versus PP121 treated stage 1 enteric ganglioid cultures. **J)** Split UMAP of subclustered NO subtypes present in stage 1 control (top) and PP121 treated (bottom) enteric ganglioid cultures. **K)** Dot plot of the average module scores of control only NO neuron subtype transcriptional signatures in PP121 treated ganglioid NO neuron subtypes. **L)** Feature plot showing the expression of ERBBs, PDGFRs and VEGFRs in D15 subclustered enteric crestospheres. **M)** Schematic of receptor tyrosine kinase (RTK) natural agonists and selected pharmacological antagonists including NO neuron enriching top hit PP121. **N)** Effect of RTK ligand treatment on stage 1 enteric ganglioid NO neuron induction. **O and P**) Effect of knocking out PDGFRA (**O**) and PDGFRB (**P**) in D15 enteric crestospheres on stage 1 enteric ganglioid NO neuron enrichment as measured by flow cytometry.

For the enrichment protocol to be reliable, it was important to confirm that PP121 treatment did not change the identity of our cell types. To compare PP121 treated and untreated stage 1 enteric ganglioids in single cell resolution, we performed snRNA-seq and combined both datasets. This analysis revealed that all cell types were represented in both conditions (**Figure 6E and E, Figure S26A**). Importantly, comparison of the average expression of all genes for matched PP121 treated and untreated cell types showed highly similar transcriptomes (R2 correlations >0.91), indicating that PP121 treatment did not change the transcriptional identity of cell types (**Figure S26B**). Interestingly, sub-clustering of the merged control and PP121 treated neurons revealed nine neuronal subtypes EN A-I (**Figure 6G**). The PP121 treated dataset subtypes showed high transcriptional similarity to EN 1-8 of the control only dataset (**Figure 6H**). EN cluster I consisted of mostly PP121 treated cells and few control cells and showed moderate transcriptional similarity to the control only EN cluster 4, suggesting that this neuronal subtype is present but rare in control cultures causing those neurons to cluster with the most similar subtype, EN 4 (**Figure 6H, Figure S26C**). Along with EN I which is roughly 25% nitrergic, PP121 treatment also enriched cultures for neuronal subtypes EN D and H (roughly 50% and 25% nitrergic, respectively), while EN A and G were less represented (**Figure 6I, Figure S26C**). Again, despite the changes in subtype abundance, control and PP121 treated neurons of the same subtype showed similar transcriptomes (R2 correlations >0.88) (**Figure S26D**). Further sub-clustering of merged control and PP121 treated nitrergic neurons revealed an enrichment for Nitrergic B (most similar to control only Nitrergic 2) and a rare population, Nitrergic C (most similar to control only Nitrergic 3) (**Figure 6J and K, Figure S26E**). Transcriptome comparison again showed highly similar gene expression of control and PP121 treated nitrergic neurons of the same subtype (R2 correlations >.7) with the highest variance between Nitrergic C neurons, likely due to the small number of neurons in this cluster (**Figure S26F**). Altogether, this data suggests that early treatment of ganglioids with PP121 causes changes in the abundance of neuronal subtypes normally found in untreated cultures without affecting the gene expression patterns of the subtypes that arise.

The ability to purify enteric NO neurons is of great interest especially for applications such as cell therapy. Access to NOS1::GFP reporter line and the ability to direct the differentiation towards NO neurons using PP121, allowed us to search for FACS-compatible surface markers for these cells. We screened a panel of 242 antibodies for human cell surface molecules (BD lyoplate) and measured GFP and surface antigen expression signals by flow cytometry (**Figure S27A**). We identified 27 antibodies that stained at least 50% of NOS1 neurons (%CD^+^GFP^+^ in GFP^+^ population, **Figure S27B top**). To identify the most specific candidates among these hits, we looked for antibodies with a >70% CD^+^GFP^+^ to CD^+^ staining ratio (**Figure S27B bottom**). CD47, CD49e, CD59, CD90, and CD181 met both criteria (**Figure S27C**). We further confirmed the expression and enrichment of CD47, CD49e, CD59 and CD90 in stage 1 ganglioid and NO neuron clusters in the human primary snRNA-seq dataset (**Figure S27D and E**). As an example, we further confirmed the localization of CD47 in NO neurons in human primary colonic myenteric ganglia using immunohistochemistry (**Figure S27F**). In addition to identifying antibodies to specifically enrich NO neurons, we found twelve antibodies that stained >70% of ganglioid cells and could serve as pan enteric neuronal surface markers (CD24, CD45RA, CD57, CD63, CD71, CD121b, CD147, CD164, CD184, CD193, CD243, CD275) (**Figure S27G**). We confirmed the enriched expression of CD24 in our enteric neurons (snRNA-seq data) and also primary human colon myenteric ganglion (**Figure S27 H and I**).

To determine the mechanism by which PP121 induced NO neuron enrichment in ganglioids, we used a combination of pharmacological and genetic approaches. PP121 is a multi-targeted receptor tyrosine kinase (RTK) inhibitor with known inhibitory activity on PDGFRs, VEGFRs and EGFRs (Apsel et al., 2008). Our crestosphere snRNA-seq analysis confirmed the expression of PDGFRA, PDGFRB, ERBB2 and ERBB3 while the mRNA for VEGFRs were not detectable (**Figure 6L**). We evaluated the induction efficiency of NO neurons in response to PDGF (PDGFR agonist), sunitinib (PDGFR and VEGFR antagonist), NRG1 (ERBBs agonist) and sapitinib (ERBBs antagonist) (**Figure 6M**). While NRG1 and sapitinib showed no significant effect on NO neuron induction, treatment with PDGF and sunitinib led to lower and higher NO neuron proportions respectively (**Figure 6N**). To genetically confirm the role of PDGFR signaling in NO neuron induction, we used CRISPR-Cas9 to knock-out PDGFRA and PDGFRB in our enteric crestospheres and analyzed the percentage of NO neurons in stage 1 ganglioids. NO neurons were enriched in both PDGFRA and PDGFRB knock-out cultures further confirming the PP121 mechanism of action (**Figure 6O and P**).

### hESC-derived NOS1 neurons engraft in *Nos1*^-/-^ mouse colon

Developing an experimental system to study the human ENS *in vivo* opens a wide range of basic science and clinical opportunities. For example, human ENS xenografts will enable studying human neuronal circuitry *in vivo* and investigating ENS-CNS and ENS-immune system-microbiome communications. They also provide platforms for disease modeling and drug development. In addition, the limited regenerative capacity of the ENS highlights the importance of developing cell therapy approaches to replace the lost populations of neurons. There is currently no clinical intervention to replace the damaged or lost neurons caused by genetic and acquired ENS pathologies such as Hirschsprung disease and diabetes. We have previously shown hPSC-derived ENC precursors can successfully engraft *in vivo* (Fattahi et al., 2016). McCann et. al. have also shown the transplanted *ex vivo* cultured murine enteric neurospheres are able to rescue GI motility defects in *Nos1*^-/-^ mice (McCann et al., 2017). However, these neurospheres are heterogeneous populations containing only a small percentage of NO neurons. Additionally, obtaining sufficient numbers of neurospheres from human primary tissue poses a significant limitation for ultimate regenerative applications. Compared to ENC precursors, transplanting mature neurons provides a post-mitotic source of cells with a lower clinical risk of tumor formation. Obtaining highly enriched NO neuron cultures encouraged us to assess the transplantation potential of our enteric ganglioids. PP121 treated enteric ganglioids were injected in the wall of distal colon in immunocompromised *Nos1*^-/-^ (B6.129S4-*Nos1^tm1Plh^*^/J^) mice. Animals were sacrificed eight weeks post-surgery and colonic longitudinal muscle myenteric plexus (LMMP) preparations were assessed by fluorescence microscopy (**Figure 7A**). Transplanted cells were distinguished by the expression of human cytoplasmic marker SC121. Notably, we observed a remarkable number of SC121^+^ cells that had integrated along the length of the colon (**Figure 7B, Videos S2-4**). Engrafted cells were detected within, and outside of myenteric ganglia and many expressed NOS1 confirming their NO fate (**Figure 7C**, **Figure S28**). In addition to the clinically-significant cell therapy application, the developed human enteric ganglioid xenograft offers previously unachievable opportunities towards understanding development, physiology and pathophysiology of the human ENS *in vivo*.

**Figure 7:**
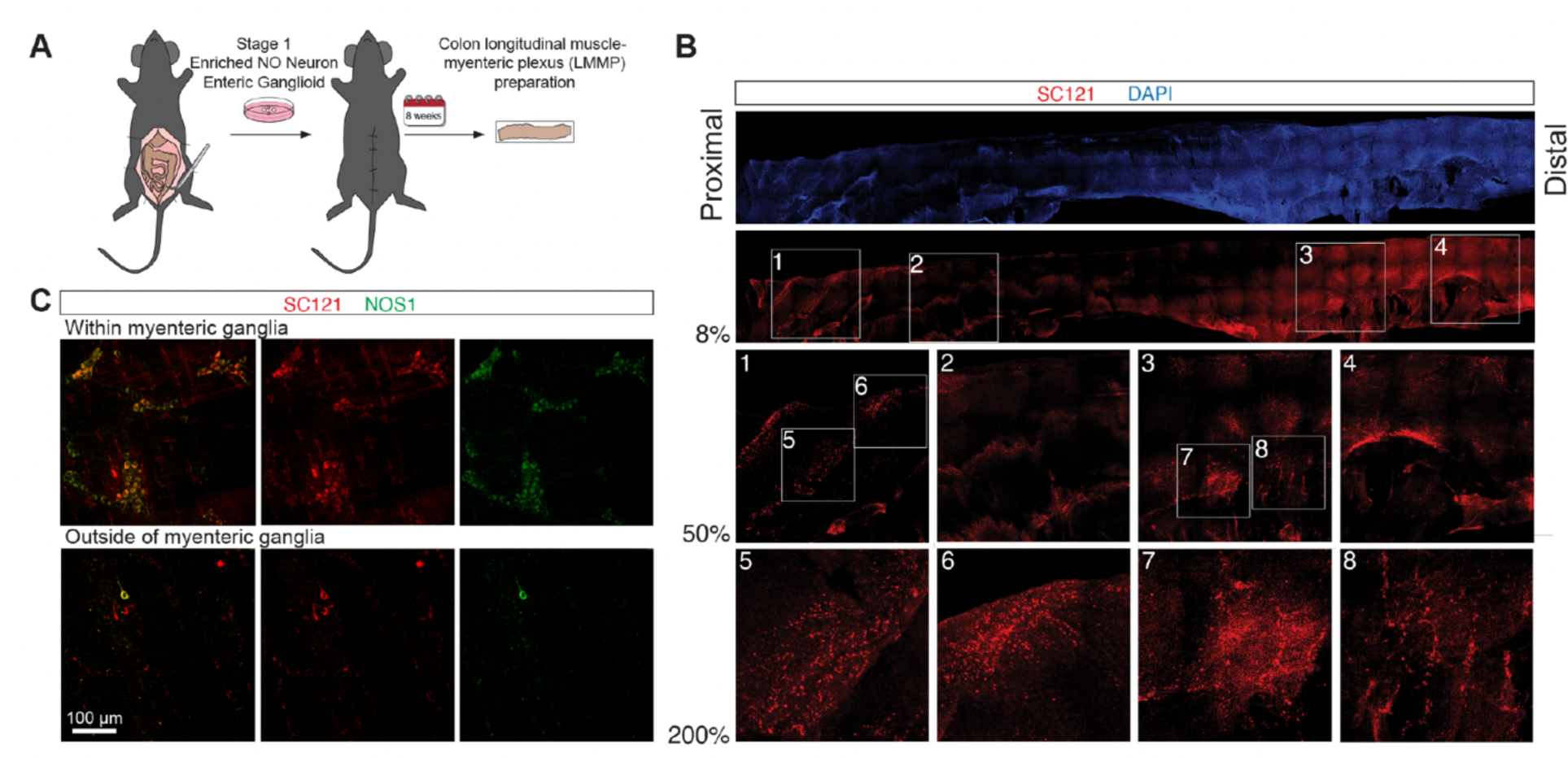
Extensive engraftment of hESC-derived enteric ganglioids in adult mouse colon. **A)** Schematic showing transplantation of hESC-derived stage 1 enteric ganglioids into mouse proximal colon. **B)** Engraftment of hESC-derived stage 1 enteric ganglioid cells into the entire length of mouse colon as shown by the expression of human cytoplasmic marker SC121 in red. **C)** Immunohistochemical analysis of human cytoplasmic protein SC121, and NO neuron marker NOS1 in *Nos1*^-/-^ mouse colon 8 weeks post transplantation.

## Discussion

The ENS is a complex network of enteric neurons and glia that controls all aspects of GI physiology (Long-Smith et al., 2020; Schneider et al., 2019; Yoo and Mazmanian, 2017) and plays a central role in initiation and progression of enteric neuropathies and diseases of the gut-brain axis (Camilleri, 2021; Niesler et al., 2021; Pesce et al., 2018). Nevertheless, our understanding of the ENS has been disproportionately affected by long standing technical challenges. Gaining access to human ENS requires invasive biopsies or surgeries as these cells only comprise 1% of the gut tissue(Drokhlyansky et al., 2020) and reside deep within muscular and mucosal layers. Moreover, large scale isolation and purification of ENS cells is extremely challenging. The majority of neuronal cell bodies are positioned in ganglia with their fragile projections extending to other parts of the gut tissue. In addition, there are no well-established surface markers for FACS-based purification of specific subtypes of enteric neurons or glia. Furthermore, animal models do not fully recapitulate the human ENS (patho)physiology. For example, rodents can well tolerate mutations that cause life-threatening enteric neuropathies in humans (Bondurand and Southard-Smith, 2016). Here, we present hPSC-derived ENS cultures as alternative models that overcome many of these challenges and enable major advances in the field of enteric neurobiology.

We thoroughly compared the composition of the hPSC-derived ENS platform against the recently published primary ENS datasets by snRNA-seq (Drokhlyansky et al., 2020; Morarach et al., 2021) and revealed diverse neuronal and glial subtypes that resemble the cellular diversity found *in vivo*. For example, the hPSC-derived enteric neurons express key markers and receptors for numerous hormones, neuropeptides and neurotransmitters that are known to exist in primary human ENS supporting the reliability and utility of our hPSC-based platform for modeling the human ENS. By clustering and further sub-clustering of our datasets we identified novel markers for each subtype offering opportunities for immunochemistry-based detection, purification, genetic manipulation, and reporter line development. Furthermore, investigation of the primary and hPSC-derived neurons shed light on the array of neurons with multiple neurochemical identities. These neurochemically diverse neurons greatly outnumber the ones that align with the traditionally and widely accepted belief that one-neuron expresses one-neurotransmitter. Although there have been immunohistochemical based reports of enteric neurons with multiple neurochemical identities (Qu et al., 2008), a comprehensive characterization has not been carried out before. The level of complexity revealed here could not be easily recognized and characterized by common lower-throughput staining based detection methods. This is of high scientific and medical value, furthering our understanding of the human ENS circuitry and autonomy, and informing the development of more targeted therapeutics with fewer side effects.

The autonomy of the ENS and its ability to perform diverse tasks independently relies on the diversification of its neuronal and glial components through elaborate fate specification processes. The precise developmental patterns that drive the differentiation of vagal neural crest into ENS progenitors that consequently mature into a myriad of neuronal and glial subtypes has remained elusive particularly in humans. Studying these complicated developmental patterns is extremely challenging due to the transient nature of many of the developmental states, technical limitations of isolating the tissue, and inter-species differences. Leveraging our stepwise ENS induction system, we generated high-resolution temporal maps revealing the complicated developmental programs that give rise to enteric neuron and glia. We begin the *in vitro* differentiation by inducing vagal and enteric neural crest that develop into enteric crestospheres, which we further differentiate into enteric neurons and glia. Interestingly, in long-term cultures we observed the appearance of enteric glial subclasses that resemble adult human primary glia. This resembles the developmental timeline in the CNS, where gliogenesis follows neurogenesis. Given the complexity of the processes influencing ENS development, it is not surprising that defects at any developmental stage lead to enteric neuropathies such as Hirschsprung disease (Lake and Heuckeroth, 2013; Rao and Gershon, 2018). Investigating the broad and cell-type specific developmental programs in the ENS provides an opportunity for understanding developmental neuropathies and facilitates the directed derivation of disease-relevant cell types.

An exceptional advantage of hPSC-derived cultures is their scalability. This is particularly important when the desired cell types are rare, and have very limited regenerative and proliferative capacity such as nervous tissue. Our ENS culture platforms have repeatedly proven to be reliable in providing scalable sources of ENS cell types that are compatible with applications that would otherwise be extremely challenging to implement, such as high-throughput screens. In particular, using our 2D ENS cultures we screened thousands of inhibitors to identify compounds that direct the differentiation towards the clinically valuable NO neurons. Investigating the mechanism of action of our top hits revealed pathways that are important in NO neuron fate specification. Using a combination of pharmacological and genetic approaches, we discovered the contribution of one such pathway, PDGFR signaling, in inducing NO neurons, which highlights the remarkable potential of hPSC-based platforms to uncover developmental mechanisms.

Our 2D and 3D ENS cultures are electrically active. Access to functional enteric neurons is extremely advantageous as it facilitates the basic understanding of neuronal circuits and cellular electrophysiology. Further, identifying cell-type specific neurochemical and functional characteristics is invaluable in drug development as it enables the identification of targeted neuromodulators and prediction of potential side effects through direct and indirect neurochemical mechanisms. As a proof of concept, we developed functional screening platforms to uncover candidate drugs that specifically modulate the activity of NO neurons. Interestingly, our hit compounds commonly target adrenergic, cholinergic and serotonergic receptors and sodium channels. Notably, these targets are overrepresented in NO neurons, highlighting the specificity of these compounds and their potential for further therapeutic development for GI indications. By testing a subset of these neuromodulators, we further demonstrated that these candidate drugs are capable of affecting colonic motility patterns in *ex vivo* organ bath assays. This is the first example of identifying candidate drugs for modulating GI motility by targeting a specific enteric neuron subtype. These findings showcase the reliability, robustness, and scalability of our hPSC derived ENS models.

Derivation of enteric ganglioids from hPSCs provides a scalable source of human ENS tissue for regenerative applications. Additionally, developing human ENS xenografts opens a wide range of basic science and clinical research avenues. In the last two decades, developing cell-based therapies for enteric neuropathies has been a major area of research (Alhawaj, 2021; Burns et al., 2016). However, a scalable source of human ENS cells suitable for transplantation is challenging to achieve. Here, we provide proof of concept results on extensive engraftment of NO neurons in *Nos1^-/-^* mice by transplanting ganglioids enriched for this neuronal subtype. Beyond cell therapy, these human ENS xenograft models provide a new experimental system for various purposes. First, these models enable the study of human ENS *in vivo* and facilitates the identification and development of therapeutic candidates with high specificity, efficacy and potency. Second, they may be used models to study human ENS pathologies *in vivo*, using strategies such as transplanting ganglioids harboring specific mutations, ganglioids exposed to specific stressors, or ganglioids derived from patient iPSCs. Third, transplanting ganglioids at different stages of differentiation enables comprehensive studies on cell fate specification and maturation in the human ENS. Finally, human ENS xenografts offer promising models for studying the crosstalk between the human ENS and local gut tissues, the CNS, and the microbiome.

Enteric neuropathies can affect any part of the GI tract at any stage of life, and represent some of the most challenging clinical disorders with no effective therapies. They can result from congenital defects affecting ENS development, can occur in response to changes in the tissue environment (toxins, microbes, immune system), or can emerge secondary to systemic diseases such as diabetes and obesity (Camilleri et al., 2011; Niesler et al., 2021; Yarandi and Srinivasan, 2014). The lack of efficient therapies stems from our inadequate understanding of ENS development, cellular architecture, and function. Our hPSC-derived 2D ENS cultures and enteric ganglioids provide human-based platforms to model enteric neuropathies. Genetic manipulation of neurons, glia, and their specific subtypes at different stages of their development is now possible. The effect of genetic background (healthy and patient-derived iPSCs) (Lai et al., 2017) and specific mutations as well as environmental stressors, infectious agents, metabolic toxins can now be studied via targeted and unbiased approaches. Furthermore, functional and fully characterized ENS cultures open avenues for investigating additional layers of complexity represented in gut physiology, such as crosstalk with the surrounding and distant tissues. For example, we can study motor functions by developing co-cultures with smooth muscle cells; investigate ENS-immune system communication by setting up co-cultures with immune cells, and interrogate ENS-gut microbiome interactions by exposing ENS cells to gut microbiome by-products.

Our hPSC differentiation strategy provides robust 2D and 3D ENS culture systems that enable developmental, molecular and functional mapping of human ENS. We provide key insights into physiological properties of NO neurons and identify the developmental programs required to specify this clinically relevant enteric neuron subtype. These models open up new avenues for drug discovery and regenerative medicine and offer a new framework for basic studies of enteric neurobiology.

## Acknowledgements

We thank UCSF Parnassus Flow Core, RRID:SCR_018206 for their excellent technical support. We acknowledge the UCSF Center for Advanced Technology (CAT), GenomicsCoLab and 10x Genomics team for high-throughput sequencing and other genomic analyses. We thank UCSF Biological Imaging Development CoLab (BIDC) for their excellent imaging technical support. We are grateful to S. Farahvashi for his excellent lab management support. We are grateful for the support from the UCSF Program for Breakthrough Biomedical Research and Sandler Foundation, the NIH Director’s New Innovator Award (DP2NS116769) and the National Institute of Diabetes and Digestive and Kidney Diseases (R01DK121169) grants to F.F. M.G.K. is supported by National Defense Science and Engineering Graduate Fellowship (00002116, M.G.K.). H.G. is supported by NIH (R01CA240984). H.M. is supported by Larry L. Hillblom Foundation postdoctoral fellowship.

We are grateful to Grant Hennig or providing the VolumetryG9a and Karlheinz Merkle and the Stanford Physics Machine Shop for help with design and fabrication of the motility monitor.

This work was supported by a ChEM-H Postdocs at the Interface Seed grant (S.D.), the Stanford Bio-X Undergraduate Research Program (R.K.S.), and by the Wu Tsai Neurosciences Institute, the Department of Neurosurgery at Stanford University School of Medicine, and a research grant from the Shurl & Kay Curci Foundation (J.A.K.).

We are grateful to Takeda Development Center Americas, Inc. for providing human colonic tissue sections and their financial support of this work.

## Author contributions

H.M. experiment design, writing of manuscript, maintenance and directed differentiation of hPSCs, cFOS induction and NO-release high-throughput neuromodulator screenings, surface marker screening, organ bath assays and data analysis, CRISPR/Cas9 mediated genetic KO, bulk RNA-seq data analysis, MEA assays, histological analyses, immunofluorescence imaging, flow cytometry, drug target prediction analysis, snRNA-seq sample preparation.

R.M.S. scRNA-seq and snRNA-seq data curation and formal analysis, writing of the manuscript J.T.R. maintenance and directed differentiation of hPSCs, histological analyses, immunofluorescence imaging, flow cytometry, snRNA-seq sample preparation A.K. maintenance and directed differentiation of hPSCs, flow cytometry, organ bath assays and data analysis K.B. maintenance and directed differentiation of hPSCs, flow cytometry, high-throughput small molecule inhibitor screening, snRNA-seq sample preparation, NO-release high-throughput neuromodulator screenings, organ bath assays and data analysis

Z.G. reporter line generation

A.K.C. imaging, optogenetic line light-stimulation measurements

A.C. scRNA-seq and snRNA-seq data analysis

M.N.R. neuromodulator treatment and flow cytometry, manuscript proofreading

S.D. organ bath assays and data analysis

M.K. MEA assay and data analysis

J.W. Computational data analysis

R.K.S. organ bath assay data analysis

C.M. mouse surgery, tissue preparation, immunofluorescence imaging

S.B. high-content image analysis

M.G.K. scRNA-seq data analysis

J.Y. bulk- and scRNA-seq sample preparations

T.J.N. Supervision of MEA assay and data analysis

H.G. Supervision of experiments and computational data analysis by M.K., J.Y., J.W.

J.A.K. Supervision of organ bath assays and data analysis

N.T. Supervision of mouse surgery, tissue preparation, immunofluorescence imaging

F.F. design and conception of the study, supervision of all experiments, writing of the manuscript

---

**Figure S1:**
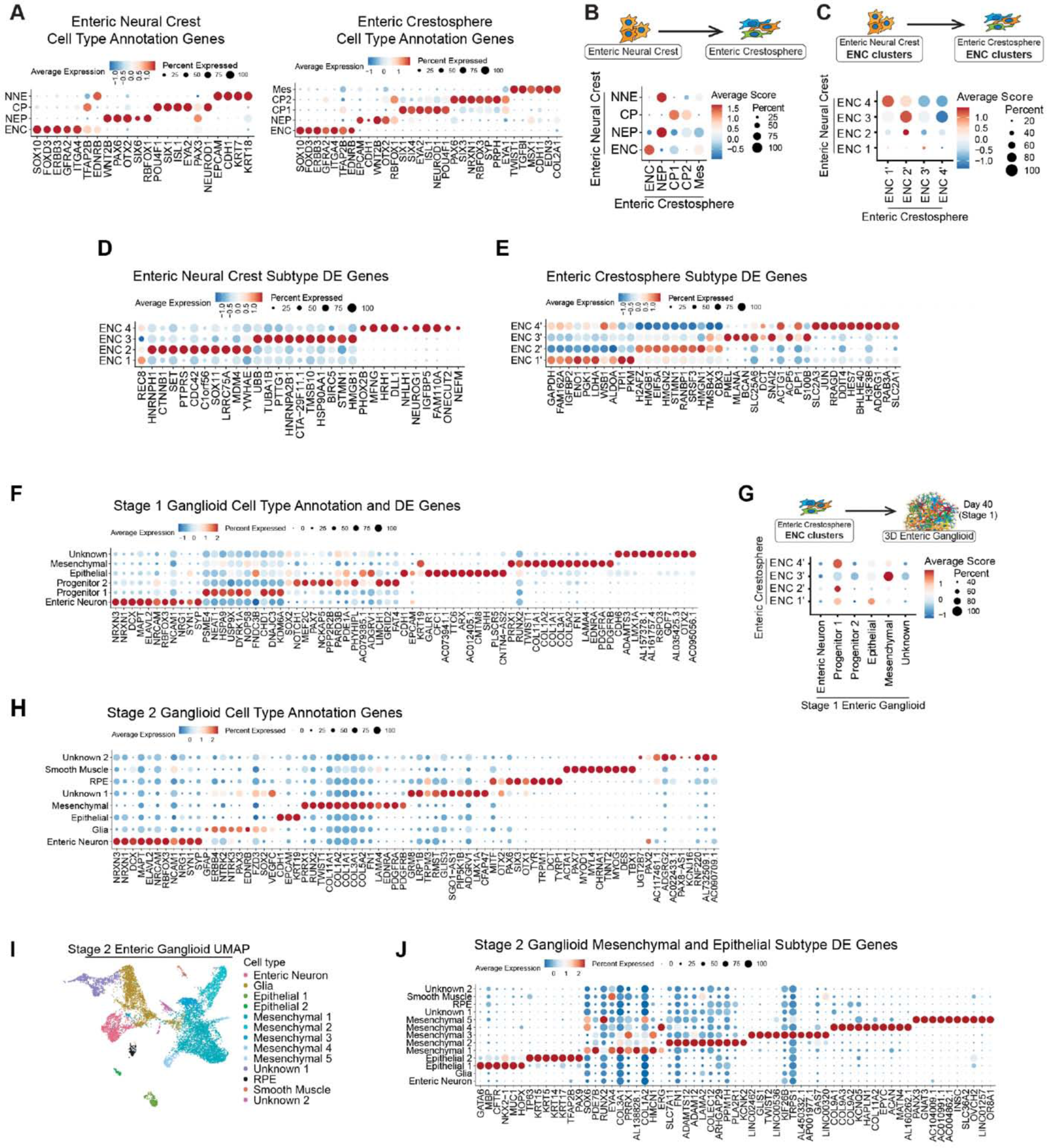
Cell-type specific molecular characterization of different developmental stages of hPSC-deried ENS models. **A)** Dot plot of the scaled average expression of cell type annotation genes for enteric neural crest (left) and enteric crestosphere (right) cell types. All data are derived from scRNA-seq analysis. **B)** Dot plot of the average module scores of enteric neural crest cell type transcriptional signatures in enteric crestosphere cell types. All data are derived from scRNA-seq analysis. **C)** Dot plot of the average module scores of enteric neural crest cells (D10) subtype transcriptional signatures in enteric crestosphere (D15) subtypes. **D-F**) Dot plot of the scaled average expression of the top 10 differentially expressed genes for each enteric neural crest (D10, **D**), enteric crestosphere (D15, **E**), stage 1 enteric ganglioid (**F**) cell type (Unknown clusters showing top 10 differentially expressed genes). **G)** Dot plot of the average module scores of enteric crestosphere (D15) subtype transcriptional signatures in stage 1 enteric ganglioid cell types. **H)** Dot plot of the scaled average expression of the top 10 differentially expressed genes for stage 2 enteric ganglioid cell types. **I)** UMAP of epithelial and mesenchymal subtypes present in stage 2 enteric ganglioid cultures. **J)** Dot plot of the scaled average expression of the 10 ten differentially expressed genes of stage 2 enteric ganglioid epithelial and mesenchymal subtypes.

**Figure S2:**
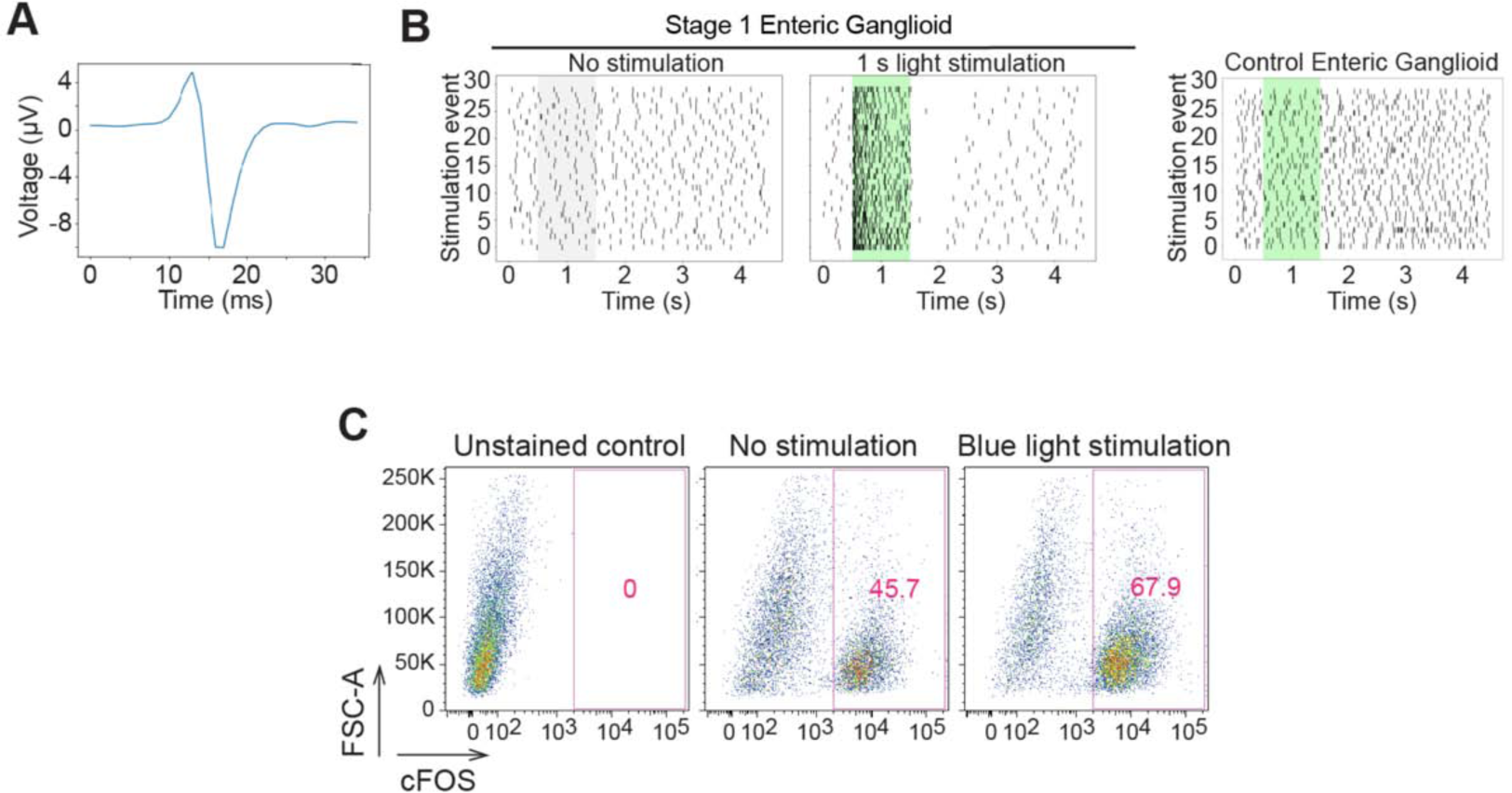
hPSC-derived enteric ganglioids comprise functionally active neurons. **A)** Representative diagram of a spontaneous neuronal firing recorded during multi-electrode array analysis (MEA) of stage 1 enteric ganglioids. **B)** MEA analysis of baseline and blue light-stimulated neuronal activities in stage 1 hSYN-ChR2-EYFP (left) and control (right) enteric ganglioids. **C)** Flow cytometry analysis of cFOS expression in hSYN-ChR2-EYFP-derived stage 2 enteric ganglioids in response to blue light stimulation.

**Figure S3:**
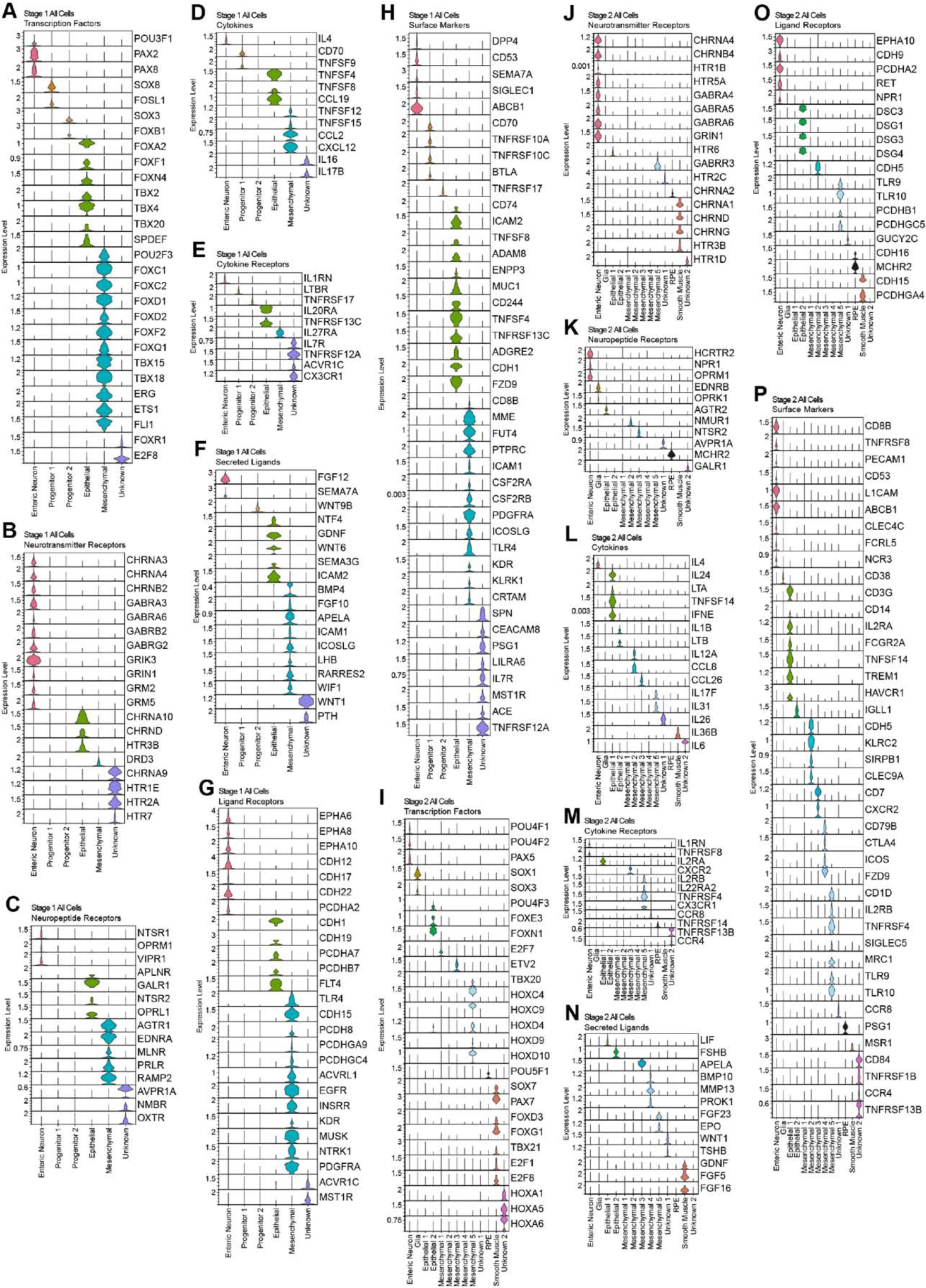
Identification of cluster specific markers by gene category in stage 1 and 2 enteric ganglioids. **A-H**) Violin plot stack of cell-type specific transcription factors (**A**), neurotransmitter receptors (**B**), neuropeptide receptors (**C**), cytokines (**D**), cytokine receptors (**E**), secreted ligands (**F**), ligand receptors (**G**), surface markers (**H**), in stage 1 enteric ganglioids. **I-P**) Violin plot stack of cell-type specific transcription factors (**I**), neurotransmitter receptors (**J**), neuropeptide receptors (**K**), cytokines (**L**), cytokine receptors (**M**), secreted ligands (**N**), ligand receptors (**O**) and surface markers (**P**) in stage 2 enteric ganglioids.

**Figure S4:**
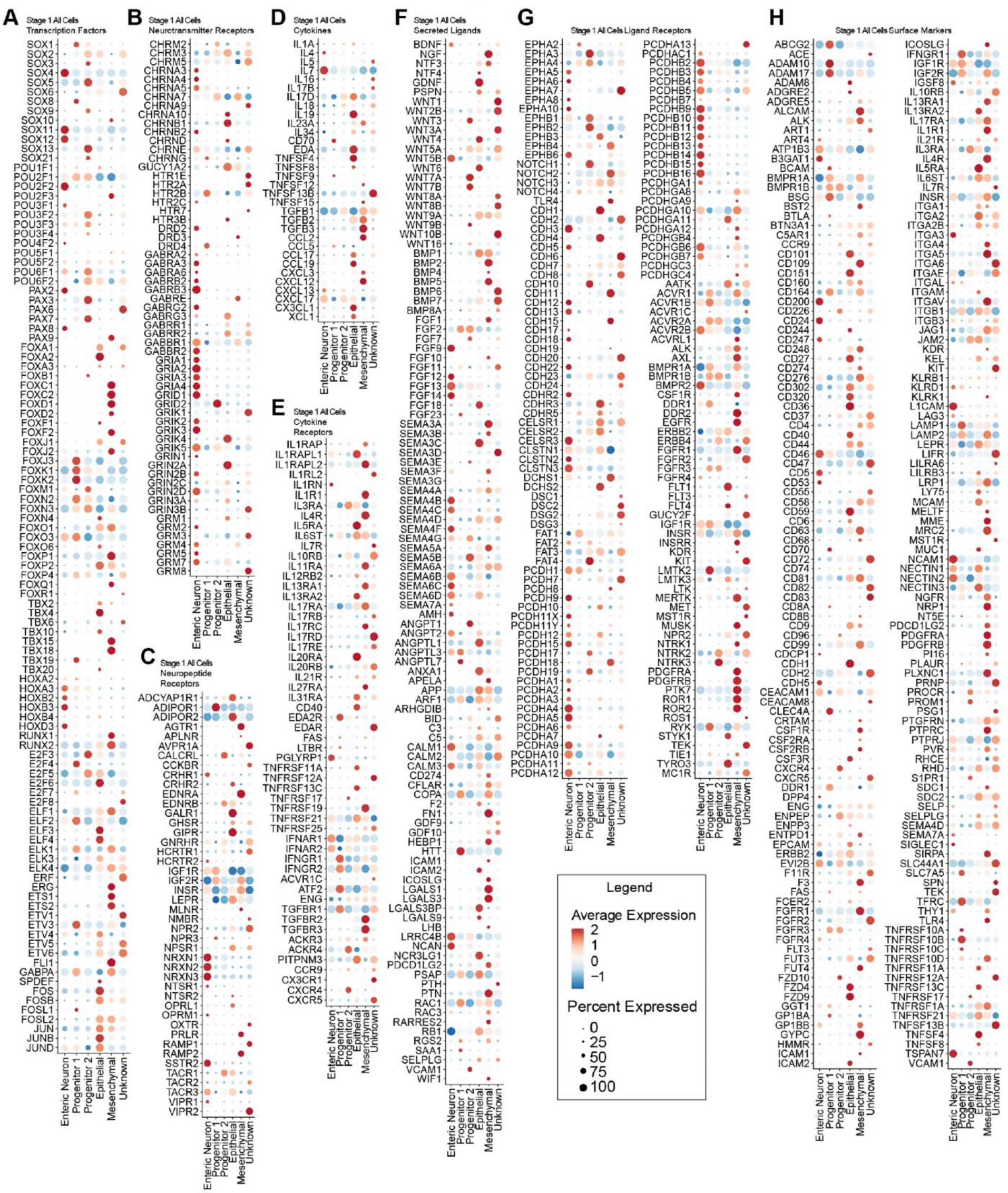
Expression profiles of selected gene categories in stage 1 enteric ganglioid cell types. **A-H**) Dot plot of the scaled average expression of selected transcription factor families (**A**), neurotransmitter receptors (**B**), neuropeptide receptors (**C**), cytokines (**D**), cytokine receptors (**E**), selected secreted ligands (**F**), selected ligand receptors (**G**) and surface markers (**H**) in stage 1 enteric ganglioid cell types.

**Figure S5:**
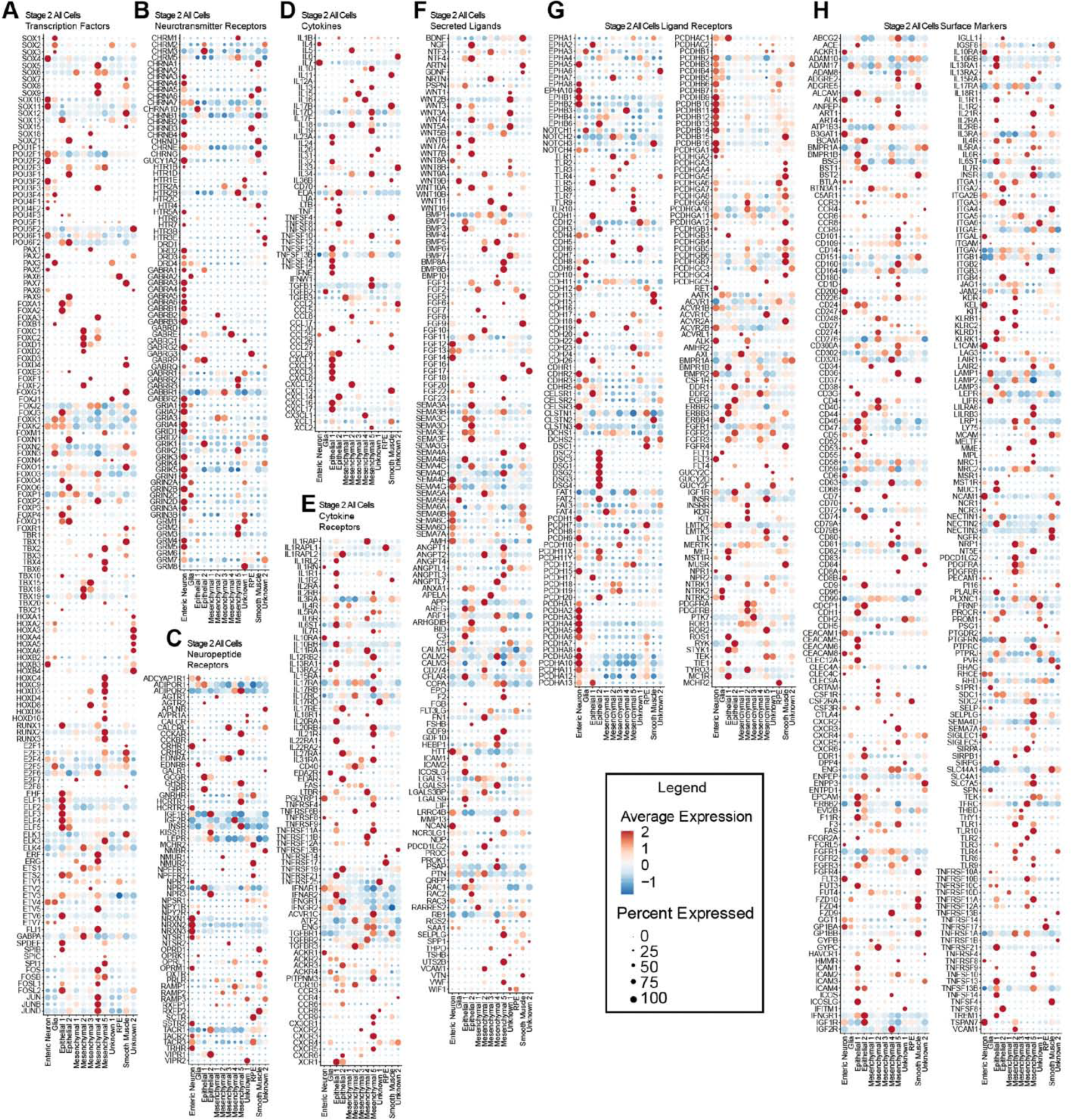
Expression profiles of selected gene categories in stage 2 enteric ganglioid cell types. **A-H**) Dot plot of the scaled average expression of selected transcription factor families (**A**), neurotransmitter receptors (**B**), neuropeptide receptors (**C**), cytokines (**D**), cytokine receptors (**E**), selected secreted ligands (**F**), selected ligand receptors (**G**) and surface markers (**H**) in stage 2 enteric ganglioid cell types.

**Figure S6:**
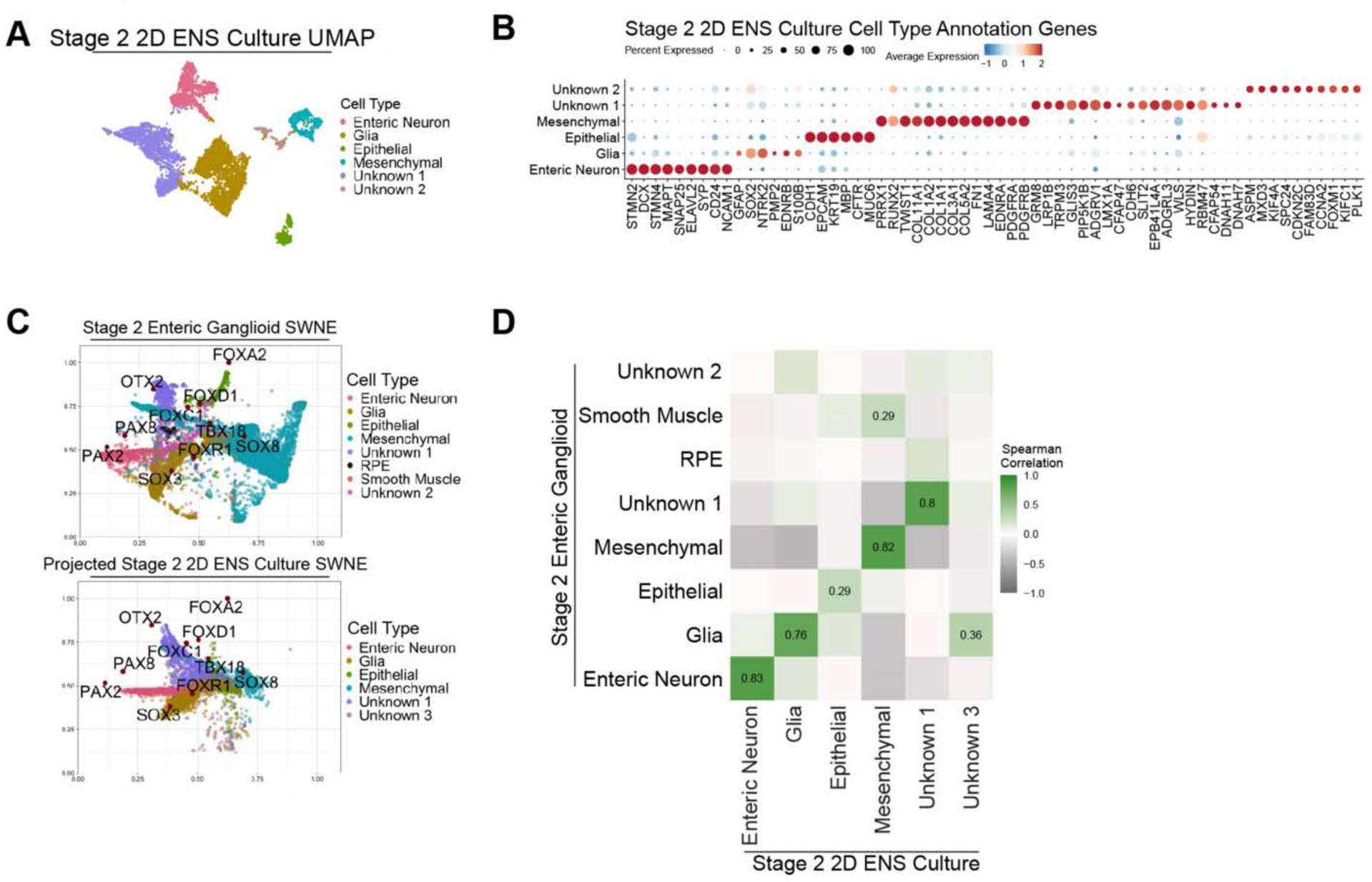
Comparison of stage 2 enteric ganglioid and 2D ENS cultures. **A)** UMAP of cell types present in stage 2 2D ENS cultures. **B)** Dot plot of the scaled average expression of cell type annotation genes for each stage 2 2D ENS culture cell type (unknown clusters showing the top 10 differentially expressed genes). **C)** Projection of stage 2 2D ENS culture cell types (bottom) onto the SWNE of stage 2 enteric ganglioid cells with overlayed projection of stage 1 cell-type specific transcription factors from **Figure S3**. **D)** Heatmap matrix of Spearman correlations based on scaled expression of 3000 anchor features shared significantly variable genes (or anchor features) between stage 2 2D ENS cultures (x-axis) and enteric ganglioids (y-axis).

**Figure S7:**
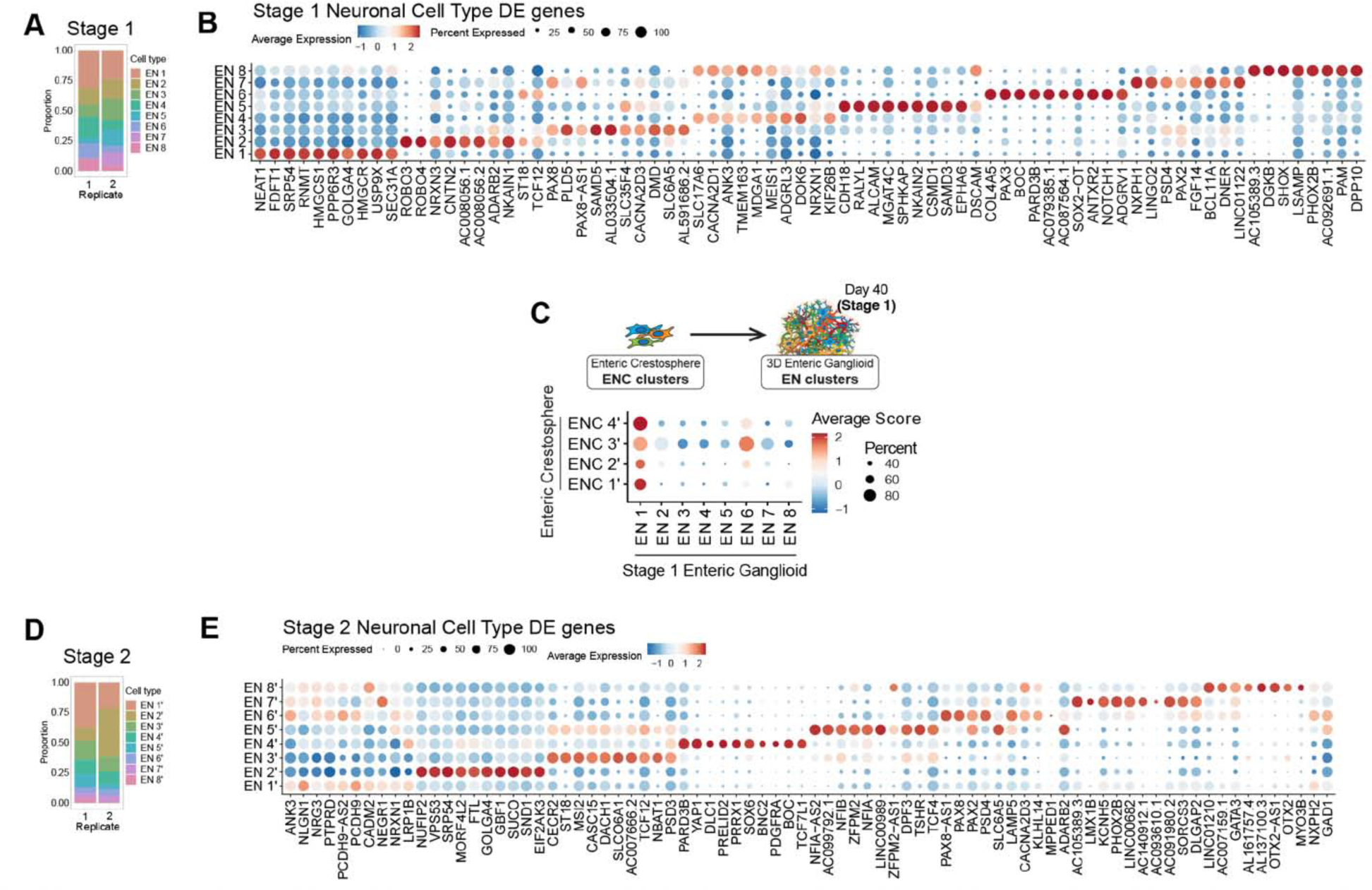
Profiling of enteric neuron subtypes present in stage 1 and 2 enteric ganglioids. **A)** Distribution of enteric neuron subtypes in biological replicates of stage 1 enteric ganglioid cultures. **B)** Dot plot of the scaled average expression of the top 10 differentially expressed genes for each stage 1 enteric ganglioid neuronal subtype. **C)** Dot plot of the average module scores of enteric crestosphere (D15) subtype transcriptional signatures in stage 1 enteric ganglioid neuronal subtypes. **D)** Distribution of enteric neuron subtypes in biological replicates of stage 2 enteric ganglioid cultures. **E)** Dot plot of the scaled average expression of the top 10 differentially expressed genes for each stage 2 enteric ganglioid neuronal subtype.

**Figure S8:**
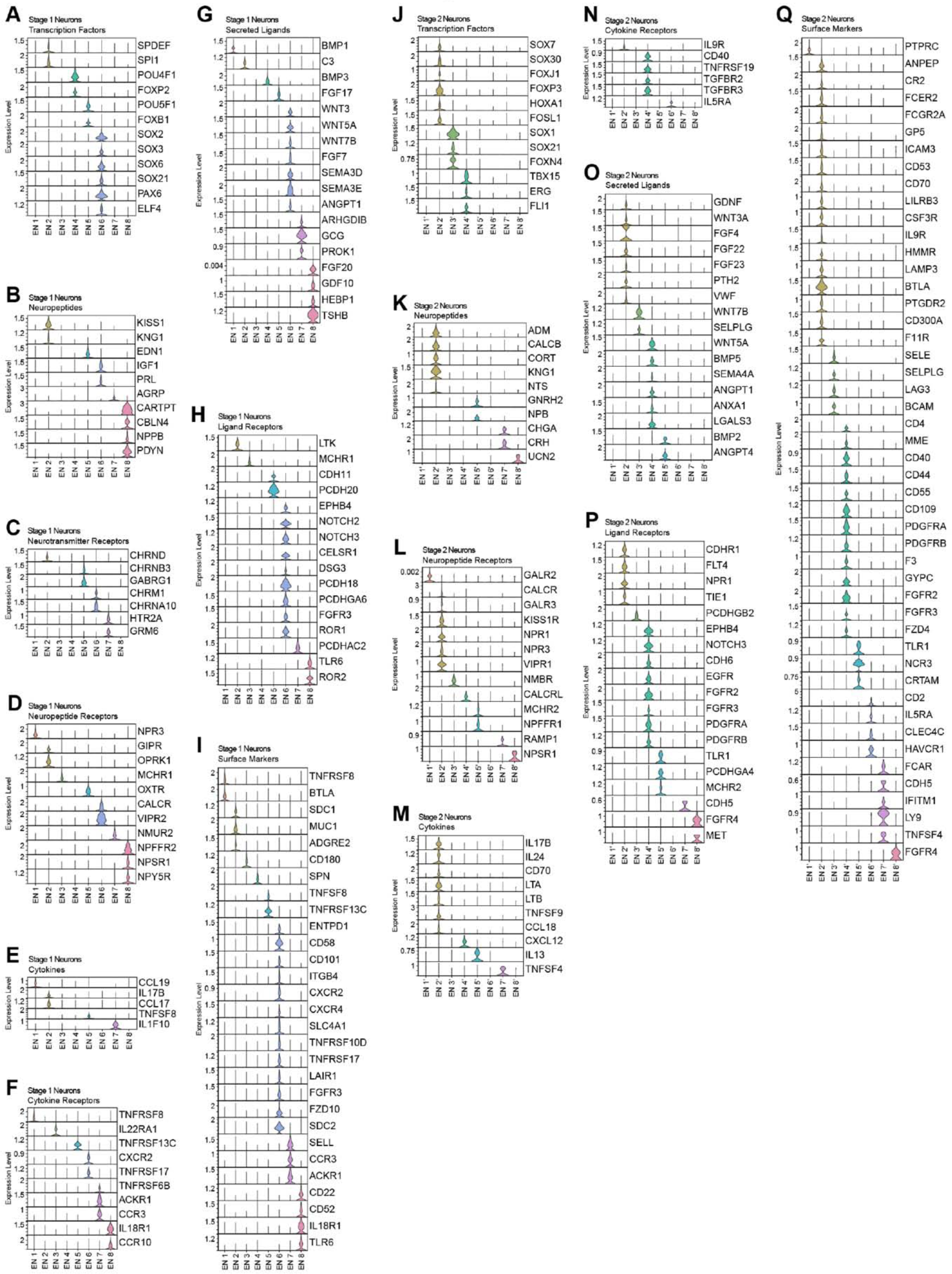
Identification of cluster specific markers by gene category in stage 1 and 2 enteric ganglioid neuronal subtypes. **A-I**) Violin plot stack of cell-type specific transcription factors (**A**), neuropeptides (**B**), neurotransmitter receptors (**C**), neuropeptide receptors (**D**), cytokines (**E**), cytokine receptors (**F**), secreted ligands (**G**), ligand receptors (**H**) and surface markers (**I**) in stage 1 enteric ganglioid neuronal subtypes. **J-R**) Violin plot stack of cell-type specific transcription factors (**J**), neuropeptides (**K**), neurotransmitter receptors (**L**), neuropeptide receptors (**M**), cytokines (**N**), cytokine receptors (**O**), secreted ligands (**P**), ligand receptors (**Q**) and surface markers (**R**) in stage 2 enteric ganglioid neuronal subtypes.

**Figure S9:**
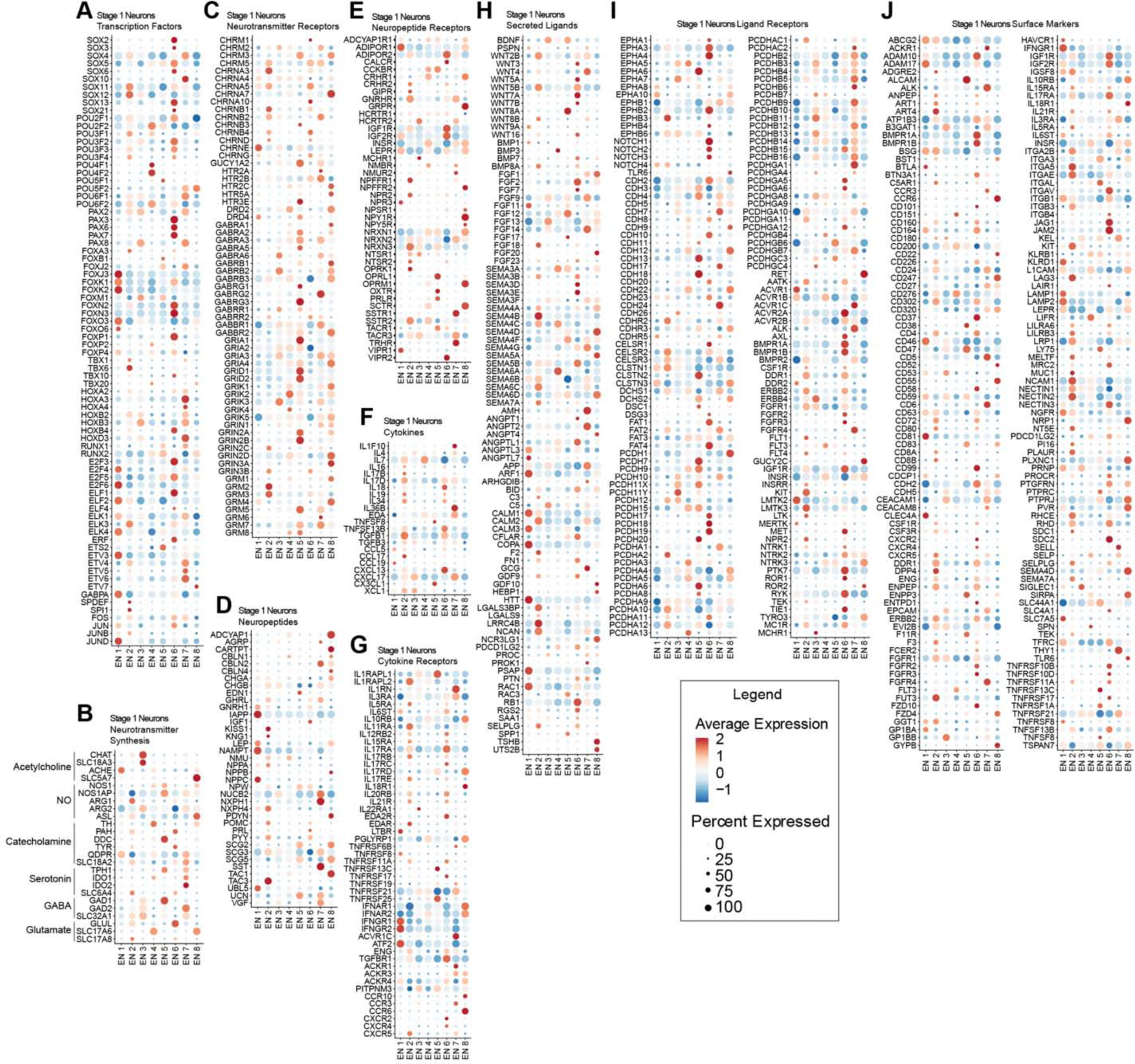
Expression profiles of selected gene categories in stage 1 enteric ganglioid neuronal subtypes. **A-J**) Dot plot of the scaled average expression of selected transcription factor families (**A**), neurotransmitter synthesis genes (**B**), neuropeptide (**C**), neurotransmitter receptors (**D**), neuropeptide receptors (**E**), cytokines (**F**), cytokine receptors (**G**), selected secreted ligands (**H**), selected ligand receptors (**I**) and surface markers (**J**) in stage 1 enteric ganglioid neuronal subtypes.

**Figure S10:**
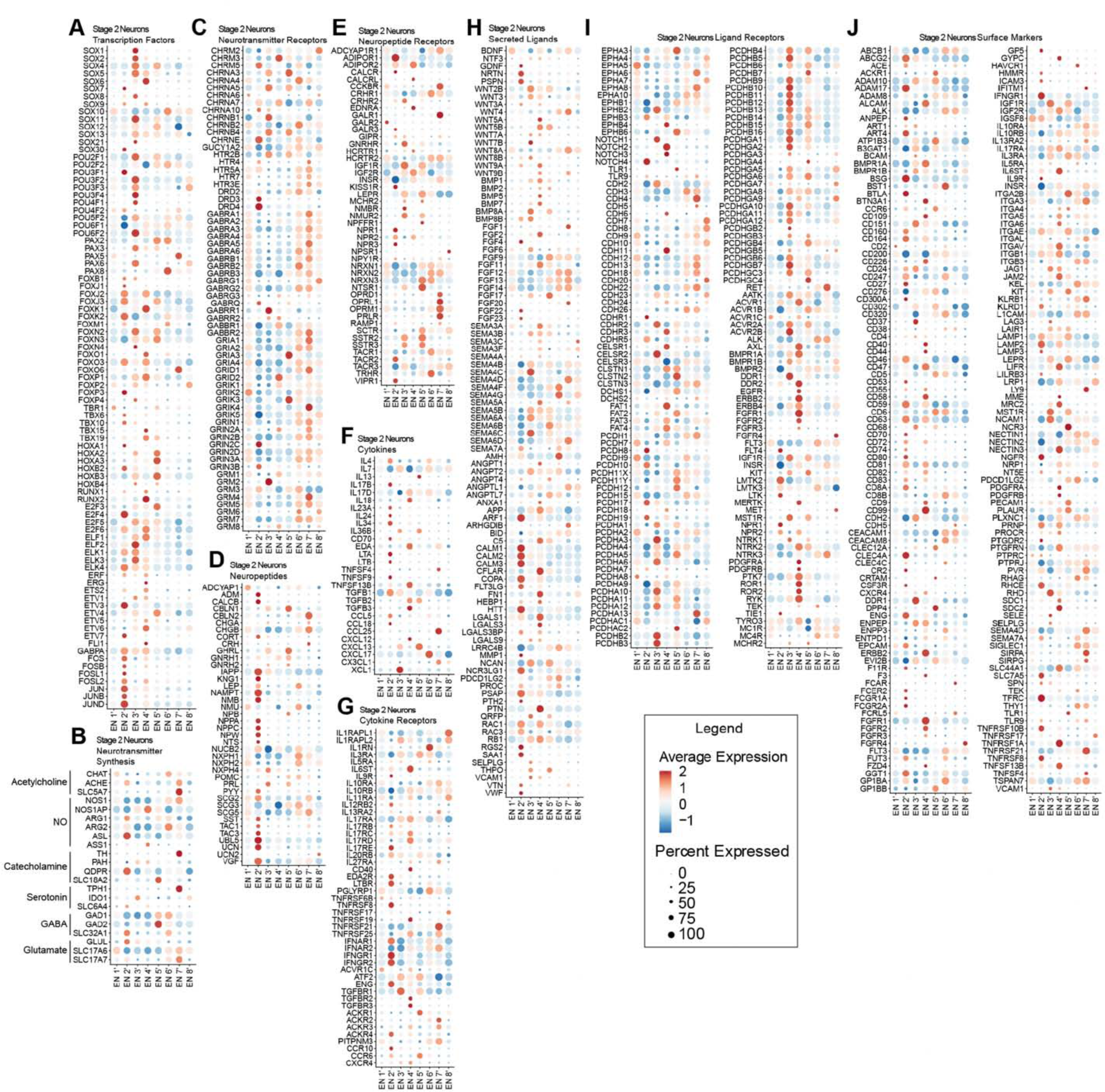
Expression profiles of selected gene categories in stage 2 enteric ganglioid neuronal subtypes. **A-J**) Dot plot of the scaled average expression of selected transcription factor families (**A**), neurotransmitter synthesis genes (**B**), neuropeptide (**C**), neurotransmitter receptors (**D**), neuropeptide receptors (**E**), cytokines (**F**), cytokine receptors (**G**), selected secreted ligands (**H**), selected ligand receptors (**I**) and surface markers (**J**) in stage 2 enteric ganglioid neuronal subtypes.

**Figure S11:**
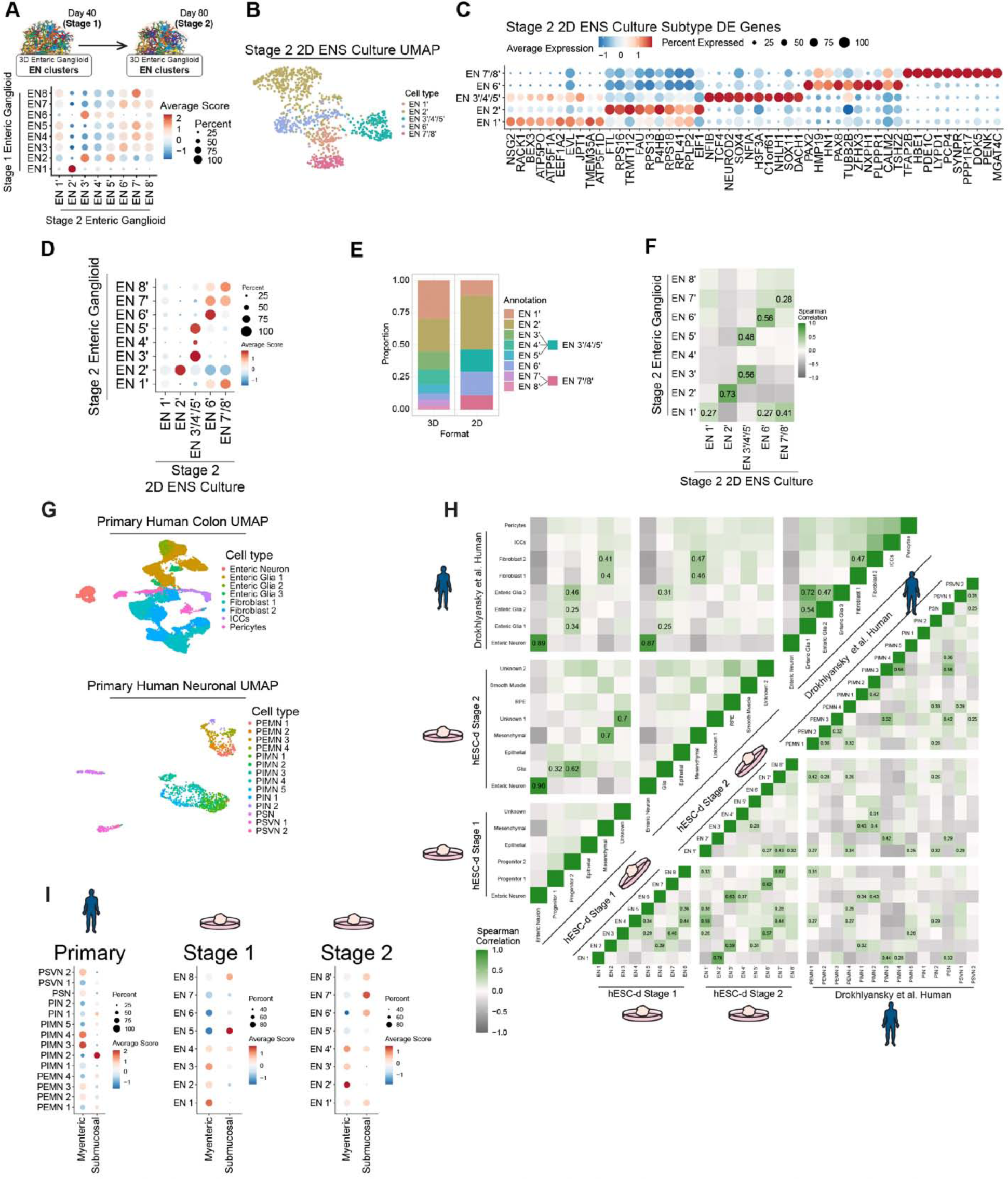
Comparative analysis of hPSC-derived ENS culture neurons and primary enteric datasets. **A)** Dot plot of the average module scores of stage 1 enteric ganglioid neuronal subtype transcriptional signatures in stage 2 enteric ganglioid neuronal subtypes. **B)** UMAP of stage 2 2D ENS culture neuronal subtypes. **C)** Dot plot of the scaled average expression of the top 10 differentially expressed genes for each stage 2 2D ENS culture neuron subtype. **D)** Dot plot of the average module scores of stage 2 enteric ganglioid neuron subtype transcriptional signatures in stage 2 2D ENS culture neuron subtypes. **E)** Comparison of the distribution of enteric neuron subtypes in 2D versus 3D enteric neuron cultures. **F)** Heatmap matrix of Spearman correlations based on scaled expression of 3000 anchor features shared significantly variable genes (or anchor features) between stage 2 2D ENS cultures (x-axis) and ganglioids (y-axis). **G)** UMAPs of cell types (top) and neuronal subtypes (bottom) present in a primary adult human colon dataset. **H)** Heatmap matrix of Spearman correlations based on scaled expression of 3000 anchor features shared significantly variable genes (or anchor features) between stage 1 and 2 ganglioid neuron subtypes, and adult human enteric neuron subtypes. **I)** Dot plot of the average module scores for myenteric and submucosal neuron transcriptional signatures in adult human enteric neuron subtypes (left) and stage 1 (middle) and stage 2 (right) ganglioid neuronal subtypes.

**Figure S12:**
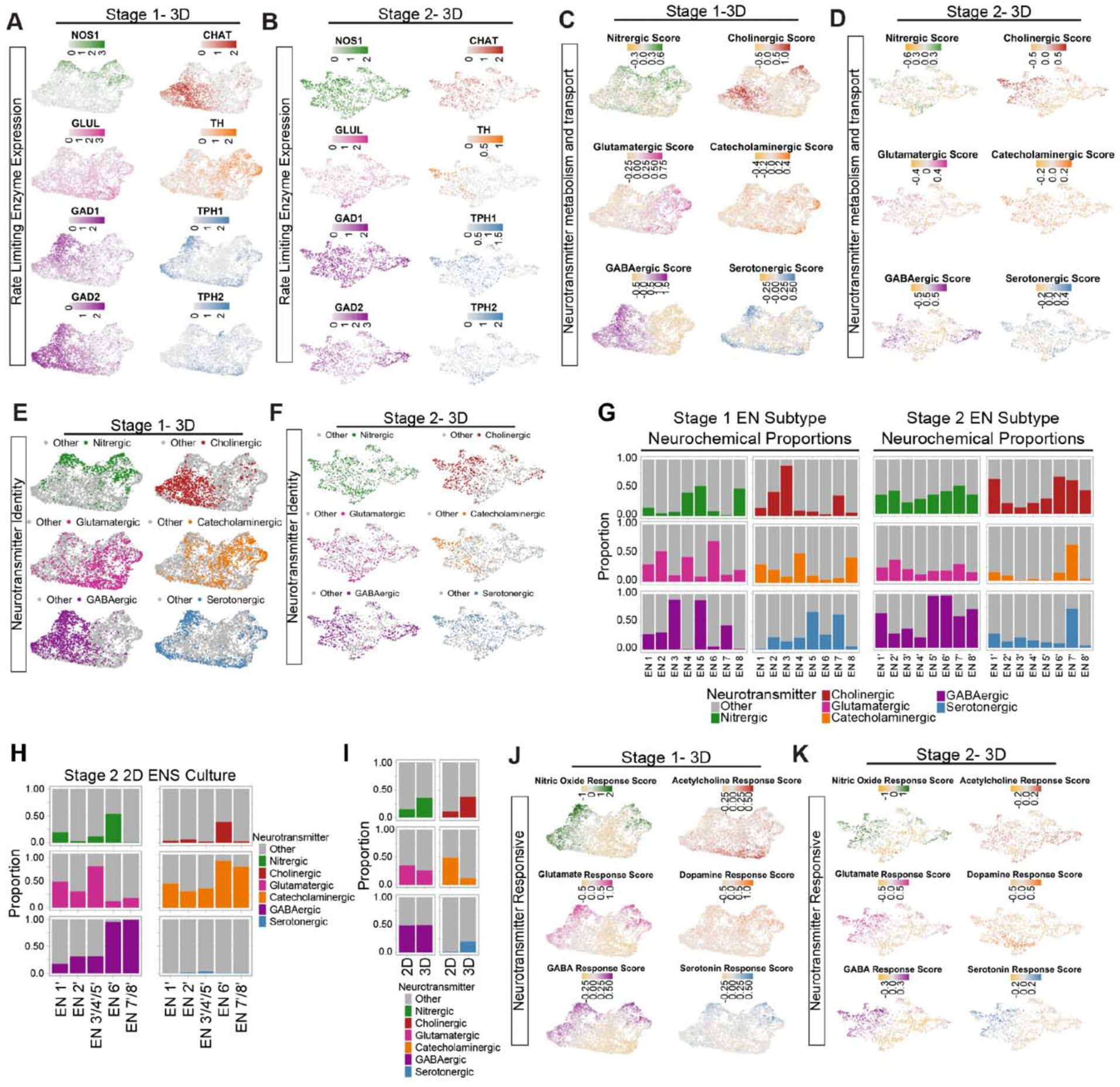
Identification of neurotransmitter producing and responsive neurons in stage 1 and stage 2 enteric ganglioid neurons. A and B) Feature plots showing the expression of rate limiting enzymes in neurotransmitter synthesis pathways by stage 1 (A) and stage 2 (B) enteric ganglioid neurons. C and D) Feature plots showing the identity score of neurotransmitters by stage 1 (C) and stage 2 (D) enteric ganglioid neurons by module scoring of genes related to each neurotransmitter’s synthesis, metabolism and reuptake. E and F) UMAP of predicted neurotransmitter producing neuron identities in stage 1 (A) and stage 2 (B) enteric ganglioids. G) Distribution of neurochemical identities in stage 1 (left) and stage 2 (right) enteric ganglioid neuron subtypes. H) Distribution of neurochemical identities in stage 2 2D culture enteric neuron subtypes. I) Comparison of the distribution of neurochemical identities in 2D culture versus 3D culture nteric neurons. J and K) Feature plots showing the predicted responsiveness of stage 1 (I) and stage 2 (J) enteric ganglioid neurons to each neurotransmitter by module scoring of neurotransmitter receptor gene families.

**Figure S13:**
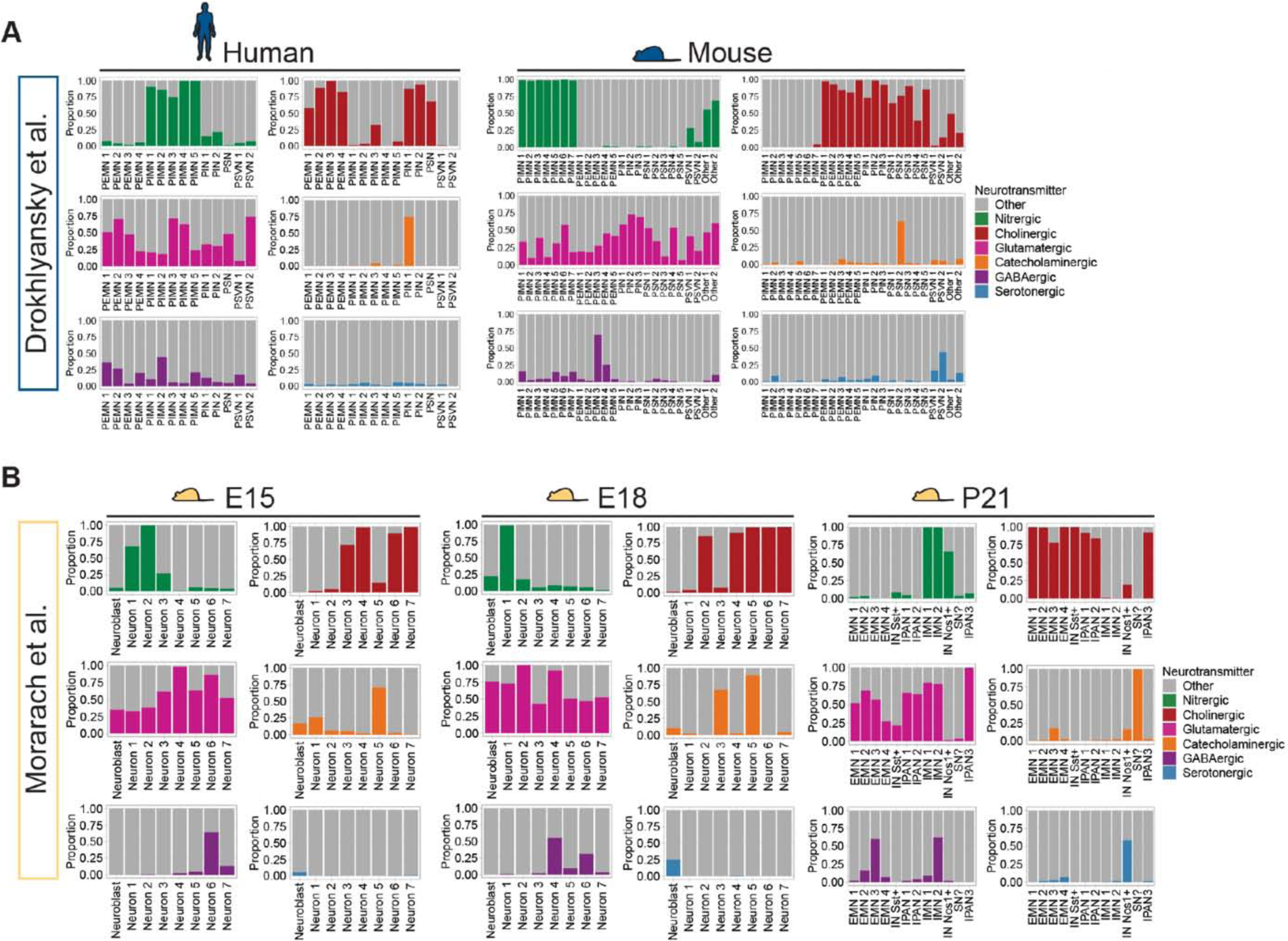
Comparison of stage 1 and 2 ganglioid cell types and subtypes to primary enteric datasets. A) Distribution of neurochemical identities in adult human (left) and adult mouse (right) enteric neuron subtypes. B) Distribution of neurochemical identities in E15 (left), E18 (middle), and P21 (right) mouse enteric neuron subtypes.

**Figure S14:**
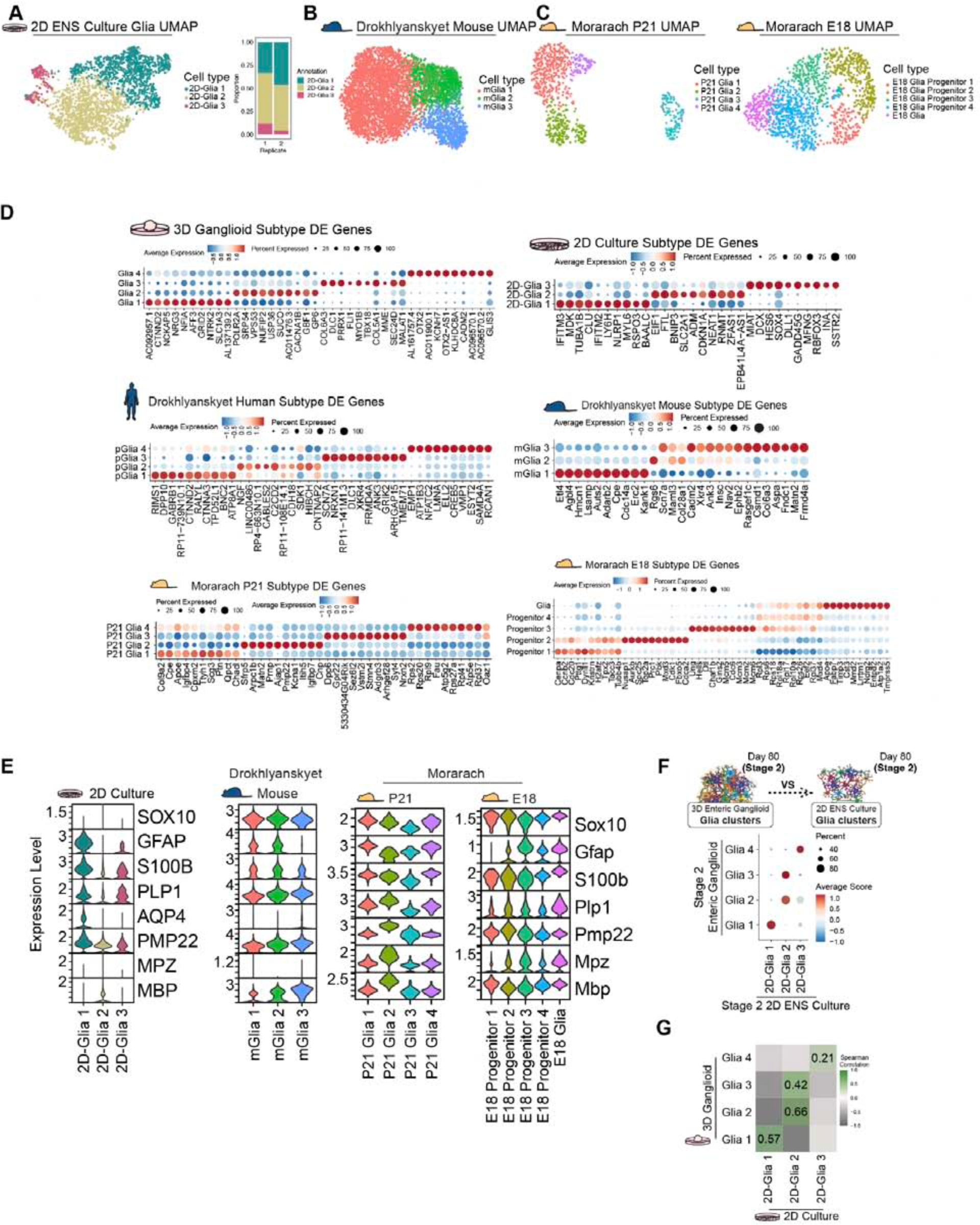
Annotation and profiling of glial subtypes in 2D ENS cultures and published primary enteric datasets. A) scRNA-seq UMAP (left) and distribution of glial subtypes in biological replicates (right) in stage 2 2D ENS cultures. B) UMAP of enteric glial subtypes present in a primary adult mouse dataset. C) UMAP of enteric glial subtypes present in a P21 (left) and enteric glia and progenitor subtypes present in an E18 (right) adult mouse dataset. D) Dot plot of the scaled average expression of the top 10 differentially expressed genes for each enteric ganglioid (top left), 2D ENS culture (top right), adult human (middle left), adult mouse (middle right), P21 mouse (bottom left) enteric glial subtypes and E18 mouse (bottom right) enteric glial and progenitor subtypes. E) Violin plot stack showing the expression of canonical glial markers in 2D ENS culture, adult mouse, and P21 and E18 mouse glial (and progenitor) subtypes. F) Dot plot of the average module scores of stage 2 enteric ganglioid glial subtype transcriptional signatures (snRNA-seq) in 2D ENS culture glial subtypes (scRNA-seq). G) Heatmap matrix of Spearman correlations based on scaled expression of 3000 anchor features shared significantly variable genes (or anchor features) between 2D ENS cultures (x-axis) and enteric ganglioids (y-axis).

**Figure S15:**
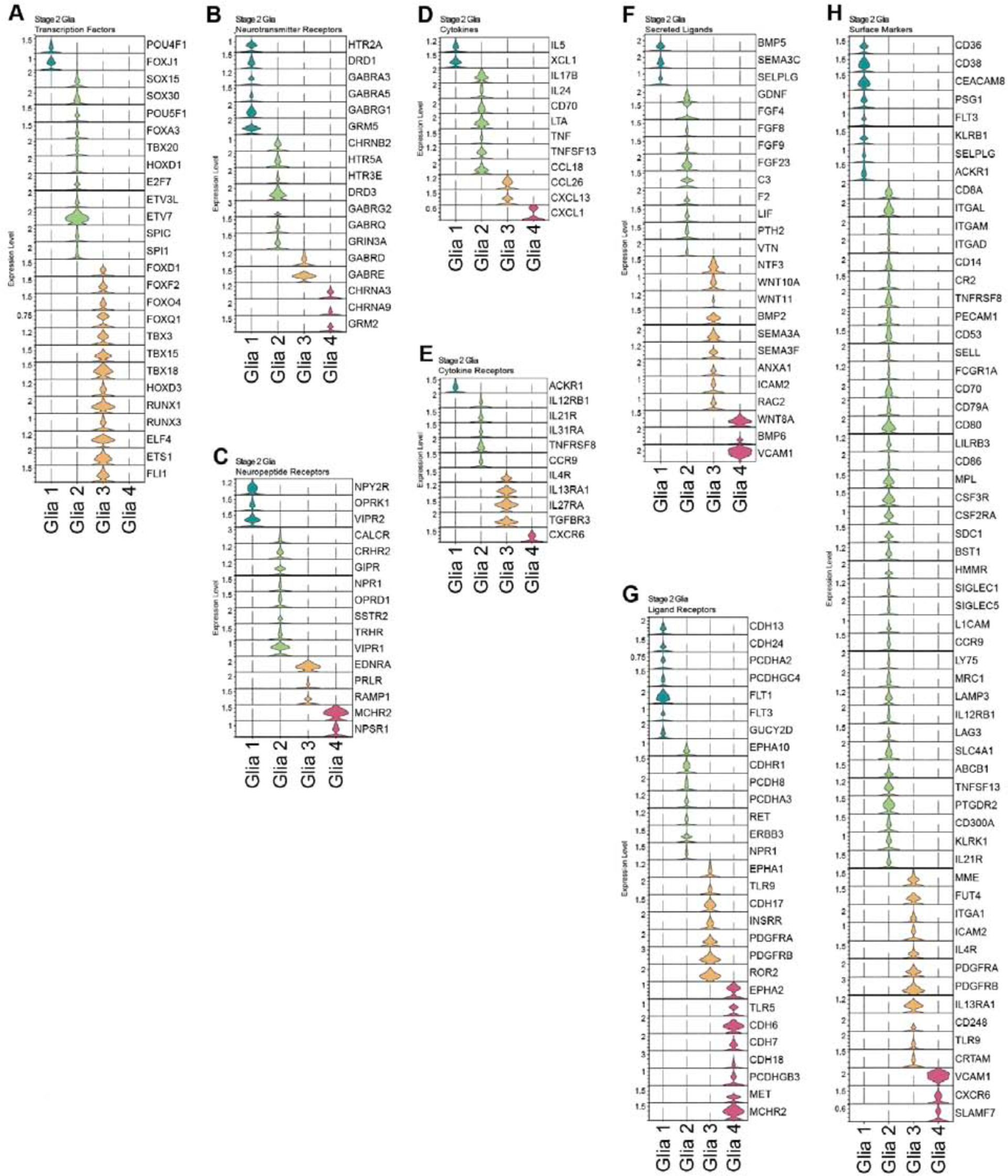
Identification of cluster specific markers by gene category in enteric ganglioid glial subtypes. **A-H**) Violin plot stack of cell-type specific transcription factors (**A**), neurotransmitter receptors (**B**), neuropeptide receptors (**C**), cytokines (**D**), cytokine receptors (**E**), selected secreted ligands (**F**), selected ligand receptors (**G**) and surface markers (**H**) in enteric ganglioid glial subtypes.

**Figure S16:**
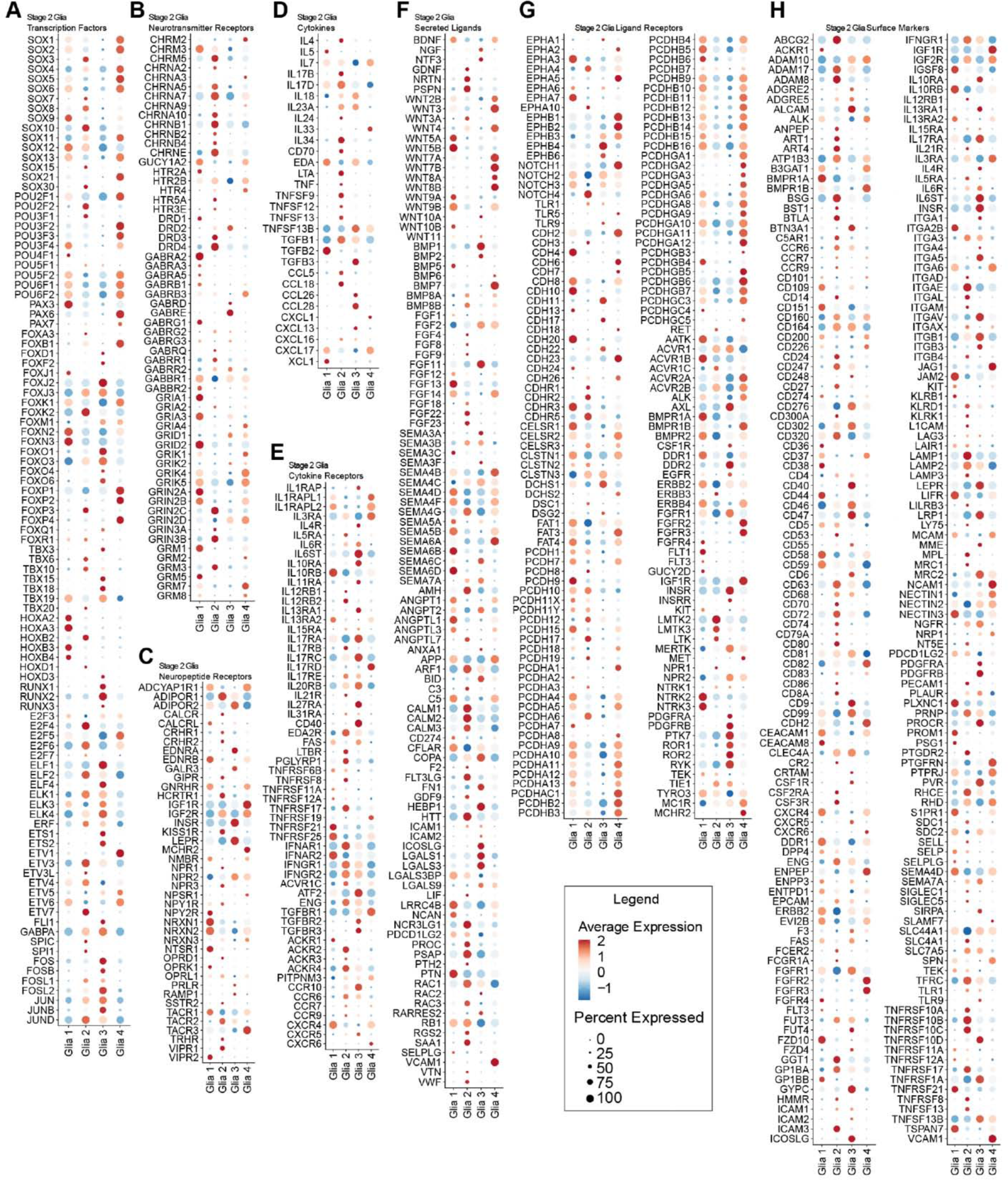
Expression profiles of selected gene categories in enteric ganglioid glial subtypes. **A-H**) Dot plot of the scaled average expression of selected transcription factor families (**A**), neurotransmitter receptors (**B**), neuropeptide receptors (**C**), cytokines (**D**), cytokine receptors (**E**), selected secreted ligands (**F**), selected ligand receptors (**G**) and surface markers (**H**) in enteric ganglioid glial subtypes.

**Figure S17:**
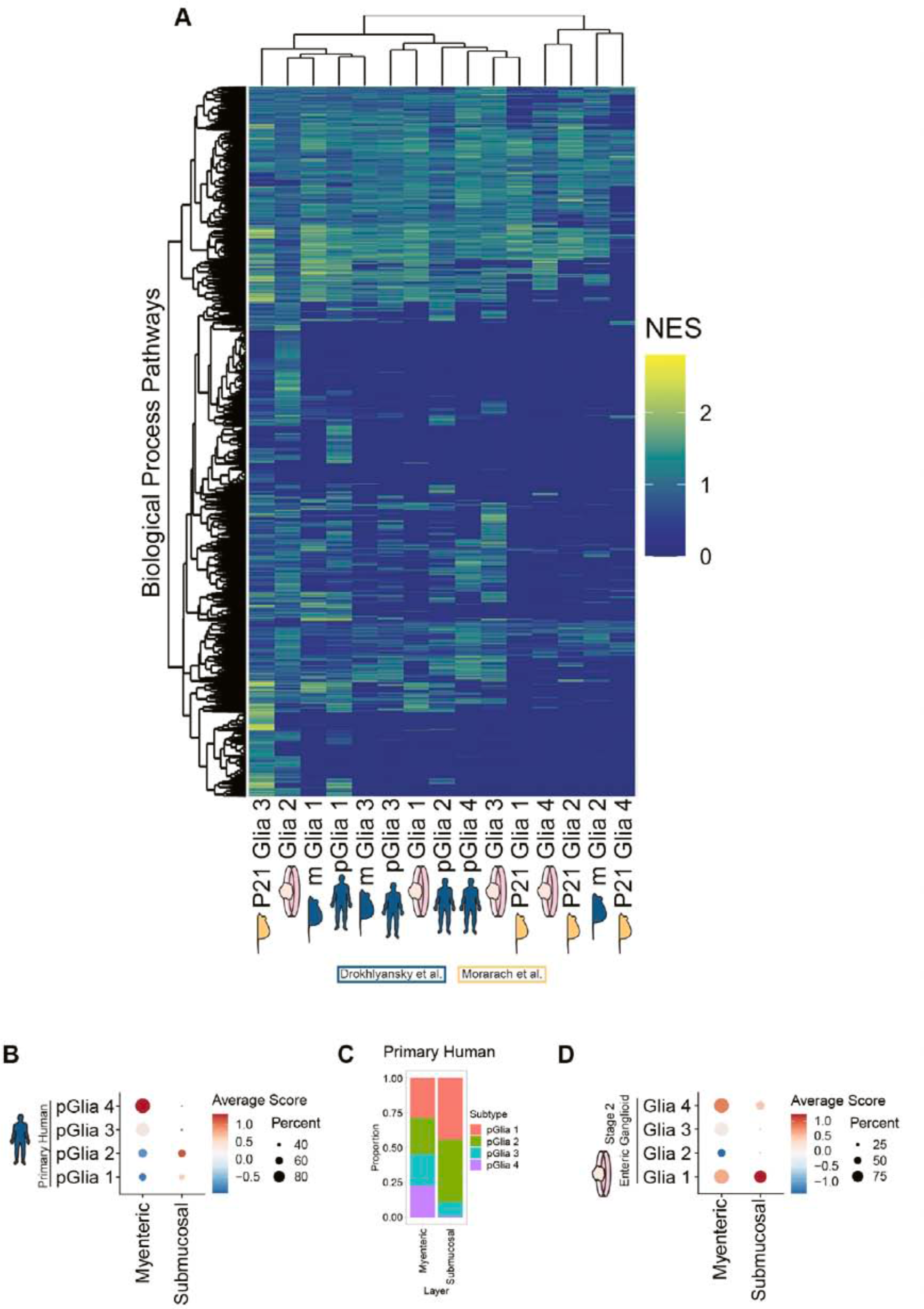
Profiling plexus identity in primary human and stage 2 enteric ganglioid glial subtypes. **A)** Hierarchical clustering of enteric ganglioid glial subtypes with primary human and mouse glial subtypes based on normalized enrichment scores of biological process gene ontology (GO) pathways. **B)** Dot plot of the average module scores for myenteric and submucosal glial transcriptional signatures in primary human enteric glial subtypes. **C)** Distribution of enteric glial subtype representation in primary human myenteric versus submucosal tissue samples. **D)** Dot plot of the average module scores for myenteric and submucosal glial transcriptional signatures in stage 2 ganglioid glial subtypes.

**Figure S18:**
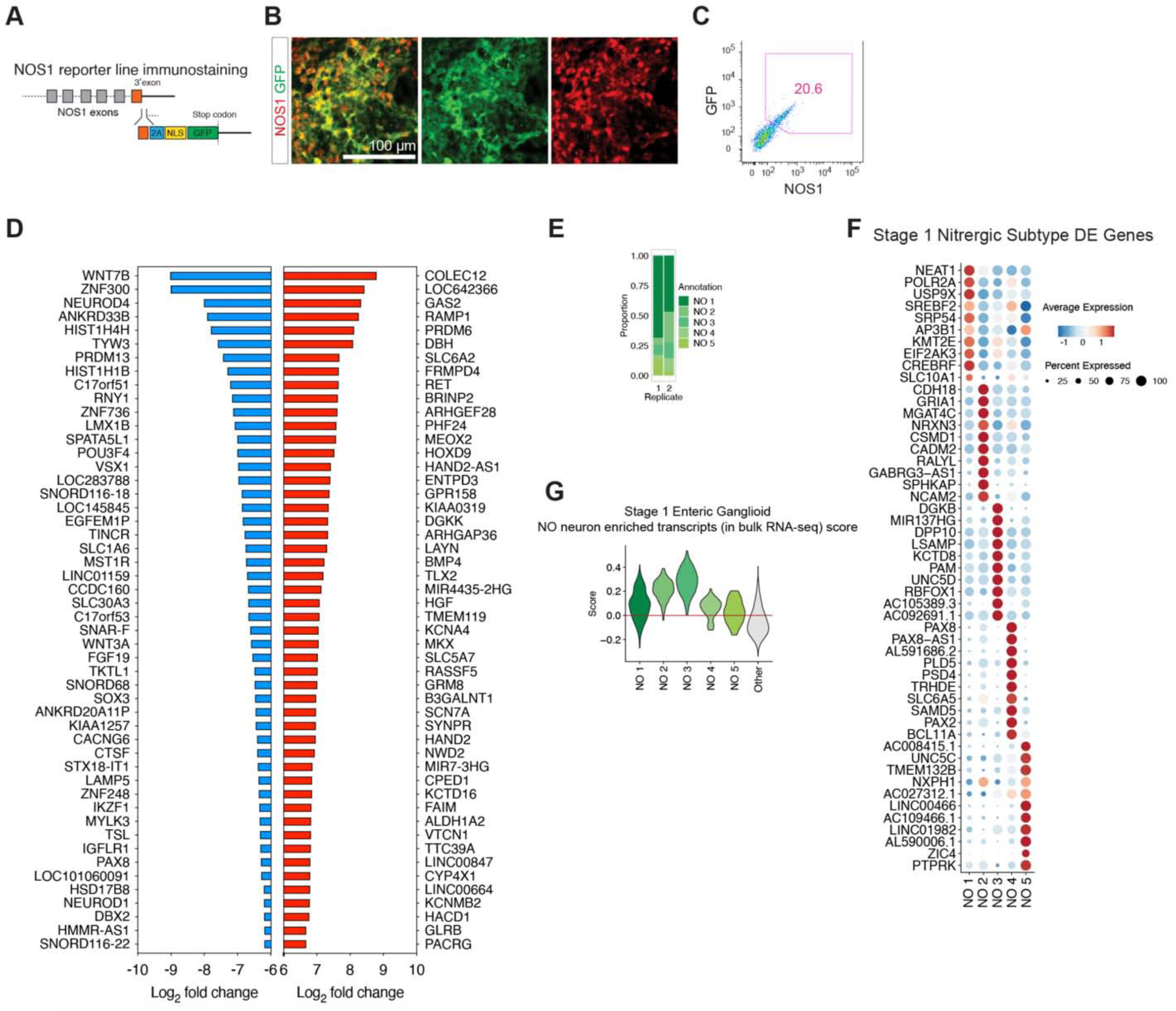
Bulk and single nuclei RNA-seq profiling of NO neurons present in stage 1 enteric ganglioids. **A)** Schematic representation of the NOS1::GFP reporter construct. **B)** Representative immunofluorescence images of a stage 2 enteric ganglioid stained for GFP and NOS1. **C)** Representative flow cytometry analysis for the expression of GFP and NOS1 in a NOS1::GFP-derived stage 1 enteric ganglioid. **D)** Bulk RNA-seq top 50 differentially expressed transcripts in FACS sorted CD24**^+^**/NOS1::GFP**^+^** cells relative to CD24**^+^**/NOS1::GFP^-^ cells. p-value < 0.05, upregulated in red, downregulated in blue. **E)** Distribution of NO neuron subtypes in biological replicates of stage 1 enteric ganglioid cultures. **F)** Dot plot of the scaled average expression of the top 10 differentially expressed genes for each stage 1 enteric ganglioid NO neuron subtype. **G)** snRNA-seq analysis violin plot of module scoring for the top 100 differentially expressed genes from CD24**^+^**/NOS1**^+^** sorted neurons versus other neurons in stage 1 enteric ganglioid NO subtypes versus other neurons.

**Figure S19:**
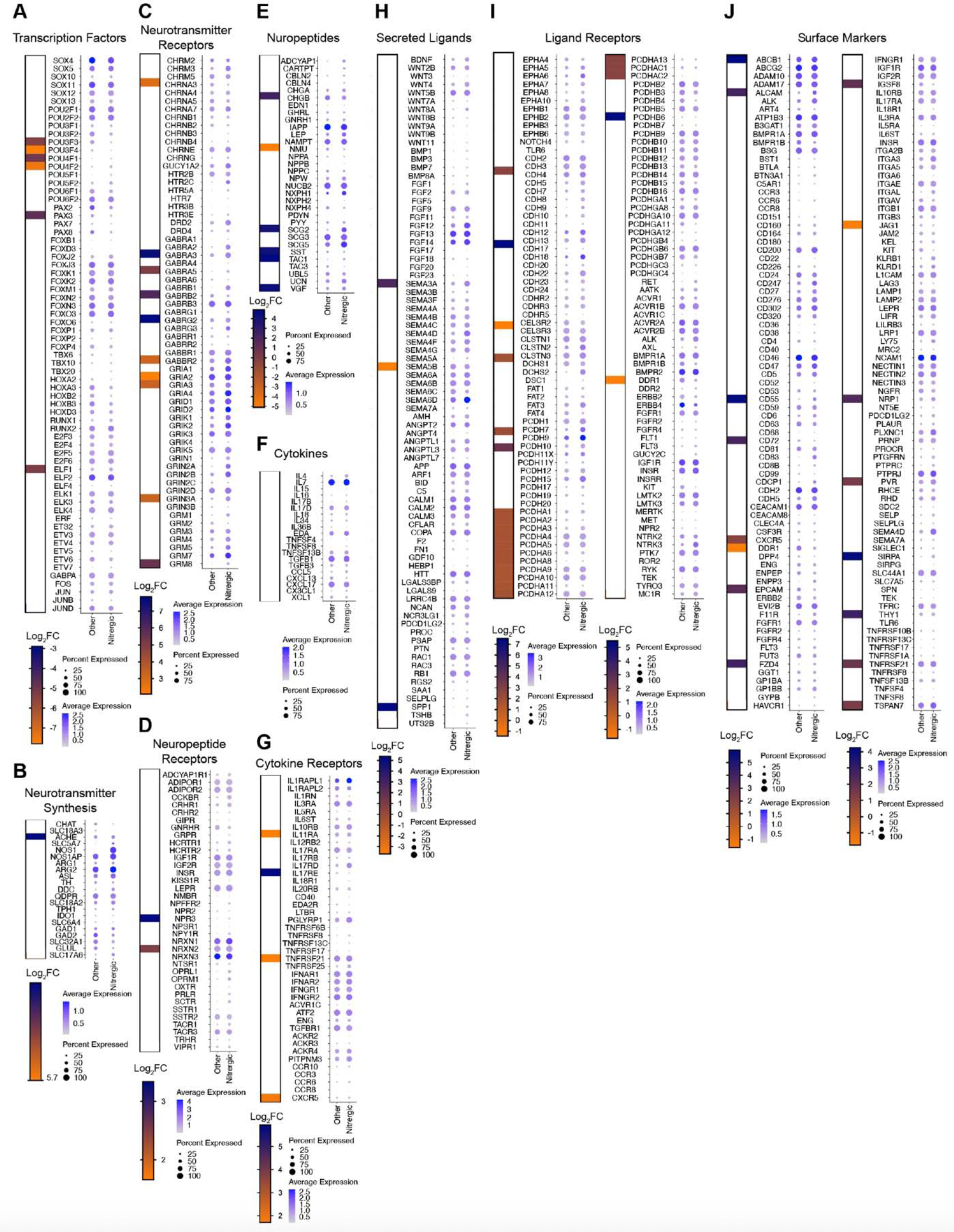
Expression profiles of selected gene categories in stage 1 enteric ganglioid NO neurons. **A-J**) Bulk RNA-seq differentially expressed (Log2FC, p-value < 0.05) genes in NOS1::GFP**^+^** neurons versus other neurons, and snRNA-seq dot plot of the average expression of selected transcription factor families (**A**), neurotransmitter synthesis genes (**B**) neurotransmitter receptors (**C**), neuropeptide receptors (**D**), neuropeptides (**E**), cytokines (**F**), cytokine receptors (**G**), selected secreted ligands (**H**), selected ligand receptors (**I**) and surface markers (**J**) in stage 1 enteric ganglioid NO neurons.

**Figure S20:**
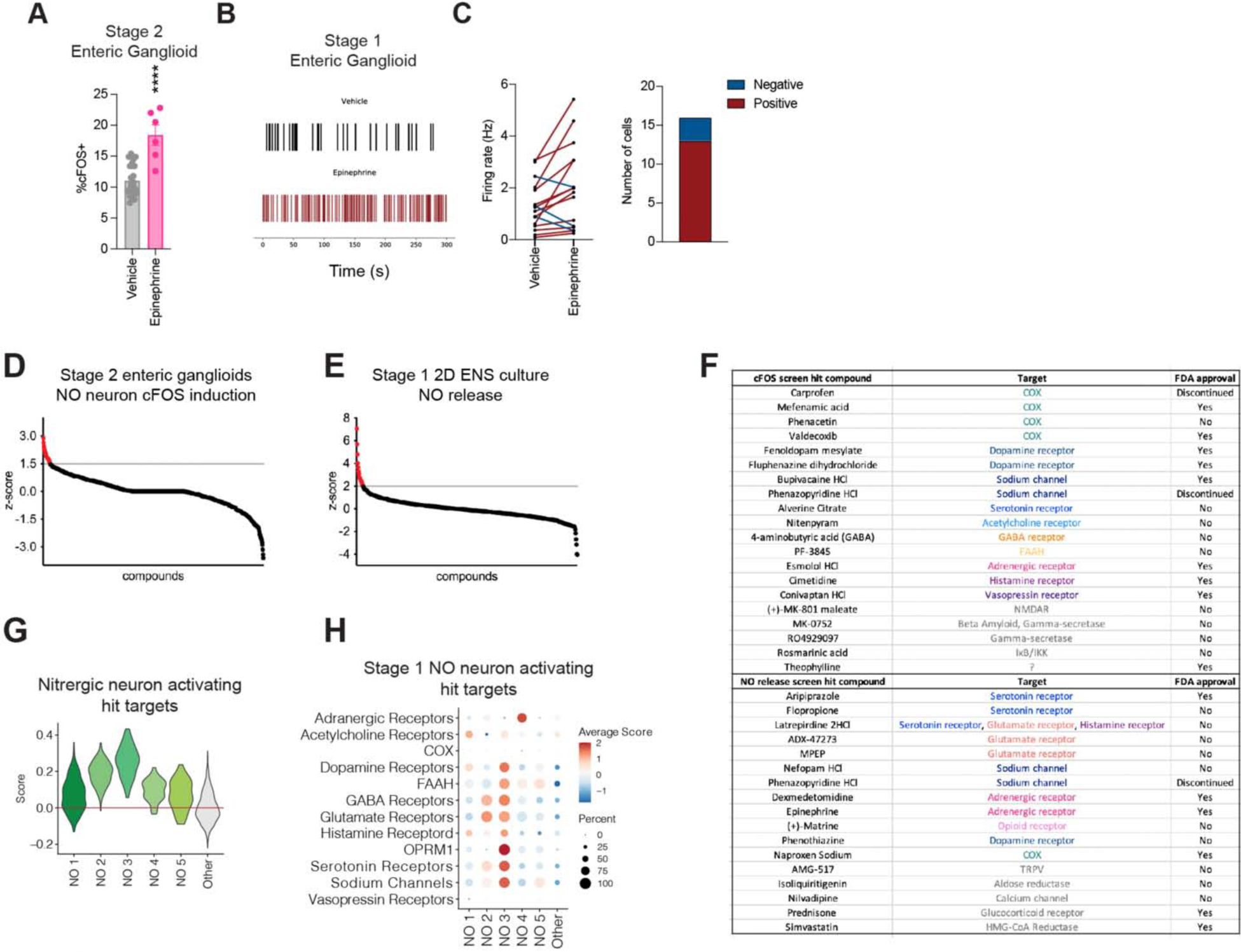
Identifying enteric NO neuron modulators by functional high-throughput screenings. **A**) Flow cytometry quantification of stage 2 enteric ganglioid cFOS expression in response to epinephrine. Mean and SEM error bars are shown, ****: p-value < 0.0001. **B and C**) Multi-electrode array (MEA) analysis (**B**) and quantification of changes in neuronal firing (**C**) in response to epinephrine in stage 1 enteric ganglioids. **D)** Identifying candidate neuromodulators that induce cFOS expression in stage 2 enteric ganglioid NO neurons by HTS. **E)** Identifying candidate neuromodulators that induce NO release in stage 1 2D ENS culture NO neurons by HTS. **F)** Predicted targets and FDA approval status of hits identified in cFOS induction and NO release screens. **G)** snRNA-seq analysis violin plot of module scoring for all predicted NO neuron activity promoting receptor genes in stage 1 enteric ganglioid NO subtypes versus other neurons. **H)** snRNA-seq analysis dot plot of the average module scores for individual categories of NO neuron activity promoting receptor gene categories.

**Figure S21:**
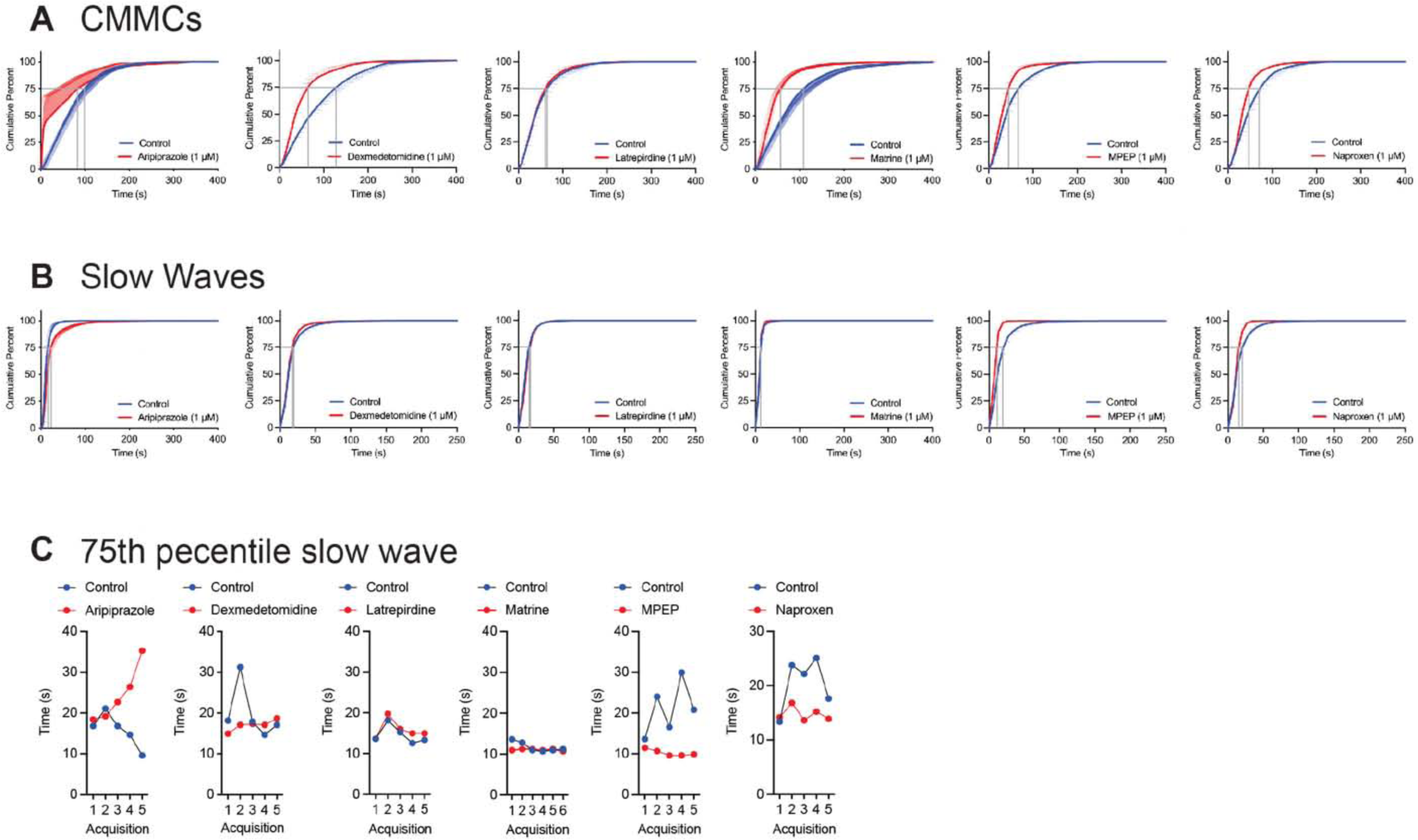
Testing selected HTS hits in *ex vivo* mouse colonic motility assays A and B) Diagrams of CMMC (A) and slow wave (B) cumulative percentiles. **C**) Quantification of latency between slow waves at 75^th^ percentile for selected HTS hits; from left to right: aripiprazole, dexmedetomidine, latrepirdine, matrine, MPEP and naproxen. One-directional SEM error bars are shown in panels **A** and **B**.

**Figure S22:**
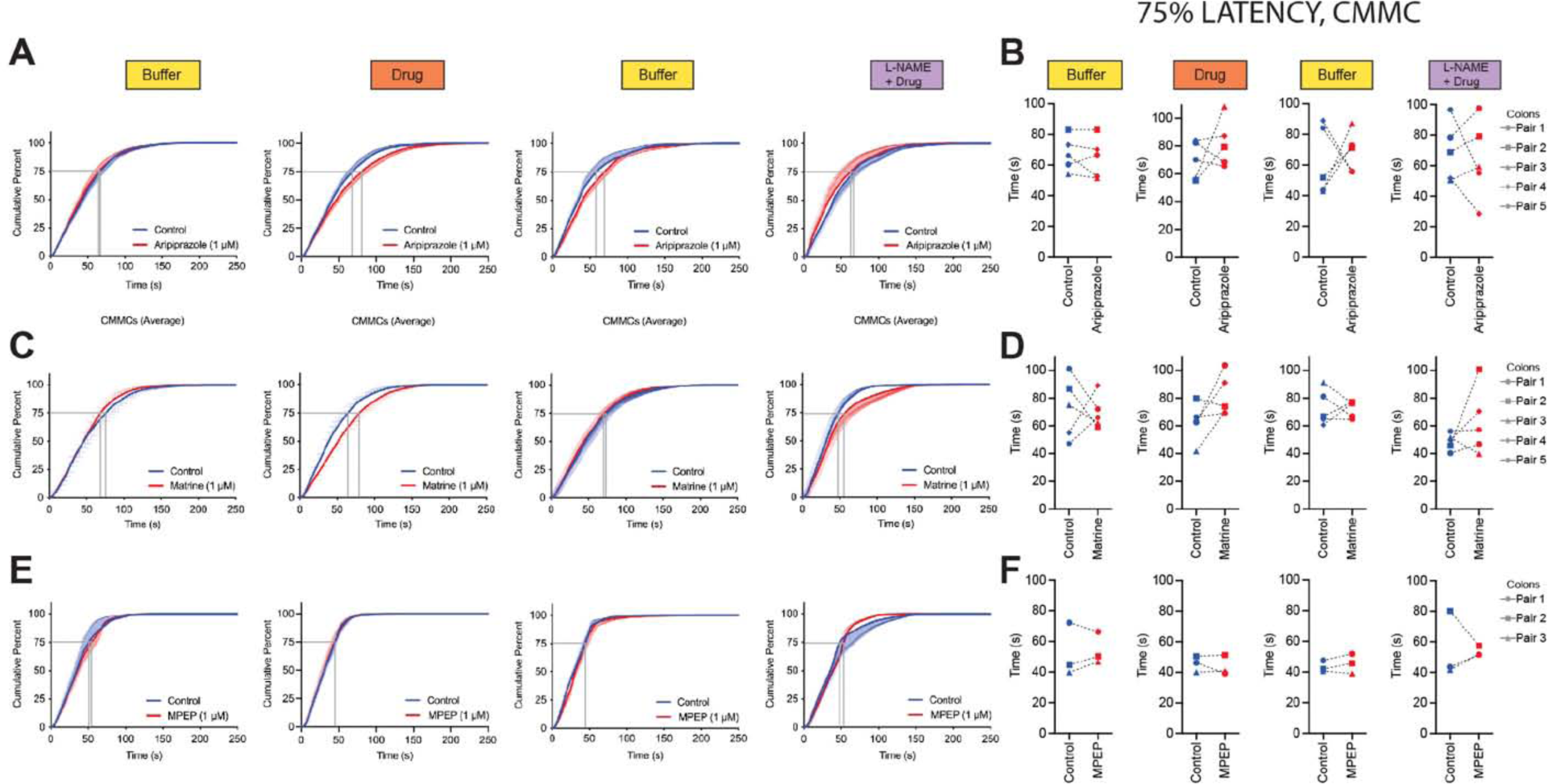
Effect of candidate drugs on *ex vivo* mouse colonic migrating motor complexes (CMMCs) A-F) Diagrams of CMMC cumulative percentile and quantifications of CMMC intervals at 75^th^ percentile for aripiprazole (A, B), matrine (C, D), and MPEP (E, F). Compounds were used at 1 μM and mean and one-directional SEM error bars are shown in panels **A**, **C** and **E**.

**Figure S23:**
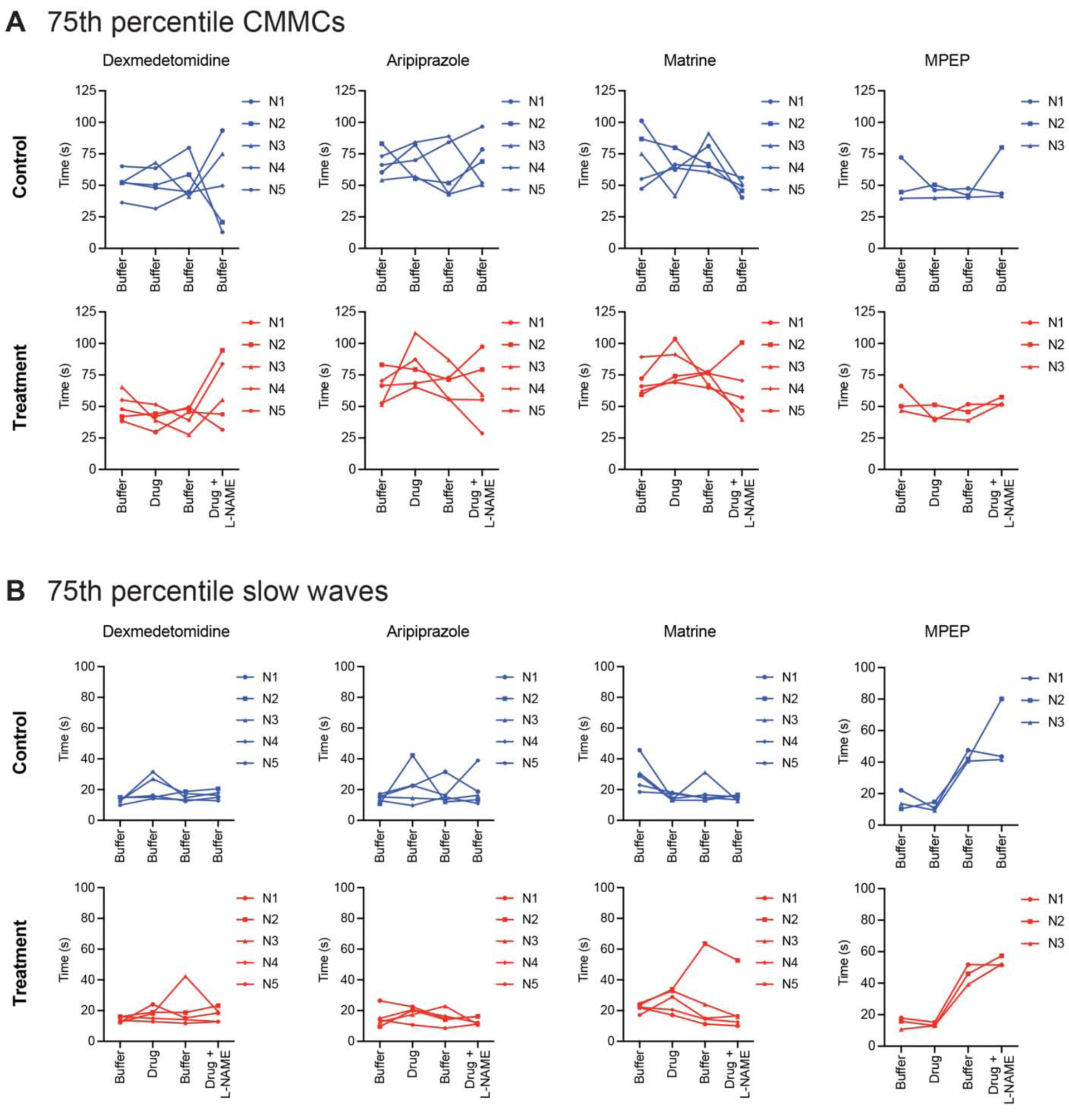
Effect of candidate drugs on *ex vivo* mouse colonic CMMC and slow wave intervals at 75^th^ percentile. **A and B**) Quantifications of CMMC (**A**) and slow wave (**B**) intervals (time difference between two consecutive contractions) at 75^th^ percentile for dexmedetomidine, aripiprazole, matrine, and MPEP. In each panel, top rows (blue) show control colons and bottom rows (red) show treatment colons within each pair. Compounds were used at 1 μM.

**Figure S24:**
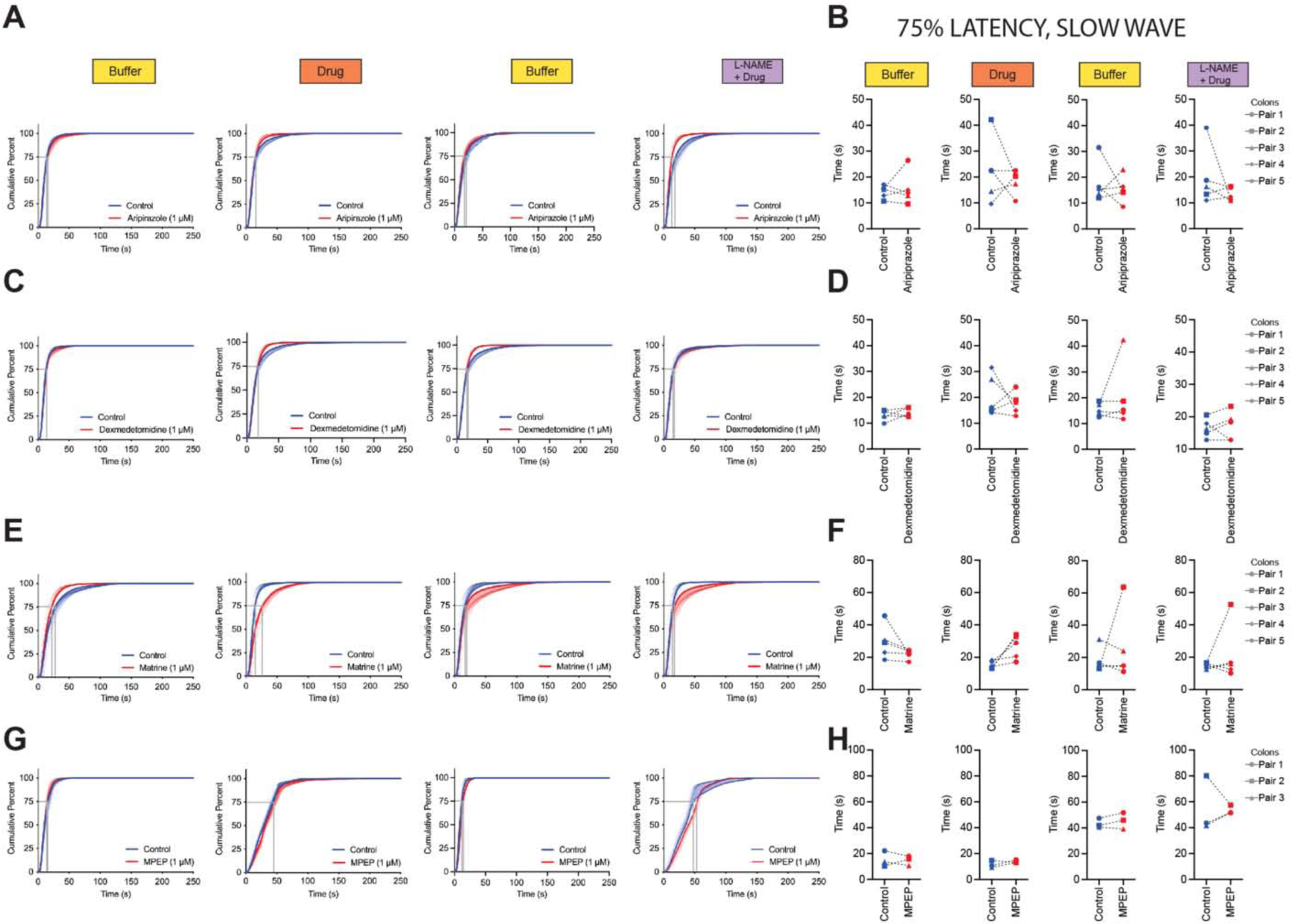
Effect of candidate drugs on *ex vivo* mouse colonic slow waves. **A-H**) Diagrams of slow wave cumulative percentile and quantifications of latency between slow waves at 75^th^ percentile for aripiprazole (**A, B**), dexmedetomidine (**C, D**), matrine (**E, F**), and MPEP (**G, H**). Compounds were used at 1 μM and mean and one-directional SEM error bars are shown in panels **A**, **C**, **E** and **G**.

**Figure S25:**
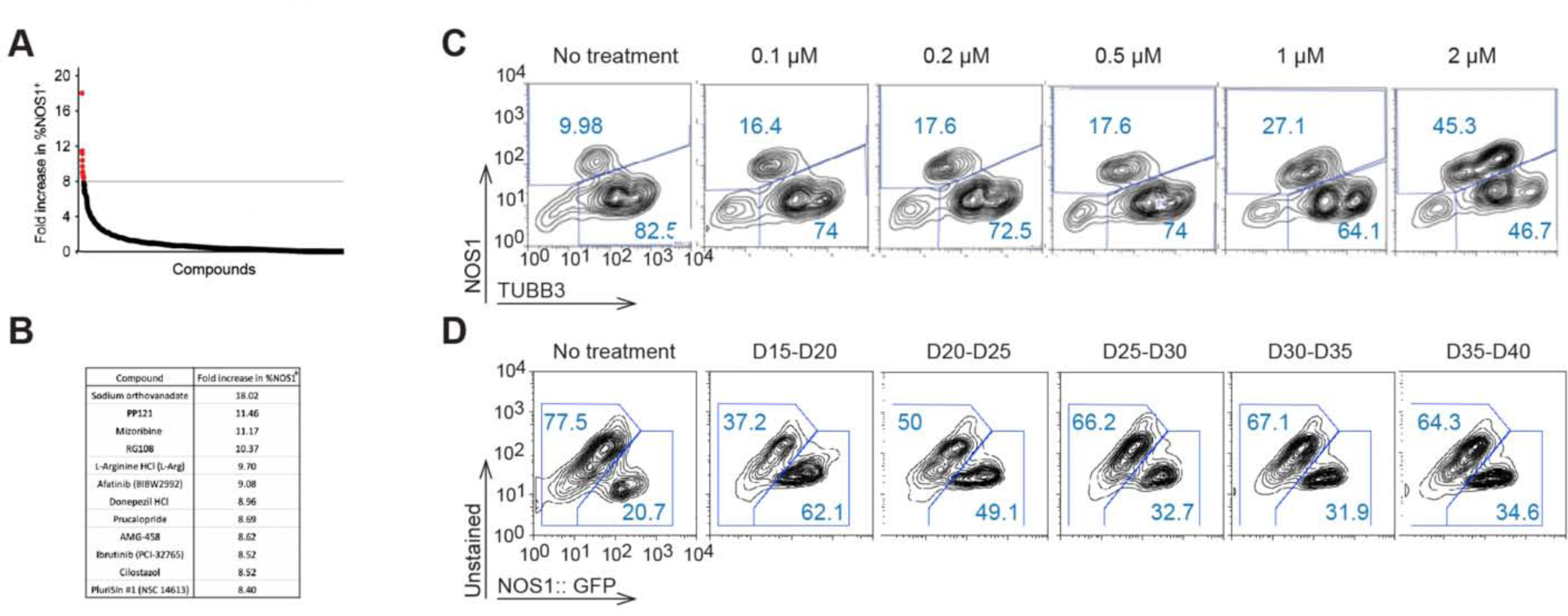
Small molecule high-throughput screening identifies compounds that enrich NO neurons in hESC-derived ENS cultures. **A)** Identifying candidate compounds that enrich NO neurons in stage 1 2D ENS cultures. **B)** HTS hits with a >8.0 fold increase in %NOS1**^+^** cells. **C and D**) PP121 induction of NOS1 expression is dose and time dependent. Representative flow diagrams showing NOS1 expression in stage 1 enteric neurons when cultures were treated with different PP121 concentrations at D15-D20 (**C**) or with 2 μM PP121 over different time-windows (**D**).

**Figure S26:**
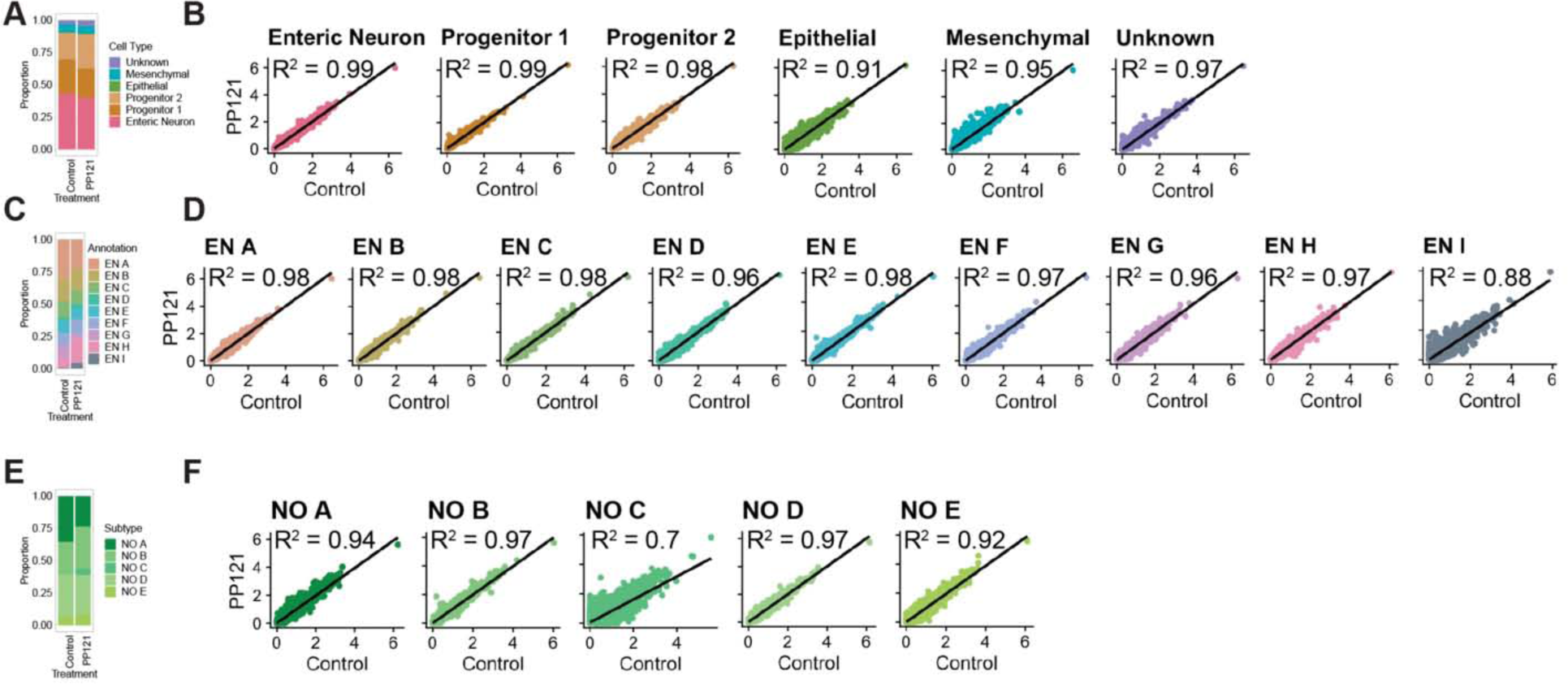
PP121 treatment enriches NO neurons without affecting their overall cellular diversity. **A)** Distribution of cell types in control versus PP121 treated stage 1 enteric ganglioid cultures. **B)** Correlation of average gene expression in matched control versus PP121 treated enteric ganglioid cell types. **C)** Distribution of neuronal subtypes in control versus PP121 treated stage 1 enteric ganglioid cultures. **D)** Correlation of average gene expression in matched control versus PP121 treated enteric ganglioid neuronal subtypes. **E)** Distribution of NO neuron subtypes in control versus PP121 treated stage 1 enteric ganglioid cultures. **F)** Correlation of average gene expression in matched control versus PP121 treated enteric ganglioid NO neuron subtypes.

**Figure S27:**
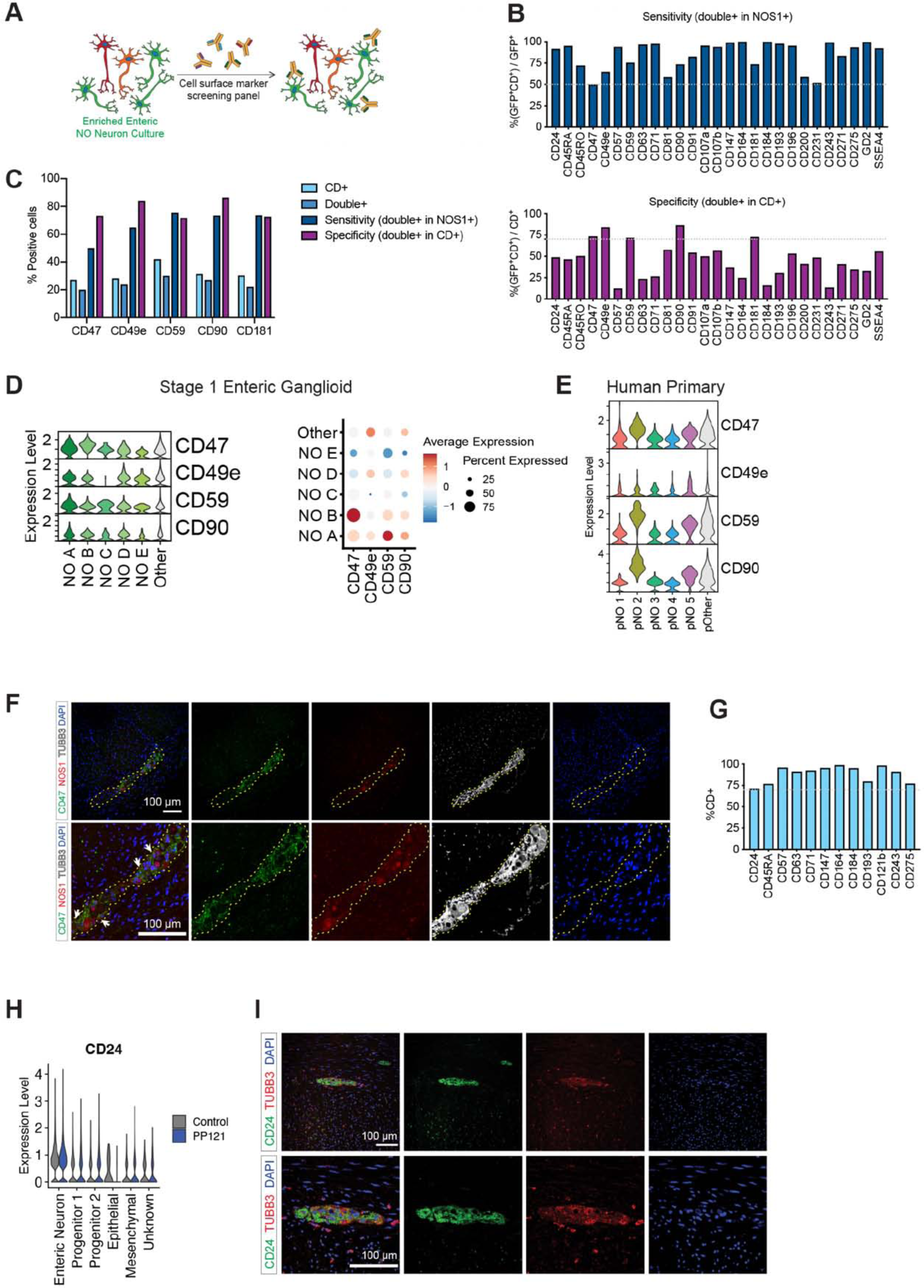
Human surface marker antibody screening identifies NO neuron specific surface markers. **A)** Schematic representation of using enriched enteric NO neuron cultures to identify their specific surface markers by a flow cytometry-based high-throughput antibody screening. **B)** Surface markers that were expressed in at least 50% of NOS1**^+^** cells, i.e. showing >50% sensitivity (top) were filtered based on their specificity for NOS1**^+^** cells (at least 70% of all stained cells being NOS1**^+^**, i.e. >70% specificity, middle). **C)** Surface marker screening top hits with the highest specificity as well as sensitivity for identifying enteric NO neurons. **D)** Violin plot stack showing the expression (left) and dot plot showing the average scaled expression (right) of surface marker screen hits in control and PP121 treated NO neuron subtypes versus other neurons. **E)** Violin plot stack showing the expression of surface marker screen hits in adult human NO neuron subtypes versus other neurons. **F)** Immunofluorescence analysis for expression of neuronal marker TUBB3, NOS1 and CD47 in adult human primary colon tissue. A myenteric plexus ganglion is shown. Arrows show colocalization of CD47 and NOS1. **G)** Surface markers that stained at least 70% of total cells as potential pan ENS cell markers. **H)** CD24 expression in stage 1 enteric ganglioid clusters in PP121 treated, and untreated samples. **I)** Representative immunofluorescence analysis of CD24 and neuronal marker TUBB3 expression in a human primary colonic section, showing a myenteric plexus ganglion.

**Figure S28:**
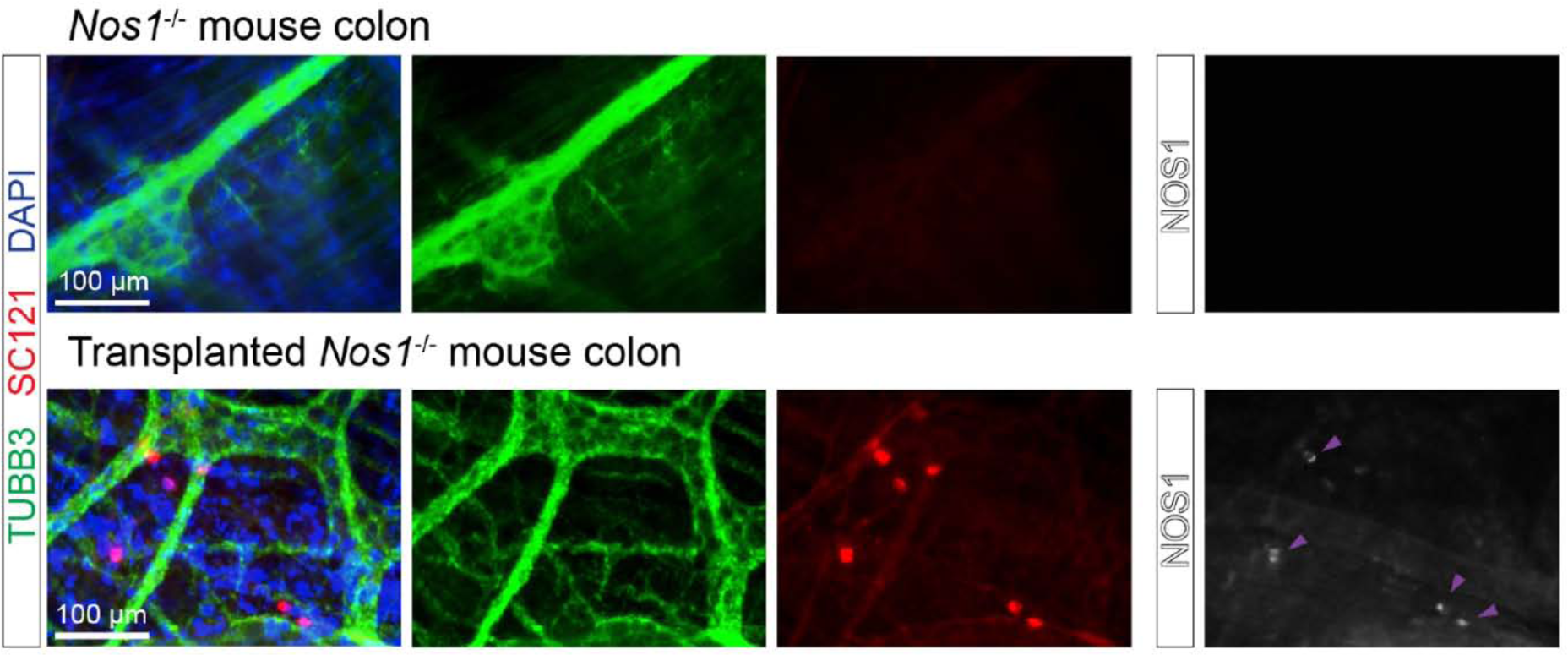
hESC-derived enteric ganglioids engraft in adult mouse colon. Immunohistochemical analysis of neuronal TUBB3, human cytoplasmic protein SC121, and NO neuron marker NOS1 in *Nos1*^-/-^ mouse colon (untreated with Cyclosporin A and non-operated, top) and 8 weeks post transplantation (bottom). Purple arrows point at NOS1^+^ cells.

## Supplementary videos

Video S1: Representative video of the effect of dexmedetomidine on mouse colonic motility *ex vivo*

Video S2: Representative video SC121^+^ human cells engraftment into the oral region of a *Nos1^-/-^* mouse colon 8 weeks post-surgery

Video S3: Representative video SC121^+^ human cells engraftment into the middle region of a *Nos1^-/-^* mouse colon 8 weeks post-surgery

Video S4: Representative video SC121^+^ human cells engraftment into the aboral region of a *Nos1^-/-^* mouse colon 8 weeks post-surgery

## Supplementary tables

Table S1: Neurochemical identity markers

Table S2: Neurochemical specific genes

Table S3: Antibodies

Table S4: Single cell and single nuclei transcriptomics quality control matrices

Table S5: Single cell and single nuclei transcriptomics clustering parameters

Table S6: Single cell and single nuclei transcriptomics cell type annotation genes

Table S7: Gene category lists for cell type specific molecular characterization

## Methods

### Culture and maintenance of undifferentiated human stem cells

Human embryonic stem cell (hESC) line H9 (WAe009-A, and reporter expressing derivatives hSYN::ChrR2-EYFP, NOS1::GFP) and induced pluripotent stem cell (hiPSC) line WTC-11 (UCSFi001-A) were plated on geltrex™-coated plates and maintained in chemically-defined medium (E8) as described previously (Barber et al., 2019). The maintenance cultures were tested for mycoplasma every 30 days.

### Enteric neural crest (ENC) induction

When the monolayer cultures of hPSCs reached about 70% confluency, a previously established 12-day enteric neural crest (ENC) induction protocol was initiated (Barber et al., 2019; Fattahi et al., 2016) (D0) by aspirating the maintenance medium (E8) and replacing it with neural crest induction medium A [BMP4 (1 ng ml^-1^), SB431542 (10 μM), and CHIR 99021 (600 nM) in Essential 6 medium]. Subsequently, on ENC induction days D2 and D4, neural crest induction medium B [SB431542 (10 μM) and CHIR 99021 (1.5 μM) in Essential 6 medium] and on D6, D8, and D10 medium C [medium B with retinoic acid (1 μM)] were fed to the cultures. Next, ENC crestospheres were formed during D12– D15 to facilitate the selection for ENC lineage and against contaminating ones in our cultures. In doing so, we removed ENC induction crest medium C on D12 and detached the ENC monolayers using accutase (30 min, 37 °C, 5% CO_2_). After centrifuging the samples at 290 x g for 1 min, we re-suspended the ENC cells in NC-C medium [FGF2 (10 ng ml^-1^), CHIR 99021 (3 μM), N2 supplement (10 μl ml^-1^), B27 supplement (20 μl ml^-1^), glutagro (10 μl ml^-1^), and MEM NEAAs (10 μl ml^-1^) in neurobasal medium] and transferred them to ultra-low-attachment plates to form free-floating 3D enteric crestospheres. On D14, when the free-floating enteric crestospheres could be observed, we gently gathered them in the center of each well using a swirling motion. Then, the old media was carefully aspirated from the circumference of each well without removing the crestospheres. After addition of the fresh NC-C medium, the cultures were incubated for 24 hours (37 °C and 5% CO_2_) prior to enteric neuron induction phase.

### Enteric neuron induction from enteric neural crests

On D15, enteric crestospheres were gathered in the center of the wells using a swirling motion and NC-C medium was removed using a P1000 micropipette in slow circular motion, avoiding the free-floating crestospheres. At this step protocol varied depending on the final desired culture layout (2D ENS cultures versus 3D enteric ganglioids). For 2D ENS cultures, after washing the enteric crestospheres with PBS, accutase (Stemcell Technologies, 07920) was added and plates were incubated for 30 minutes at 37 °C to dissociate the crestospheres. Then, remaining spheroids were broken by pipetting ENC medium [GDNF (10 ng ml^-1^), ascorbic acid (100 μM), N2 supplement (10 μl ml^-1^), B27 supplement (20 μl ml^-1^), glutagro (10 μl ml^-1^), and MEM NEAAs (10 μl ml^-1^) in neurobasal medium]. Cells were spun (2 min, 290 x g, 20-25 °C) and supernatant was removed. Pellet was resuspended in ENC medium and cells were plated on poly-L-ornithine (PO)/laminin/fibronectin (FN) plates at 100,000 viable cells per cm^2^. For 3D enteric ganglioids, we avoided accutase treatment and enteric crestospheres were fed with the same volume of ENC medium [GDNF (10 ng ml^-1^), ascorbic acid (100 μM), N2 supplement (10 μl ml^-1^), B27 supplement (20 μl ml^-1^), glutagro (10 μl ml^-1^), and MEM NEAAs (10 μl ml^-1^) in neurobasal medium]. Feeding continued every other day with ENC medium until D30-D40, after which, feeding frequency could be reduced to once or twice per week but with a larger volume of feeding medium.

### Immunofluorescence

For immunofluorescence (IF) staining, cells were initially fixed in 4% PFA in PBS (30 min, room temperature (RT), and then blocked and permeabilized by permeabilization buffer (PB) (Foxp3/Transcription Factor Staining Buffer Set, 00-5523) for another 30 minutes at RT. After fixation and permeabilization steps, cells were incubated in primary antibody solution overnight at 4 °C, and then washed three times with PB before their incubation with fluorophore-conjugated secondary antibodies at RT. Before imaging, stained cells were incubated with DAPI fluorescent nuclear stain and washed an additional three times. The list of antibodies and working dilutions is provided in Table S3.

### Preparation of enteric ganglioid frozen sections

hPSC-derived ganglioids were collected at stage 1 (day 37-50) and stage 2 (day 70-90), rinsed twice in PBS and fixed on ice in 4% PFA (SCBT sc-281692) for 3 hours, followed by replacing 90% of the supernatant with PBS for storage at 4 °C for up to 6 months. Ganglioids were treated with 5% sucrose (RPI Research Products 524060) in PBS for 10 minutes at room temp, followed by 10% sucrose in PBS for 2 hours at room temp and 20% sucrose at 4 °C overnight. Sucrose-treated ganglioids were positioned in cryomolds (Tissue-Tek® Cryomold® medium, VWR 25608-924), all 20% sucrose removed and incubated in 2:1 20% sucrose:OCT (Tissue Plus O.C.T. Compound Fisher HealthCare 5484) for 2 hours at room temperature before flash freezing in ethanol/dry ice. 12-20 μm sections were taken on a cryostat (Leica 3050S) adhered to Superfrost® Plus Micro Slide, Premium (VWR 48311-703) and dried on 42 °C slide dryer for up to 2 hours before storing at -80 °C for up to a year.

### Preparation of paraffin-embedded human colon sections

Human sigmoid colon tissue was received from the International Institute for the Advancement of Medicine (IIAM) that provides non-transplantable organs from Organ Procurement Organizations for biomedical research purposes. Colon tissue was obtained under sterile conditions, flushed with isotonic solution, submerged in organ transplant solution, and shipped on ice to laboratory within 24 hours post mortem. Full-thickness tissues pieces (∼2 cm2) were fixed overnight (<24 hours) in 10% neutral buffered formalin (Cancer Diagnostics, FX1003). Samples were transferred to 70% ethanol prior to paraffin embedding (Leica ASP6025, tissue processor). Approximately 5μM thick transverse tissue sections were cut onto coated glass slides (Superfrost® Plus Micro Slide; VWR, 48311-703) and air-dried overnight. All following slide preparation steps were performed at room temperature. Slides with paraffin sections were washed three times in clean xylene substitute (Sigma A5597), then once each in 100% ethanol, 95% ethanol, and 70% ethanol. Slides were then run under house DI water for 5 minutes before being placed in 1X PBS for storage at 4 °C for up to 4 weeks. Prior to staining, paraffin sections underwent antigen retrieval in either citrate buffer (Vector Laboratories Antigen Unmasking Solution H-3300) or TE buffer (Thermo 17890, brought to pH 9.0 with 1 M NaOH). Slides were incubated in buffer for 10 minutes at 95 °C using a Pelco BioWave Pro+ set to 400 watts.

### Staining enteric ganglioid frozen sections and paraffin-embedded human colon sections

Unless otherwise specified, all steps were performed at room temperature. Ganglioid frozen sections and paraffin-embedded human normal colon sections were prepared as above and then washed three times in PBS and blocked for 1-2 hours in serum (10% donkey or 10% goat) with 0.5% (v/v) Triton X-100 (VWR 0694). Slides were then incubated with primary antibody diluted in serum (10% donkey or 10% goat) with 0.1% Triton X-100 at 4 °C for 12-20 hours. Slides were washed six times for 20 minutes each in PBS with 0.1% Tween-20 (Sigma P1379) and incubated for 1 hour with Alexa Fluor conjugated secondary antibodies. The diluted secondary antibody solution was removed and replaced with 1.0 μg/mL DAPI in water for 10 minutes. The slides were washed six times for 20 minutes each in PBS with 0.1% Tween-20 and coverslips were mounted with Fluoromount-G (Southern Biotech 0100-01). The list of antibodies and working dilutions is provided in Table S3. Images were acquired on a Leica SP8 inverted confocal or on the Echo Revolve. For images that were stitched we used Leica’s LAS X tiling feature or the Grid/Pairwise stitching plugin for FIJI (PMID 19346324).

### 2-Photon fluorescence imaging

Imaging experiments were conducted on a custom-built upright 2-photon microscope operating with µManager software (San Francisco, CA). The excitation source was a 2-photon Coherent Chameleon Vision II laser operating at 760nm (Coherent, Santa Clara, CA). Images were collected using an Olympus LWD 1.05 NA water immersion objective (Olympus, Tokyo Japan). An emission filter collecting light between 380nm-420nm (Chroma, Bellow Falls VT) were used to image DAPI, while the fluorescence emission of Alexa 568 was collected using a filter between 565nm and 635nm (Chroma, Bellow Falls VT).

### Macro fluorescence imaging

Images were taken on a Nikon AZ100M “Macro” laser scanning confocal configured with long working distance low magnification lenses. The microscope is equipped with the standard 405 nm, 488 nm, 561nm, and 640 nm laser lines and has PMT detectors with a detection range from 400 - 700 nm. To reduce signal drop-off at the image edges we used an optical zoom factor of 2.1x and increased our lateral resolution using a digital zoom factor of 1.873x.

### Flow cytometry

For preparation of samples for flow cytometry analysis, cells were initially dissociated into single cell suspensions by accutase treatment (Stemcell Technologies, 07920, 30-60 min, 37 °C, 5% CO_2_) and then fixed and permeabilized using fixation/permeabilization buffers (Foxp3/Transcription Factor Staining Buffer Set, 00-5523). Cells were stained with primary and secondary antibodies as described above for immunofluorescence. Flow cytometry was conducted using a BD LSRFortessa cell analyzer and data were analyzed using Flowjo™ ( FlowJo™ Software Version *8.7*). The list of antibodies and working dilutions is provided in Table S3.

### Human synapsin::channelrhodopsin2-EYFP enteric ganglioids blue light activation

Enteric ganglioids were either exposed to blue light (100% laser intensity, 3 x 1-min exposure with 30 s intervals, EVOS FL) or left out in ambient light. Enteric ganglioids were then incubated for 45 minutes at 37 °C before dissociation, fixation and permeabilization for flow cytometry (see above). Cells were stained using antibodies against cFos (abcam, ab190289) and TUBB3 (Biolegend, 801202).

### Bulk RNA-seq data analysis

Total RNA was extracted using PureLink™ RNA Mini Kit. First strand cDNA was then synthesized with the Quantseq Forward Library preparation kit from Lexogen. Illumina compatible RNA sequencing libraries were prepared with Quantseq and pooled and sequenced on Illumina Hiseq 4000 platform at the UCSF Center for Advanced Technology. UMIs were extracted from the fastq files with umi_tools, and cutadapt was used to remove short and low-quality reads. The reads were aligned to human GENCODE v.34 reference genome using STAR aligner, and the duplicate reads were collapsed using umi_tools. Gene level counts were measured using HTSeq and compared using DESeq2.

### Single cell and single nuclei RNA sequencing sample preparation and data collection

All tubes and pipet tips used for cell harvesting were pre-treated with 1% BSA in 1X PBS. Cells were dissociated in Accutase (Stem Cell) at 37 °C, in 10 min increments, with end-to-end rotation, until single cell suspension was obtained. The cells were washed in Cell Staining Buffer (Biolegend) and stained with TotalSeq HTO antibodies for 30 min on ice. The cells were washed twice in Cell Staining Buffer and filtered through a 40 µm pipette tip strainer (BelArt). The cells were counted using Trypan Blue dye and hemocytometer and pooled for sequencing. scRNA-seq libraries were prepared with Chromium Next GEM Single Cell 3′ Kit v3.1 (10x Genomics), with custom amplification of TotalSeq HTO sequences (Biolegend). The libraries were sequenced on Illumina NovaSeq sequencer in the Center for Advanced Technologies (UCSF). The cell feature matrices were extracted using kallisto/bustools, and demultiplexed using seurat.

### Quality control and cell filtration

Datasets were analyzed in R v4.0.3 with Seurat v4 (Hao et al., 2021). The number of reads mapping to mitochondrial and ribosomal gene transcripts per cell were calculated using the “PercentageFeatureSet” function. Cells were identified as poor quality and subsequently removed independently for each dataset based on the number of unique features captured per cell, the number of UMI captured per cell and the percentage of reads mapping to mitochondrial transcripts per cell. Dataset specific quality control metric cutoffs can be found in Table S4.

### Dimensionality reduction, clustering and annotation

Where applicable, biological replicate samples were first merged using the base R “merge” function. Counts matrices were log normalized with a scaling factor of 10,000 and 2,000 variable features were identified using the “vst” method. For datasets specified in Table S5, count matrices of biological replicate samples were integrated using Seurat integration functions with default parameters. Cell cycle phase was predicted using the “CellCycleScoring” function with Seurat’s S and G2M features provided in “cc.genes.” The variable feature sets were scaled and centered, and the following variables were regressed out: nFeatures, nCounts, mitochondrial gene percentage, ribosomal gene percentage, S score and G2M score. Principal Components Analysis (PCA) was run using default settings and Uniform Manifold Approximation and Projection (UMAP) dimensionality reduction was performed using the PCA reduction. The shared nearest neighbors (SNN) graph was computed using default settings and cell clustering was performed using the default Louvain algorithm. Quality control metrics were visualized per cluster to identify and remove clusters of low-quality cells (less than average nFeatures or nCounts and higher than average mitochondrial and ribosomal gene percentage) (Table S5). The above pipeline was performed again on datasets after the removal of any low-quality cell clusters and for the sub-clustering analysis of the enteric neural crest, enteric neurons, nitrergic neurons and enteric glia. The number of principal components used for UMAP reduction and SNN calculation was determined by principal component standard deviation and varied for each dataset. The number of principal components used for SNN and UMAP calculation and the resolution used for clustering of each dataset can be found in Table S5. Cluster markers were found using the Wilcoxon Rank Sum test and clusters were annotated based on the expression of known cell type marker genes (Table S6). Following cell type annotation, gene dropout values were imputed using adaptively-thresholded low rank approximation (ALRA) (Linderman et al., 2018). The rank-k approximation was automatically chosen for each dataset and all other parameters were set as the default values. The imputed gene expression is shown in all plots and used in all downstream analysis unless otherwise specified.

### Analysis of Published Datasets

#### Quality control

Criteria used by the original authors of each dataset was used to identify and remove poor quality cells. Dataset specific quality control metric cutoffs can be found in Table S4.

#### Dimensionality Reduction and Clustering

Datasets were analyzed with Seurat using the methods and parameters described by the original authors. Morarach et al.: For all datasets, count matrices were normalized, mitochondrial gene percentage was regressed and 3000 variable features were returned using the “SCTransform” function. Highly expressed sex-specific and immediate early genes (Xist, Gm13305, Tsix, Eif253y, Ddx3y, Uty, Fos, Jun, Junb, Egr1) were removed form the variable feature list prior to running PCA. The dataset specific parameters used for the “RunUMAP”, “FindNeighbors” and “FindClusters” functions can be found in Table S5. Cell annotations determined by the authors were used for cell types and neuronal subtypes. Drokhlyansky et al.: For all datasets, count matrices were log normalized with a scaling factor of 10,000 and 2,000 variable features were identified using the “vst” method. Batch correction by “Unique_ID” was performed using mutual nearest neighbors correction (MNN) with the “RunFastMNN” Seurat Wrappers function. The dataset specific parameters used for the “RunUMAP”, “FindNeighbors” and “FindClusters” functions can be found in Table S5 Cell annotations determined by the authors were used for cell types and neuronal subtypes.

For consistency of comparison, gene dropout values were imputed using ALRA for all published datasets using automatically determined rank-k approximations and all other default values. The imputed gene expression is shown in all plots and used in all downstream analysis unless otherwise specified.

Glia Sub-clustering analysis. Glia were sub-clustered using methods similar to the original analysis pipeline described by each author above.

Morarach et al.: The E18 dataset contained a single transcriptionally homogenous glia cluster, so the glia and progenitor populations were sub-clustered together to provide comparative cell populations needed for downstream analysis. Subset datasets were then normalized, mitochondrial gene percentage was regressed and 3000 variable features were returned using the “SCTransform” function. Highly expressed sex-specific and immediate early genes (Xist, Gm13305, Tsix, Eif253y, Ddx3y, Uty, Fos, Jun, Junb, Egr1) were removed form the variable feature list prior to running PCA. The dataset specific parameters used for the “RunUMAP”, “FindNeighbors” and “FindClusters” functions can be found in Table S5.

Drokhlyansky et al.: Glia subset datasets were log normalized with a scaling factor of 10,000 and 2,000 variable features were identified using the “vst” method. Batch correction by “Unique_ID” was performed using mutual nearest neighbors correction (MNN) with the “RunFastMNN” Seurat Wrappers function. The dataset specific parameters used for the “RunUMAP”, “FindNeighbors” and “FindClusters” functions can be found in Table S5.

### Gene group expression characterization

Gene lists were compiled for genes belonging to ten different functional groups (transcription factors, neurotransmitter synthesis, neuropeptides, neurotransmitter receptors, neuropeptide receptors, cytokines, cytokine receptors, secreted signaling ligands, ligand receptors, and surface markers) (Table S7). For each dataset, the gene lists were filtered to remove low abundance genes (detected in less than 25% of cells of each cluster). Genes from these lists were determined to be exclusively expressed by a cluster if greater than 25% of cells of only a single cluster expressed the gene.

### Cell type transcriptional signature scoring

To find transcriptionally similar cell populations between two datasets, first the differentially expressed (DE) genes of the reference dataset are calculated from the non-imputed gene counts with the “FindAllMarkers” function using the Wilcoxon Rank Sum test and only genes with a positive fold change were returned. The DE gene lists are first filtered to remove genes not present in the query dataset. Then for each cell cluster in the reference dataset, a transcriptional signature gene list is made from the top 100 DE genes sorted by increasing adjusted p-value. The query dataset is then scored for the transcriptional signature gene lists of each reference dataset cell cluster using the “AddModuleScore” function based on the query dataset’s imputed gene counts.

### Spearman correlations

The transcriptional correlation of cell clusters in two datasets was calculated from the non-imputed gene counts and utilized Seurat’s integration functions to first find 3,000 anchor features based on the first 30 dimensions of the canonical correlation analysis and then integrate the two datasets using the same number of dimensions. The expression of these 3000 anchor features was then scaled and centered in the merged data object and the average scaled expression of each anchor feature was calculated for each dataset’s cell clusters of interest using the “AverageExpression” function. A Spearman correlation matrix comparing all cell clusters to all cell clusters was generated based on the average scaled expression of the 3000 anchor features.

### SWNE projections

The reference and query dataset counts matrices are first filtered to only include genes detected in both datasets. Similarly Weighted Nonnegative Embeddings (SWNE) are then generated for the reference dataset using the SWNE v0.6 package. First, nonnegative matrix factorization (NFM) generates component factors from the 3000 variable features calculated from the reference dataset non-imputed gene counts. Two dimensional component factor embeddings are calculated using sammon mapping and the cells and specified key genes are embedded in 2D relative to the component factors. Finally, a SNN network is calculated from the reference dataset and is used to smooth the cell positions. The query dataset is then mapped onto the reference dataset’s 2D component factor space by first projecting the query dataset onto the reference dataset’s NFM factors. The resulting query dataset cell embeddings are then smoothed by projection onto the reference dataset’s SNN network.

### Myenteric and submucosal Scoring

Patient metadata published by the authors was used to separately group neurons or glia by tissue layer origin. Pan-neuronal and pan-glial myenteric and submucosal gene signatures were created by performing the Wilcoxon Rank Sum test to identify DE genes between the myenteric and submucosal cell groupings. Neuronal and glial datasets were scored with the cell-type specific tissue layer signatures by first ordering the gene lists by increasing adjusted p-value and removing genes not detected in the dataset to be scored. The “AddModuleScore” function was then used to score the cells for the 100 most significantly enriched genes for each tissue layer.

### Neurochemical identification of neurons

The neurochemical identification of neurons was performed independently for each neurotransmitter to accommodate multi-neurochemical identities. For each neurotransmitter, a core set of genes were selected consisting of the rate-limiting synthesis enzyme(s), metabolism enzymes and transport proteins (Table S1). Cells were first scored for each neurotransmission associated gene set using the “AddModuleScore” function. A cell was then annotated as “x-ergic” if the cell’s expression of a rate limiting enzyme was greater than 0 and the cell’s module score for the corresponding gene set was greater than 0. A cell was annotated as “Other” if both criteria were not met. Multi-neurochemical identities were determined by concatenating the individually determined single neurochemical identities of each cell. The overall prevalence of each neurochemical identity per dataset was calculated by summing the total number of cells annotated for each single identity and calculating the percentage of each “x-ergic” identity from this sum total.

### Neurotransmitter response scoring

Separate gene lists were created containing all receptors activated by each neurotransmitter. Cells were scored for their expression of each neurotransmitter receptor family gene set using the “AddModuleScore” function.

### Glia GSEA hierarchical clustering

For each sub-clustered glia dataset, DE genes for each glial subtype were calculated using the “FindAllMarkers” function. Gene set enrichment analysis (GSEA) for the MSigDB gene ontology sets was performed on each glia subytpe’s upregulated DE genes (positive log2 fold change only) sorted by decreasing log2 fold change using fgsea v1.16. Normalized Enrichment Scores (NES) were calculated for gene sets containing a minimum of 15 genes in the DE gene list with the scoreType set to “positive”. Each glial subtype’s GSEA results were filtered to only include biological process gene sets but not filtered based on significance as to not limit the result to pathways enriched in the highest fold change genes. The NES of the filtered gsea results for all glial subtypes were then merged and pathways not detected in a glial subtype were assigned a NES of 0. Hierarchical clustering was then performed based on the NESs to cluster both the gene ontology pathways and the glial subtypes. After glia classes were determined by clustering, pathways enriched in each class were identified by filtering for pathways with an NES greater than 1.1 in all subtypes of a given class.

### PP121 vs control gene expression correlation

To compare the gene expression of control and PP121 treated cell types, neuronal subtypes and NO neuron subtypes, a subset dataset of each cell type and subtype annotation was first created. For each subset, the non-imputed average expression of all genes was then calculated for the control and PP121 treated cells using the “AverageExpression” function and natural log transformed for plotting. R^2^ values comparing the control and PP121 natural log expression values were calculated from linear modeling using the “y ∼ x” formula.

### cFOS expression screening

Stage 2 enteric ganglioids were dissociated using accutase and single cell suspensions (in ENC medium) were distributed in wells of V-bottom 96-well plates. Compounds from a neuronal signaling compound library (Selleckchem, USA) were added at 1 μM using a pin tool and cells were incubated for 75 minutes at 37 °C. Afterwards, cells were washed with PBS, and were immediately fixed for flow cytometry.

### NO release assay

For high-throughput measures of nitric oxide (NO) release, stage 1 2D ENS cultures (96-well plates) were used. After washing cells with Tyrode’s solution [NaCl (129 mM), KCl (5 mM), CaCl_2_ (2 mM), MgCl (1 mM), glucose (30 mM) and HEPES (25 mM) at pH 7.4], 70 μl/well of Tyrode’s solution was added to each well. Neuronal signaling compounds (Selleckchem, USA) were added at 1 μM using a pin tool. After a 45 minutes incubation at 37 °C, supernatants were used to determine NO release using an NO assay kit (Invitrogen, EMSNO). Briefly, the kit uses the enzyme nitrate reductase that converts nitrate to nitrite which is then detected as a colored azo dye absorbing light at 540 nm. NO release for each compound was presented as the A_540_ nm relative to the vehicle (DMSO).

### High-throughput screening to identify compounds that enrich NO neurons

Day 15 H9 hESC-derived enteric crestospheres were dissociated into single cells (accutase, Stemcell Technologies, 07920, 30 min, 37 °C), resuspended in ENC medium and were transferred into 384-well plates. Plates were incubated for 2 hours for cells to attach. Using a pin tool, drugs from a library of 1694 inhibitors (SelleckChem, USA) were added to wells at the final concentration of 1 μM and plates were incubated with drugs until D20, when media were changed to ENC with no drugs. At day 40, cells were fixed, stained for NOS1 and imaged using InCellAnalyzer 2000 (GE Healthcare, USA). Hits were selected based on the fold increase of the percentage of NOS1^+^ cells relative to the wells treated with vehicle (DMSO).

### Surface marker screening

For human surface marker screening, PP121-treated NOS1::GFP enteric ganglioids from four independent differentiations were pooled, dissociated into single cells (accutase, Stemcell Technologies, 07920, 30-60 min, 37 °C, 5% CO_2_) and fixed (Foxp3/Transcription Factor Staining Buffer Set, 00-5523, 30 min, 4 °C). Cells were permeabilized and blocked (same staining kit) prior to incubation with anti GFP antibody (abcam, ab13970, 4 °C).

After three washes, cells were stained with Alexa Fluor 488-conjugated secondary antibody (40 min, RT). Secondary antibody solution was removed (3 x washes) and cells were incubated with a blocking buffer containing PBS and 2% FBS (30 min, on ice). Cells were divided in a 240:16 ratio corresponding to the number of library antibodies raised in mouse and rat, and received anti-mouse and anti-rat Alexa Fluor 647-conjugated secondary antibodies respectively. Then, they were distributed into V-bottom 96-well plates and treated with library antibodies for 30 minutes on ice (BD Biosciences, 560747). After two washes, surface marker and GFP signals were quantified by high-throughput flow cytometry (BD LSRFortessa). NO neuron specific surface markers were identified based on the highest sensitivity (highest percentage of CD^+^GFP^+^ cells) and highest specificity (lowest percentage of CD^+^GFP^-^ cells).

### Drug target interaction prediction

We obtained canonical SMILES of our hits from PubChem <u>(De Giorgio et al., 2016; Niesler et al., 2021)</u> and generated a list of their known and predicted targets by combining data from the following databases: BindingDB (https://www.bindingdb.org/), Carlsbad (http://carlsbad.health.unm.edu/), DINIES (https://www.genome.jp/tools/dinies/), PubChem BioAssay (https://pubchem.ncbi.nlm.nih.gov/, filtered for active interactions), SEA (http://sea.bkslab.org/, filtered for MaxTC >0.4), SuperDRUG2 (http://cheminfo.charite.de/superdrug2/) and SwissTargetPrediction (http://www.swisstargetprediction.ch/).

### *In vivo* cell transplantation

Specified pathogen free (SPF) homozygote neuronal nitric oxide synthase knockout mice (B6.129S4-*Nos1^tm1Plh^*/J; *nNos1^-/-^*) were bred and maintained, in individually ventilated cages (IVC), for use as recipients. Animals used for these studies were maintained, and the experiments performed, in accordance with the UK Animals (Scientific Procedures) Act 1986 and approved by the University College London Biological Services Ethical Review Process. Animal husbandry at UCL Biological Services was in accordance with the UK Home Office Certificate of Designation. As *Nos1^-/-^* mice are immunocompetent, cyclosporin A (250 μg/ml in drinking water) was administered orally two days prior to transplantation to reduce the possible rejection of donor human cells. Cyclosporin A*-* treated *Nos1^-/-^* mice were chosen at random, from within littermate groups, and stage 1 enteric ganglioids were transplanted into the of P23-P27 mice, via laparotomy under isoflurane anesthetic. Briefly, the distal colon was exposed and enteric ganglioids, containing 0.5-1 M cells were subsequently transplanted to the serosal surface of the distal colon, by mouth pipette, using a pulled glass micropipette. Each transplanted tissue typically received 3 ganglioids which were manipulated on the surface of the distal colon, with the bevel of a 30G needle, to ensure appropriate positioning. Transplanted *Nos1^-/-^* mice were maintained with continued free access to cyclosporin A (250 μg/ml) treated drinking water for up to 8 weeks post-transplantation, to ensure extended immunosuppression, before sacrifice and removal of the colon for analysis. As cyclosporin A can affect several signaling pathways and induce gene expression changes, it is crucial to verify immunofluorescence results using appropriate controls such as tissue from cyclosporin A treated untransplanted animals in follow up studies. In addition, other immunocompromised backgrounds (e.g. NSG) will be important to further verify these engraftment results.

### Tissue preparation and fixation

Following the excision, the entire colon was pinned in a Sylgard (Dow, MI, USA) lined petri dish and opened along the mesenteric border. The mucosa was subsequently removed by sharp dissection and tissues were fixed in 4% PFA in PBS (45 min-1 hour, 22 °C) for further processing.

### Tissue staining

Colonic longitudinal muscle myenteric plexus (LMMP) tissues were fixed with 4% PFA (1 h on ice), Thermo scientific, J19943-K2) and blocked and permeabilized with a buffer containing 1% BSA and 1% triton X-100 (in PBS, 45 min, RT). Then, tissues were incubated with primary antibody solutions (in the same buffer, overnight, 4 °C) and were washed three times before treatment with fluorophore-conjugated secondary antibodies (1 h, RT). Samples were stained with DAPI and washed prior to mounting using vectashield (Vector Laboratories, H-1400). Antibodies are listed in Table S3.

### Multi-Electrode Array (MEA) analysis

#### Data acquisition

Neuron activity was recorded with the Axion Maestro Edge on Cytoview MEA 24-well plates in 1-hour recording sessions for each condition. Neuormodulator or vehicle were added by removing the plate from the Maestro Edge, half-changing the media with 2x concentrated neuromodulator or vehicle in pre-warmed media, and immediately returning the plate to the Axion to resume recording. Optogenetic stimulation was performed with the Axion Lumos attachment, stimulating all wells of the plate with 488 nm light at 50% intensity, 1 second on, 4 seconds off, 30 times.

#### Data processing

Raw data were first spike sorted with a modified version of SpikeInterface (https://github.com/SpikeInterface) using MountainSort to identify high quality units by manually scoring based on amplitude, waveform shape, firing rate, and inter-spike interval contamination. For pharmacology experiments, neurons were matched between vehicle and neuromodulator recordings by examining all detected units on a specific electrode after spike scoring and identifying units with identical waveforms. Firing rates of these “paired” units from all wells that received the treatment were compared across the control and neuromodulator conditions. Positive responders were units that had a firing rate change greater than +0.1 Hz; negative responders had a firing rate change less than -0.1 Hz; neutral responders had a firing rate change between -0.1 to +0.1 Hz. For optogenetic experiments, individual units were again extracted with SpikeInterface and manually scored. Recordings were separated into “on” times when the LED was active and “off” times when it was not. All units were compiled and firing rates for each unit were compared during the on and off windows.

### *Ex vivo* colonic motility assays

#### Preparation of solutions

Krebs buffer [NaCl (117 mM), KCl (4.7 mM), NaH_2_PO_4_ (1.2 mM), MgCl_2_ (1.5 mM), CaCl_2_.2H_2_O (2.5 mM), NaHCO_3_ (25 mM), Glucose (11 mM), pH 7.4] was placed in a 37°C water bath and aerated with 95% O_2_ and 5% CO_2_ (carbogen) gas mixture for at least 30 minutes prior to experiment onset. “Drug” treatment solutions were freshly prepared by adding the drug compound into Krebs buffer before starting data acquisition. The solution with NOS1 inhibitor was prepared by adding N omega-nitro-L-arginine methyl ester hydrochloride (L-NAME) to the drug solutions making “Drug+L-NAME”.

#### Tissue dissection

For each experimental replicate, a pair of 8-week-old wild type C57BL6 mice (male) were placed in a sealed chamber and euthanized using CO_2_ asphyxiation followed by cervical dislocation. The lower GI tract (cecum and colon) was removed and immediately transferred to 37 °C carbogenated Krebs buffer, with the fecal matter still inside. Adipose tissue and mesentery were removed before placing the colons in the organ bath reservoir of gastrointestinal motility monitor (GIMM) apparatus. GIMM had two reservoirs making simultaneous acquisition of control, and drug-treated colons possible.

#### Experimental set-up and procedure

GIMM was designed based on a previously reported model (Swaminathan et al., 2016). The organ reservoir of GIMM has two-chambers for recording two specimens simultaneously. It is connected to working solutions kept at 37 °C via a 4-channel peristaltic pump (WPI, PERIPRO-4LS). Lower GI tract was harvested and transferred to the organ bath with the Krebs buffer was flowing through. The cecum was pinned down at the proximal tip and the distal end of the colon was pinned through the serosa/mesentery. Five 10-min (for the first experiments) or sequential 6-min (for the sequential drug treatment in the presence and absence of L-NAME) videos were recorded using the IC capture software (Imaging Source) with a high resolution monochromatic firewire industrial camera (Imaging Source^®^, DMK41AF02) connected to a 2/3’ 16 mm f/1.4 C-Mount Fixed Focal Lens (Fujinon HF16SA1). While tissue in the control chamber was only exposed to Krebs solution, the order of solutions in the experimental chamber was: Krebs, drug compound, Krebs (6 min each), L-NAME (2 min), L-NAME in the presence of a drug compound (6 min) and Krebs (6 min). The chambers were cleaned after each acquisition.

#### Data and statistical analysis

VolumetryG9a was used to generate the spatiotemporal map (STM) of each acquisition (Spear et al., 2018). Slow waves (SW) and colonic migrating motor complexes (CMMC) data were generated from STMs. Statistical analyses were performed using PRISM.

### Generating figure schematics

We used Adobe Illustrator (version 25.4.1) to generate schematics for the figures.

